# MTALTND4, a second protein coded by *nd4* impacts mitochondrial bioenergetics

**DOI:** 10.1101/2022.04.28.489924

**Authors:** Laura Kienzle, Stefano Bettinazzi, Marie Brunet, Thierry Choquette, Hajar Hosseini Khorami, Xavier Roucou, Christian R Landry, Annie Angers, Sophie Breton

## Abstract

Recent evidence suggests that the coding potential of the mitogenome is underestimated. We found a downstream alternative ATG initiation codon in the +3 reading frame of the human mitochondrial *nd4* gene. This newly characterized alternative open reading frame (altORF) encodes a 99-amino acids long polypeptide, MTALTND4, which is conserved in primates. This small protein is localized in mitochondria and cytoplasm and is also found in the plasma, and it impacts mitochondrial physiology. Alternative mitochondrial peptides such as MTALTND4 may offer a new framework for the investigation of mitochondrial functions and diseases.

## MAIN

The recent identification in bacterial and eukaryotic genomes (including human) of thousands of unannotated yet translated small open reading frames (smORFs) and alternative open reading frames in reference genes (altORFs) has brought to light a whole new class of important genes (*1-4*). Although the elucidation of their function remains challenging, peptides derived from smORFs and altORFs reveal a new facet of eukaryote and bacterial proteomes, enabling a better understanding of key evolutionary processes such as the birth of novel genes and proteins, and providing novel fundamental biological knowledge as well as new avenues for therapeutic discovery. Mitochondria-derived proteomes do not escape this reality, given that the proteomes of their bacterial ancestors and ‘modern host cells’ are richer than previously anticipated (*1-4*). Up to now, 16-38 amino acid long micropeptides of functional importance have been shown to be encoded in the human mitochondrial 16S and 12S rRNA genes (*5-9*). These small proteins, called Humanin, SHLPs (small humanin-like peptides 1-6) and MOTS-c appear to modulate mitochondrial and cellular biology (*8*), as well as global physiology, acting as signaling molecules secreted from cells and found in circulation (plasma) (*5, 6*). However, whether altORFs and smORFs exist in mitochondrial protein-coding genes or elsewhere in the human mitogenome has not been addressed yet.

A preliminary *in silico* examination of the unresolved protein-coding potential of the human mtDNA has revealed 227 unannotated ORFs of at least 60 nucleotides on either strand (Fig. 1). To identify potentially novel mtDNA-encoded proteins, unannotated ORFs (Fig. 1 and table S1) were selected for further investigation using three different approaches: (i) we searched among the 227 unannotated ORFs for those having a minimal conserved Kozak sequence known to be favorable for efficient translation initiation (*5, 10*) (table S2), (ii) we interrogated the OpenProt database (table S3), which enforces a polycistronic model of eukaryotic genome annotations (*11*), and (iii) we interrogated mitochondria-derived mass spectrometry datasets to see if peptides derived from our 227 unannotated ORFs could be detected (table S4 and fig. S1). These three approaches respectively returned 13, 14 and 36 candidates (two OpenProt candidates were found also in the MS dataset) and we respectively chose one, 4 and 4 of these candidates for antibody production (Fig. 1 and tables S1-S4). Immunoblotting was used to assess their expression in human HeLa and HEK-293T cells.

**Figure 1.**
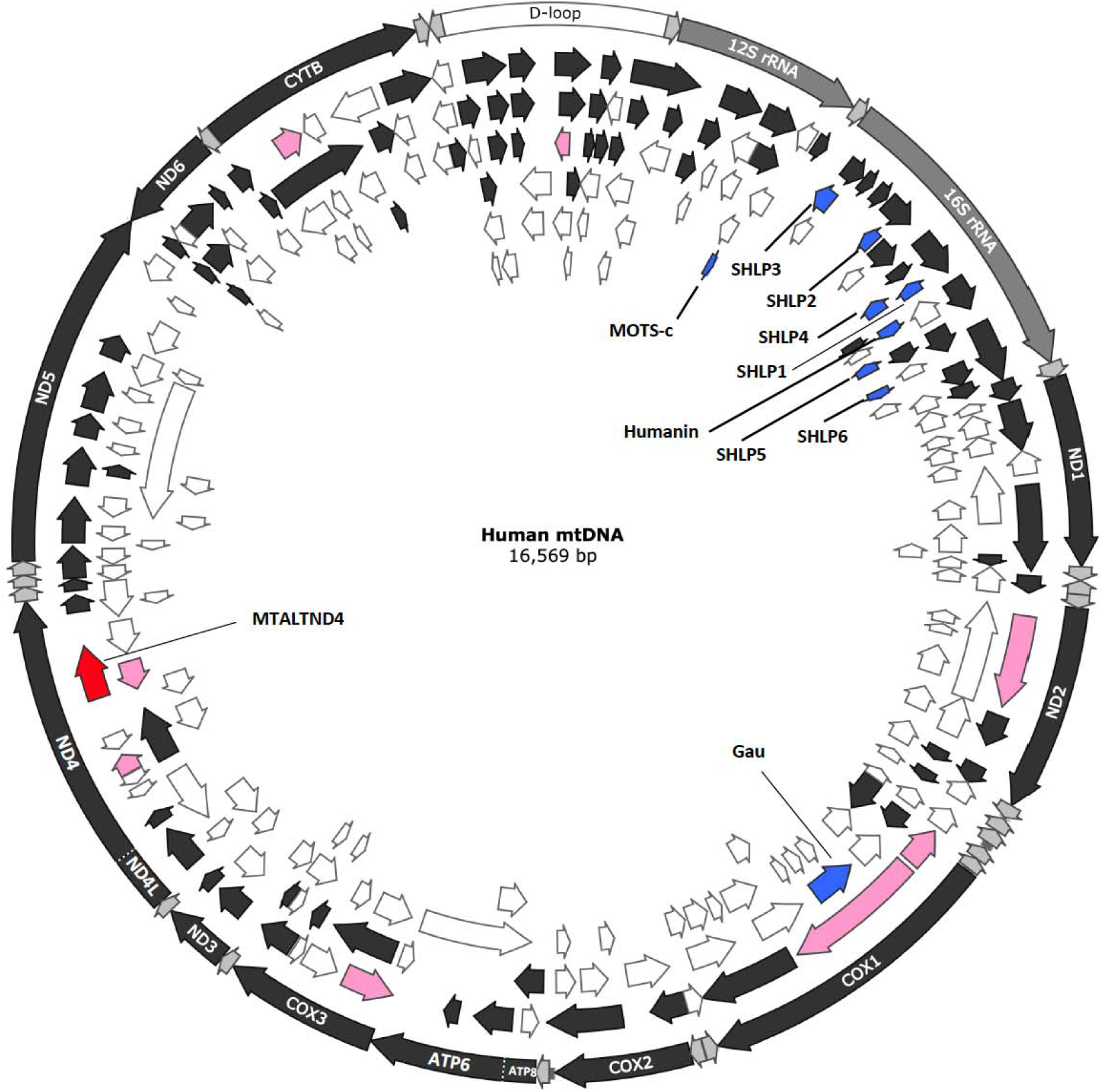
Small ORFs and alternative ORFs in the human mitochondrial genome. In dark gray in the external circle: the typical 13 protein-coding genes. In gray in the external circle: the two rRNA genes. In white in the external circle: the d-loop. Arrows positioned inside the circle indicate smORFs and altORFs on the main coding 5’-3’ (frames 1, 2 and 3: black arrows) and complementary 3’-5’ (frames -1, -2 and -3: white arrows) strands. In blue: the ORFs coding for the micropeptides humanin, SHLP1-6 and MOTs-c found in the 16S and 12S rRNA genes. The putative gau gene is also shown. In pink: the eight chosen candidates for antibody production. In red: MTALTND4.

Of these nine candidates, one unannotated ORF-derived peptide, which was identified by mass spectrometry (approach 3; see tables S1 and S4), was detected by western blotting and its mitochondrial origin was validated with the loss of western blot signal in cells devoid of endogenous mtDNA (Rho0) and in cells treated with chloramphenicol that inhibits mitochondrial protein synthesis, limiting the possibility of a nuclear origin due to a nuclear mtDNA transfer [NUMT; (*14*)] (fig. S2). This previously unannotated ORF-derived peptide is referred to as MTALTND4 (Mitochondrial Alternative ND4 protein) because the ORF sequence is found within the mitochondrial *nd4* gene in an alternative reading frame (Fig. 1 and Fig. 2*A-E*). The *mtaltnd4* altORF is 300 nucleotides long and code for a putative 99 amino-acid peptide (with no predicted transmembrane helix) using the mitochondrial vertebrate genetic code and considering that human mitochondria use only UAA and UAG as stop codons as recently suggested (*12, 13*) (Fig. 2*A* and table S1; see supplementary text 1). Multiple peptide sequence alignments indicate that MTALTND4 is well conserved in primates in general (Fig. 2*B* and fig. S3; see supplementary text 2). Although somewhat higher than expected, a specific 25 kDa western blot signal was observed in both HeLa and HEK-293T cells for endogenous MTALTND4 (Fig. 2*C*). Our results showed that endogenous MTALTND4 is localized in the cytoplasm where it is co-localized to mitochondria (Fig. 2*F*), a pattern that is similar also for Humanin, MOTs-c and SHLPs (*5-7*), and the absence of signal in HeLa-chloramphenicol treated cells confirmed again the mitochondrial origin of the alternative peptide (fig. S4). Our custom antibody, but not the pre-immune serum, was able to immunoprecipitate MTALTND4 from HeLa cell lysates, confirming the existence of an endogenous MTALTND4 peptide (Table S5).

**Figure 2.**
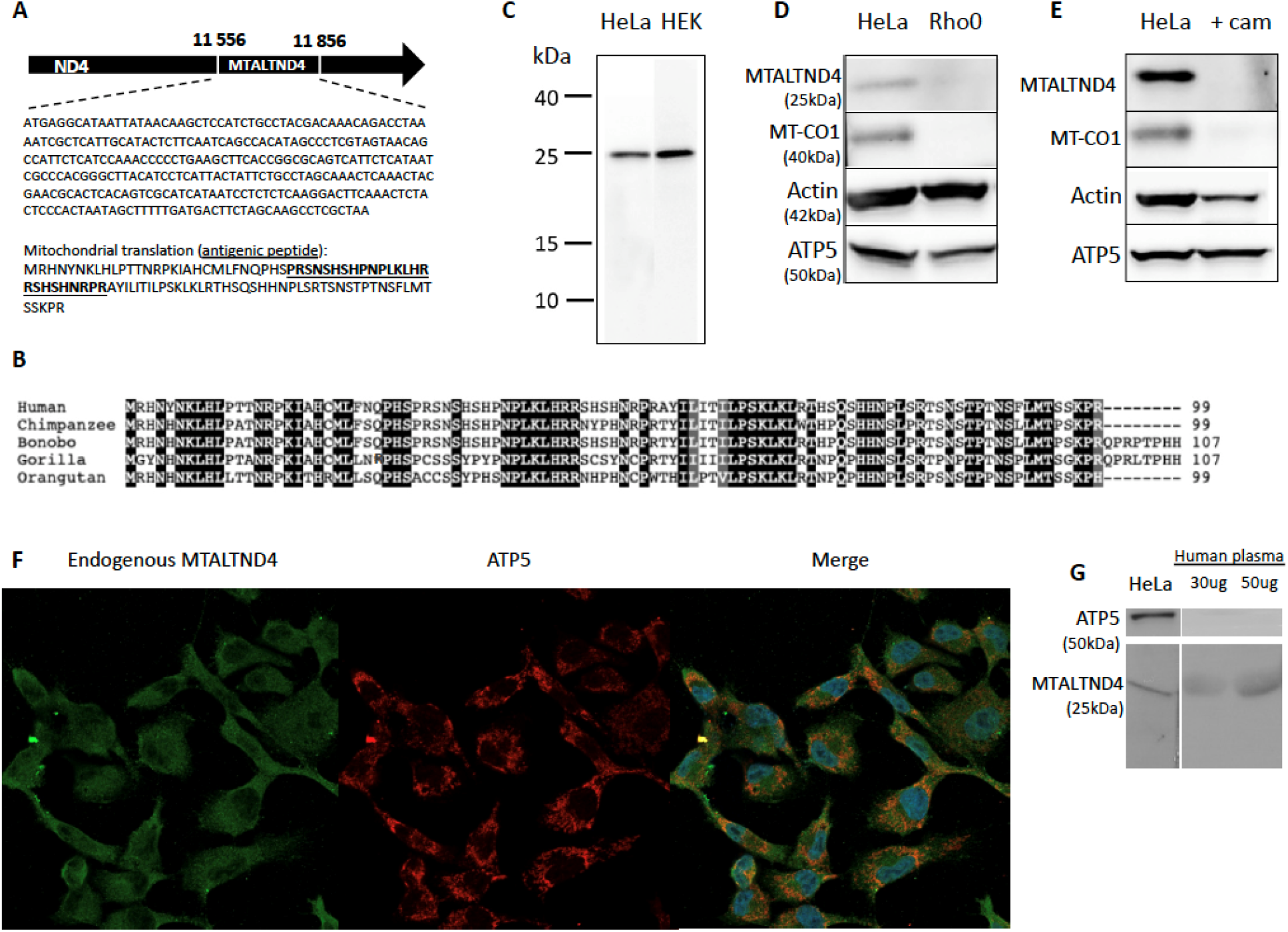
Identification of an alternative protein in the mtDNA-encoded *nd4* gene. *(A)* The MTALTND4 ORF and its DNA and protein sequences (antigenic sequence uderlined). This ORF is frameshifted by 2 nucleotides relative to ND4 reading frame. *(B)* Multiple peptide sequence alignment in five primate species. Black background for residues conserved in all species and grey for residues conserved in 4 out of 5 species. *(C)* MTALTND4 detected by western blotting in HeLa and HEK-293T cells. *(D)* MTALTND4, mtDNA-encoded cytochrome c oxidase subunit I (CO1), nuclear-encoded Actin and nuclear-encoded mitochondrial ATP5 in HeLa (control) and HeLa Rho0 cells, which are devoid of mitochondrial DNA. *(E)* MTALTND4, mtDNA-encoded cytochrome c oxidase subunit I (CO1), nuclear-encoded Actin and nuclear-encoded mitochondrial ATP5 in HeLa (control) and HeLa cells treated with chloramphenicol, which inhibits mitochondrial translation. *(F)* Detection of MTALTND4 and nuclear-encoded mitochondrial ATP5 by immunofluorescence in HeLa cells. *(G)* Detection of MTALTND4 and nuclear-encoded mitochondrial ATP5 by western blotting in human plasma.

Similar to Humanin, MOTS-c and SHLPs (*5-7*), the presence of MTALTND4 in the plasma was confirmed by western blot (Fig. 2*G*), which was not the case for the mitochondrial protein ATP5 (encoded by the nuclear DNA and also devoid of predicted TMH). Following its detection in the plasma, we hypothesized that MTALTND4, as other mitochondria-derived micropeptides (*6, 7*), could act as a signaling molecule secreted from cells in the circulation and regulate cell physiology. We thus examined its exogenous effect on cellular metabolism using HeLa and HEK-293T cells treated with the complete synthetic peptide at different concentrations. Overall, we observed a strong, immediate effect of MTALTND4 on mitochondrial respiration. Specifically, the presence of the peptide decreased oxygen consumption in both HeLa and HEK-293T cells. The effect appeared to be dose-dependent, although the concentration threshold differed between cell lines. For example, MTALTND4 at a concentration of 5 µM was sufficient to impact the basal oxygen consumption rate (i.e. the ROUTINE respiration) of HeLa cells, whereas up to 30 µM was necessary in the case of HEK-293T cells (Fig. 3*A,B*; supplementary table S6). Such effect was not observed using the shorter antigenic peptide as a control. It is worth mentioning that a concentration of 30 µM did not negatively impact HeLa or HEK cell proliferation or viability after 24h treatment (see below).

**Figure 3.**
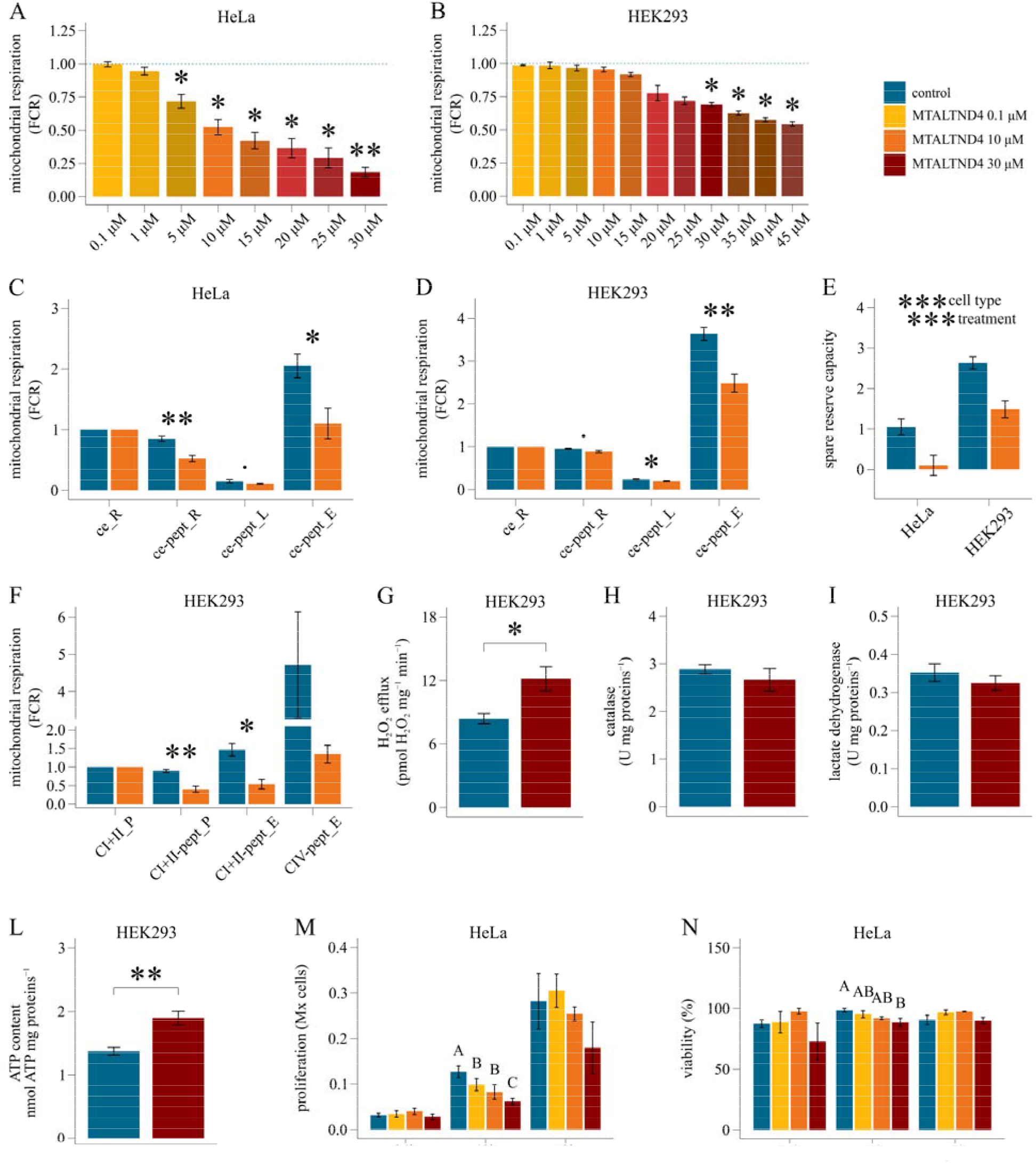
Impact of MTALTND4 upon mitochondrial and cell physiology. *(A-B)* Dose-dependent effect on intact (ce) HeLa and HEK-293T cells routine respiration (*n*=3 and *n*=4, respectively). Respirometry data were normalized for the routine respiration in absence of the peptide. For each titration point, data are represented as fraction of their own simultaneous control (blue dotted line). *(C-D)* Mitochondrial respiration in intact (ce) HeLa and HEK-293T cells in presence of 10 µM MTALTND4 (*n*=6-9 and *n*=6, respectively). Respirometry data were expressed as flux control ratios (FCR), normalized for the routine respiration before treatment (*‘ce_R’*). *‘ce-pept_R’*: routine respiration with MTALTND4; *‘ce-pept_L’*: leak respiration with MTALTND4; *‘ce-pept_E’*: maximal uncoupled respiration with MTALTND4. *(E)* Mitochondrial spare reserve capacity (SRC) in HeLa and HEK-293T cells in presence of 10 µM MTALTND4 (*n*=6 and *n*=6, respectively). *(F)* Mitochondrial respiration in permeabilized (pce) HEK-293T cells in presence of 10 µM MTALTND4 (*n*=6). Respirometry data were normalized for the CI+II-sustained coupled respiration before treatment (*‘CI+II_P’*). *‘CI+II-pept_P’*: CI+II-linked coupled respiration with MTALTND4; *‘CI+II-pept_E’*: CI+II-linked uncoupled respiration with MTALTND4; *‘CIV-pept_E’*: Cytochrome *c* oxidase standalone capacity with MTALTND4. *(G)* Hydrogen peroxide efflux rate (pmol H_2_O_2_ · mg proteins^−1^ · minute^−1^) in intact HEK-293T cells in presence of 30 µM MTALTND4 (*n*=5). *(H)* Catalase activity (U · mg proteins^−1^) in intact HEK-293T cells incubated 4 hours with 30 µM MTALTND4 (*n*=5). *(I)* Lactate dehydrogenase activity (U · mg proteins^−1^) in intact HEK-293T cells incubated 4 hours with 30 µM MTALTND4 (*n*=5). *(L)* ATP content (nmol ATP · mg proteins^−1^) in intact HEK-293T cells incubated 4 hours with 30 µM MTALTND4 (*n*=5). *(M-N)* Proliferation (Mx cells) and viability (%) of HeLa cells measured at different time points (24, 48 and 72 hours) and different MTALTND4 concentrations (0, 0.1, 10 and 30 µM) (*n*=3-5). Statistical analyses: *(A-B)* one sample *t* test; *(C, D, F, G, H, I, L)* paired *t* test; *(E)* linear mixed model. Factors *‘cell type’* (2 levels) and *‘treatment’(2 levels)* plus interaction. *(M, N)* linear mixed model. Factor ‘*treatment’* (4 levels), with letters indicating statistical difference following a *post hoc* multi comparison test. Data shown as mean ± sem. **·**0.05 > *p* ≤0.09; **\****p* ≤0.05; **\*\****p* ≤0.01; **\*\*\****p* ≤0.001. A detailed summary is reported in supplementary tables S6-S10.

Additional evidence of the exogenous impact of MTALTND4 comes from analyzing mitochondrial respiration in different respiratory states (Fig. 3*C,D*; supplementary table S7). In HeLa cells, 10 µM MTALTND4 significantly decreased both the basal respiration (routine respiration, ce-pept_R) and the maximal uncoupled respiration (ce-pept_E), but not the residual leak respiration (ce-pept_L) (Fig. 3*C*). In HEK-293T cells, both leak respiration and maximal uncoupled respiration, but not the routine respiration, were impacted in presence of 10 µM MTALTND4 (Fig. 3*D*). This is in line with the dose-dependent effect of the peptide and the higher threshold revealed for HEK-293T compared to HeLa cells (Fig. 3*A,B*). The analysis of the ‘spare respiratory capacity’ (SRC) – i.e. the difference between the basal respiration and the maximal uncoupled respiration (E−R) [(*17*) and reference therein)] – revealed a main effect of both factors ‘treatment’ (control vs peptide addition) and ‘cell type’ (HeLa vs HEK-293T), with no interaction between the two (Fig. 3*E*; supplementary table S7). In accordance with the general notion that cancer cells generally show lower SRC levels compared to their normal counterparts (*17*), our results suggest that HeLa cells have a physiological respiration much closer to their maximum capacity, thus a lower mitochondrial reserve compared to HEK-293T cells. A smaller reserve capacity likely makes these cells more sensitive to MTALTND4 treatment, as a lower decrease in maximal respiration is needed to impact the physiological routine respiration. In other words, HeLa cells were less able to withstand the inhibitory effect of MTALTND4 because when E-R reaches zero, mitochondrial energy production fails to meet the minimal needs of the cell.

The impact of MTALTND4 upon mitochondrial respiratory properties does not require cells to be intact as we examined its effect in permeabilized HEK-293T cells and observed that 10 µM MTALTND4 impacted both coupled (CI+II-pept_P) and uncoupled maximal respiration (CI+II-pept_E) sustained by CI+II-linked substrates. Although not significant, a sharp decreasing trend in O_2_ consumption was also revealed when testing cytochrome *c* oxidase standalone capacity (CIV-pept_E) (Fig. 3*F*; supplementary table S8). This result is of particular interest and deserves further investigations, as it may indicate that the site of action for MTALTND4 within the electron transport system (ETS) could reside in its final oxidase.

A change in reactive oxygen species (ROS – hydrogen peroxide) homeostasis was also observed. Intact HEK-293T cells in presence of 30 µM MTALTND4 showed an immediate, steady increase in H_2_O_2_ efflux but no increase in the total H_2_O_2_ scavenging capacity measured after 4 hours of incubation (Fig. 3*G,H*; supplementary table S9). An increased antioxidant capacity would be expected to lower cell oxidative stress. However, an alternative role for the revealed increased ROS efflux is also possible. Rather than being just deleterious by-products of an impaired mitochondrial respiration, mitochondrial ROS are pivotal mediators for many physiological processes including mitochondrial biogenesis, cell proliferation and differentiation, aging, apoptosis and response to hypoxia (*17, 18*). Given its impact on mitochondrial activity (potentially targeting CIV itself) and ROS metabolism, the potential role of MTALTND4 in cell adaptive response to stress such as hypoxia surely need further attention.

The reduced mitochondrial respiration was not accompanied by a parallel increase in anaerobic glycolysis to partially fulfill the cell energy demand. Four hours of exogenous treatment with MTALTND4 (30 µM) had no impact on the activity of lactate dehydrogenase in HEK-293T cells (Fig. 3*I*; supplementary table S9). Interestingly, despite a reduced mitochondrial capacity and no-change in lactic fermentation, the total ATP-content of treated cells increased (Fig. 3*L*; supplementary table S9). These results are potentially in line with a general cell metabolism undergoing downregulation, i.e. involving a coordinate downregulation of ATP-producing and ATP-demanding processes. This is further supported by the analyses of the effect of MTALTND4 (0.1 µM, 10 µM and 30 µM) upon HeLa or HEK-293T cell proliferation and viability at three different time points (24, 48 and 72 hours). A dose-dependent impact of the peptide upon cell proliferation was revealed at 48 hours, with a trend still perceivable at 72 hours although no more significant (Fig. 3*M*; supplementary table S10). The impact of the peptide upon cell viability was less pronounced, with no significant impact after 72 hours of treatment (blue trypan) (Fig. 3*N*; supplementary table S10), or with a small, but significant negative impact mainly for HeLa cells after 48h (alamarBlue) (fig. S5).

To summarize, we identified a novel human mitochondrial alternative peptide of 99 amino acids (i.e. longer than ATP8 [68 a.a.] and ND4L [98 a.a.]) whose sequence resides inside the *nd4* gene. MTALTND4, which is relatively well conserved in other primate species, is translated inside mitochondria and is also found in the plasma, and it appears to help regulate mitochondrial physiology. The sequence of MTALTND4 was detected by mass spectrometry after immunoprecipitation from HeLa cell extracts. Other proteins were also found that could be potential interacting partners of MTALTND4 (table S5). One of these proteins, the complement component 1q subcomponent binding protein (C1qbp), was also identified as an interacting partner using pull down assays, an interaction that was validated by Western blot (table S5). C1qbp, also known as p32, gC1qR, or HABP1, is a multifunctional and multicompartmental protein mostly localized in the mitochondrial matrix but also present at the cell surface, cytosol, and nucleus (*19, 20*). C1qbp is an evolutionarily conserved protein that can interact with a diverse array of mitochondrial and cellular, plasma, and microbial proteins (*20*). C1qbp regulates cellular energy metabolism through modulation of mitochondrial translation or through protein-protein interactions (*21, 22*). It has also a regulatory role in ROS production and can protect cells against oxidative stress (*20, 23*). This interaction could potentially explain the observed MTALTND4-mediated changes in mitochondrial respiration and ROS production. The lack of potential compensatory mechanisms accompanying the MTALTND4-mediated increase in H_2_O_2_ efflux and downregulation of mitochondrial respiration such as increased antioxidant activity and upregulation of anaerobic glycolysis, together with the no/little impact on cell viability, suggest that the effects of MTALND4 do not appear to be the result of a deleterious impairment of mitochondrial functions. This bioenergetic depression does however link with a sharp decrease in cell proliferation and increased total ATP content, suggesting that the cells might enter a sort of quiescent state.

Our findings have several implications. First, they suggest that mitochondrial genes in humans might have gone unnoticed. This calls into question the evolution of the mitochondrial genome as well as the selection pressures exerted on the mitochondria and the mechanisms allowing the translation of these alternative proteins. Our findings also offer a new framework for the investigation of mitochondrial diseases. Mitochondrial DNA mutations are responsible for a variety of human disorders due to defects in oxidative energy metabolism, and they are also involved in aging, aging-related diseases and multiple types of cancer (*24, 25*). The diversity of pathologies caused by mtDNA mutations is remarkable and the relationship between genotype and disease phenotype is not always straightforward. For example, mutations within genes sometimes cannot be directly linked to effects on oxidative energy metabolism or mitochondrial protein synthesis. Breton (*26*) proposed the mitochondrial Russian doll genes hypothesis to explain some of these discrepancies between theoretical expectation and experimental observation when studying mitochondrial diseases linked to mutations in the mtDNA. Indeed, molecular screening of mitochondrial disorders has been usually restricted to common harmful mutations (i.e. nonsynonymous or “non-silent” mutations) and deletions of mtDNA, i.e. without taking into account rare mutations or silent mutations in protein-coding genes (*27*). However, a silent mutation in a typical mitochondrial gene could be non-silent in an altORF. In other words, the existence of an alternative mitochondrial proteome could well explain some of the discrepancies observed previously, and therefore, this possibility clearly requires a systematic analysis. In sum, not only does this study enhance our understanding of mitochondrial biology, but it also provides a new mechanism through which mtDNA mutations might generally affect health. Collectively, we anticipate that more studies on the alternative mitochondrial proteome will expedite the discovery of new mitochondrial genes with key biological roles.

