## Supplementary Material and Text for "MTALTND4, a second protein coded by *nd4* impacts mitochondrial bioenergetics"

**SUPPLEMENTARY MATERIALS**

**Materials and Methods**

**Identification of mitochondrial smORFs and altORFs and choice of candidates for antibody production**. Three different approaches were used to search for potential smORFs and altORFs in the human mtDNA and to choose ORF-derived peptide sequences against which antibodies would be generated (table S1). First, an *in silico* search was performed for ORFs on both strands encoding peptides >20 amino acids using the vertebrate mitochondrial genetic code. However, we considered AGA and AGG as arginine codons as it was recently suggested that they are not normally used as stop codons in human mitochondria (*12, 13*) (see supplementary text 1). This analysis returned 249 sequences encoding 20 to ≈600 amino-acid-long peptides (the human mtDNA reference sequence GenBank accession number NC_012920 was used; see *28*). Of these sequences (i.e. 227 unannotated ORFs + the 13 typical protein-coding genes + humanin, SHLPs 1-6, Mots-c and gau [*29*]), 14 had a minimal conserved Kozak sequence known to be favorable for efficient translation initiation (*5, 10*), i.e. with an A or G at position -3 preceding the initiation codon and a G at position +4 (table S2). Of these 14 smORFs or altORFs, one was SHLP2, the small humanin-like peptide 2 encoded in the mtDNA 16S rRNA gene identified by Cobb et al. (*6*), and six were conserved outside the Homo/Pan (chimpanzees and bonobos)/Gorilla group, but two of them were too hydrophobic for immunisation. We decided to generate an antibody against the one candidate with the lowest number of hits showing >70% similarity with sequences of the human nuclear genome (tables S1 and S2).

In the second approach, we interrogated the OpenProt database (*30*). OpenProt is the first database that enforces a polycistronic model of eukaryotic genome annotations. Specifically, OpenProt retrieves transcripts from two standard annotations (Ensembl and NCBI RefSeq), which constitutes an exhaustive transcriptome. A three-frame *in silico* translation then yields the ORFeome: any ORF longer than 30 codons and starting with an ATG in any frame of any transcript. This ORFeome is then filtered to categorize predicted ORFs. The first filter retrieves all known protein (all ORF already annotated in Ensembl, NCBI RefSeq, and/or UniProtKB), these are the RefProts. The second filter looks at the homology of the currently not annotated ORFs with the RefProt of the same gene (if applicable) and retrieves novel predicted isoforms. The remaining ORFs encode novel proteins, called AltProts (*30*). For mitochondrial transcripts, OpenProt uses the vertebrate mitochondrial genetic code (but with AGA and AGR considered as stop codons). To increase confidence in ORF expression, OpenProt also cumulates several lines of evidence, such as (i) conservation evidence (for every ORF annotated, OpenProt identifies orthologs and paralogs across the 10 species currently supported by OpenProt), (ii) translation evidence (OpenProt retrieves publicly available ribosome profiling datasets and re-analyse them using the Price algorithm with a stringent 1 % FDR), and (iii) expression evidence (OpenProt retrieves publicly available mass spectrometry datasets and re-analyse them using multiple search engines, and a stringent 0.001 % FDR) (*30*). The search in OpenProt returned 14 altORFs encoding 29-61 amino-acid-long peptides, eight of them were found within the 12S and 16S ribosomal genes and six in protein-coding genes (table S3). We choose to focus on these latter six, which were all conserved in *Pan troglodytes* except for one altORF. Moreover, these six altORFs generated very few hits >70% similarity (i.e. less than 4) when searched against the nuclear genome (data not shown). Four of these altORFs were used for antibody production (tables S1 and S3).

In the third approach, we interrogated the CPTAC_Breast_Cancer mass spectrometry dataset to see if peptide sequences resulting from our first approach could be detected. Specifically, we used the novel tool PepQuery to interrogate spectra from the published CPTAC_Breast_Cancer dataset. This approach compares spectrum annotation with peptides from our novel sequences to annotation with any protein from the reference database (known proteins). We chose the CPTAC_Breast_Cancer dataset as it is one of the largest in terms of MS/MS scans with more than 32 millions. We first identified all possible peptides from our 249 proteins (see approach 1). We performed an *in silico* Trypsin digestion, allowing up to 2 miscleavages, a minimum peptide length of 7 amino acids and a mass from 400 to 6000 Da to mimic experimental conditions. Then, each peptide was queried using the PepQuery tool (see fig. S1).

From the PepQuery tool (*31*), we retrieved any Peptide Spectrum Match (PSM) that is better matched with our queried peptide than with any protein from the reference database. Various filters were used to validate the results as confident or not. For a PSM to be considered as confident, a *p-*value under 0.01 and a score higher than any peptide carrying any post-translational modification (PTM) from the reference database were needed. Using these criteria, we could be confident a given spectrum corresponds to our queried peptide and not to a reference peptide with any PTM (*31*). As such, supplementary table S4 contains for each of the 249 predicted proteins (approach 1) the number of PSMs with higher score than a referenced peptide, the number of confident PSMs, the best confident PSM (score), its corresponding *p-*value, and its total mass. This analysis returned 46 protein sequences with confident detection by mass spectrometry (including the 13 reference genes). Antibodies have been generated against 4 of the putative proteins (tables S1 and S4).

**Analyses of mtaltnd4 sequences**. To test for the presence of nuclear mtDNA sequences (NUMTs) identical or very similar to MTALTND4 due to recent mitochondrial transfer (*14*), we used the NCBI nucleotide basic local alignment search tool (BLASTN) (*32*) against the human genome reference (fig. S2). The degree of conservation of MTALTND4 was assessed by aligning the protein sequences from primate and mammal species obtained from the NCBI database. TBLASTn searches were used to ensure correct extraction of the protein sequences, which were aligned using ClustalW Multiple Alignment (*33*) (see Fig. 2 and fig. S3). Putative transmembrane (TM) helices were searched using TMPred (*34*), but none was found. Putative post-translational modifications (PTM) of MTALTND4 were searched using some of the PTM prediction webservers recently reviewed by (*35*), only scores with 80% probabilities were retained (fig. S6).

**Rabbit anti-MTALTND4 antibody generation and MTALTND4 peptide synthesis**. The custom rabbit anti-MTALTND4 was ordered from MediMabs (Montreal, QC, Canada) (and also our other antibodies; see Fig. 1 and table S1). High-titer polyclonal anti-sera against MTALTND4 was obtained. The complete MTALND4 peptide was synthesized, using Fmoc and Boc chemistry, microwave technology, and a proprietary solid support resin, by LifeTein (Somerset, NJ, USA). The purity of the peptide was verified by HPLC by the supplier. MTALTND4 (M.W. 11,532.06 gr∙mol^−1^) was shipped lyophilized to the University of Montreal and resuspended to a final stock concentration of 4 mM in ultrapure H_2_O following the manufacturer instructions. A working solution of 0.4 mM was prepared, and aliquots stored at −80°C for further experiments.

**Cell culture, HeLa rho0 cells and chloramphenicol treatments.** HeLa and HEK-293T cells were cultured in Dulbecco's Modified Eagle Medium (DMEM) supplemented with 10% calf bovine serum, penicillin, streptomycin and fungizone and were kept at 37˚C with 5% CO_2_. Chloramphenicol-treated HeLa cells were produced by adding 2ml of chloramphenicol diluted at 50µg/ml in 95% ethanol in a 10cm petri dish of 80% cell confluency. Cells were treated for 48 hours before being harvested for western blot analysis (*36*). Ethidium Bromide-treated HeLa-𝝆0 cell lysates (vs. control) were obtained from Abcam (ab154479).

**Western blotting**. HeLa or HEK-293T cells were washed in PBS, harvested, and lysed with a sonic dismembrator sonicator (Fisher) in a homogenizer buffer with a pH of 7.4 containing 10mM HEPES, 150mM NaCl and a cocktail of protease inhibitors (benzamidine, PMSF, aprotinin and leupeptin). Triton X-100 was added to 10% final concentration after sonication. Total protein concentration was estimated using the Bradford dosage method. The samples were then mixed with a Laemmli buffer containing DTT and ß-mercaptoethanol reducing agents and heated at 95˚C for 5 minutes before being loaded at equal protein concentration (100 µg) on Tricine-SDS-PAGE gels. SDS-PAGE gels were composed of three parts (separating, spacer and compaction gels) (*37*). To maximize the separation of low molecular weight proteins, the migration was first done for an hour at low voltage (30V). Then the voltage was increased to 43V, and the migration carried out overnight at room temperature. For the transfer, PVDF membranes of 2 µm were used which allows better retention of small proteins. The transfer was done for one hour at 1000 mA at 4˚C in a transfer buffer containing 194 mM glycine, 25 mM Tris-base and 20% methanol in H_2_0. Before incubation with the primary antibodies, membranes were blocked for 30 minutes at room temperature in a blocking buffer containing 5% milk powder and 0.05% tween-20 diluted in PBS. Primary antibodies rabbit anti-MTALTND4 (1:1000), mouse anti-ATP5 (1:1000; Abcam [ab14748]), mouse anti-Cox1 (1:500; Abcam) and mouse anti-actin (1:2000; Abcam [ab14705]) were diluted in a PBS + 0.05% tween-20 solution. Membranes were incubated with the primary antibodies for two hours at room temperature. After incubation, membranes were washed 3 X 5 minutes in TBS-T at room temperature. Secondary antibodies goat anti-rabbit IgG (1:2000) and 46 goat anti-mouse IgG (1:2000) coupled to the horseradish peroxidase (HRP) were diluted in a PBS + 0.05% tween-20 solution. Membranes were incubated with secondary antibodies for 30 minutes at room temperature. Finally, membranes were washed 3 X 10 minutes with TBS-T and 1 X 5 minutes with TBS at room temperature. Protein signals were visualized by adding Montreal Biotech Inc.’s enhancer and substrate solutions and images were captured with a FUSION FX chemiluminescence imaging system.

To verify if predicted post-translational modifications (fig. S5), i.e. glycosylation and phosphorylation, were responsible for the observed MTALTND4 band (~25kDa) at higher MW than expected (11,5kDa), whole cell lysates were treated with a complete protein deglycosylation mix II kit (NEB, Cat# P6044) following the manufacturer’s protocol, as well as with a lambda protein phosphatase (NEB, P0753S) following the manufacturer’s protocol (fig. S7). To verify the specificity of the anti-MTALTND4 antibody, the synthetic antigenic peptide was added to the primary antibody solution at a 10X concentration before the incubation, to competitively chelate every antigenic site of the primary antibody and to show the attenuation of the specific MTALTND4 band (fig. S7). To verify if a SDS resistant dimer could be responsible for the observed MTALTND4 band at higher MW than expected, we also exposed our samples to different concentrations of ß-mercaptoethanol (0%, 5% and 10%) and different heating times (5min, 30min and 3 hours) (*38*) (fig. S7). The presence of MTALTND4 in the plasma was verified by western blotting.

**Immunofluorescence**. HeLa cells were cultured on an 8-well PCA detachable microscope slide in DMEM supplemented with 10% calf bovine serum, fungizone, penicillin and streptomycin and were kept at 37˚C with 5% CO_2_. Each chamber was washed with PBS and fixed with 4% paraformaldehyde for 20 minutes. Cells were then washed 3 X 5 minutes in PBS before their membranes were permeabilized by adding 0.2% Triton for 4 minutes. Cells were washed 3 X 5 minutes with PBS and then blocked with 5% NGS in PBS for 30 minutes at room temperature before incubating for 1 hour with primary antibodies rabbit anti-MTALTND4 antibody (1:50) and mouse anti-ATP5 antibody (1:250; Abcam) diluted in 1% NGS in PBS. Cells were washed 3 X 5 minutes in PBS before incubating with the secondary antibodies (Alexa Fluor ™ 488 goat anti-rabbit IgG [1:500] and Alexa Fluor ™ 594 goat anti-mouse IgG [1:500]) in PBS containing 1% NGS for 1 hour at room temperature. Finally, cells were washed 3 X 5 minutes in PBS before adding a mounting drop containing DAPI and visualized with an EVOS M5000 microscope (fig. S4) or Zeiss-LSM800 (Fig. 2F).

**Immunoprecipitation and mass spectrometry analysis.** HeLa cells were washed with PBS and harvested in a lysing buffer containing 10mM HEPES, 150mM NaCl and a cocktail of protease inhibitors (benzamidine, PMSF, aprotinin and leupeptin). Cells were then lysed with a sonic dismembrator sonicator (Fisher) 3 times for 30 seconds before adding 100 µl of Triton X-100 10%. Cell lysates were incubated on ice for 20 minutes and then centrifuged for 15 minutes at 15000 RPM at 4˚C. The supernatant was kept. 20 µl of protein A agarose was added to 1ml of samples and incubated at 4˚C for 1 hour. Samples were then centrifuged at 15000 rpm for 1 minute at 4˚C and the supernatant was kept. 20 µl of rabbit anti-MTALTND4 antibody and 20 µl of protein A agarose were added to the samples before incubating at 4˚C overnight. 20 µl of rabbit pre-immune serum was used as control. Samples were washed 6 times with lysing buffer. Samples were then sent to the proteomics platform of the McGill University Health Center (MUHC) Research Institute for mass spectrometry analysis. Data were analyzed with Mascot (Matrix Science) against a human database and a final report was generated using Scaffold5 (Proteome Software) (table S5).

**Pull down assay and mass spectrometry analysis.** Transformation of competent cells and protein expression were performed following manufacturer’s instructions (Thermo Scientific, BL21(DE3) Competent Cells). For the transformation, 1 µl of GST-tagged MTALTND4 expression plasmid (LifeTein) was added to 50 µl of BL21(D3) competent cells originating from an *E. coli* B strain. A GST expression plasmid was used as control. Cells were incubated on ice for 30 minutes, then heat-shocked for 30 seconds in a 42°C water bath before returning on ice for 2 minutes. 250 µl of LB broth growth medium was then added, and the tube was placed in a shaking incubator at 225 rpm for 1 hour at 37°C. The transformed cells were then spread on previously prepared LB plates containing 25 µl of ampicillin and incubated at 37°C overnight. For protein expression, a transformant colony was picked and inoculated in 5 ml of LB medium with 5 µl ampicillin. The culture was incubated overnight with shaking at 37°C, allowing it to grow to saturation. 200 µl of overnight culture was added to 10ml of fresh LB medium containing 10 µl of antibiotic. The culture was incubated for 2 hours, 10 µl of IPTG was then added, and it was incubated for an additional 2 hours. The induced culture was centrifuged at 2500 rpm for 10 minute and the pellet was resuspended in 2 ml of PBS 1X with 4M urea. A cocktail of protease inhibitors was added, and the pellet was lysed as described above for the HeLa cells.

For protein extraction, the bacterial lysates were incubated overnight with 100 µl of a 50% slurry of glutathione sepharose beads (Abcam) at 4°C for pre-coupling. The beads were washed 3 times with PBS 1X and the protein quantity assessed via a Bradford protein assay. HeLa and HEK-293T cell lysates were then prepared as described above and pre-cleared using 50 µl of glutathione sepharose beads and 25 µg of GST protein (Sino Biological) for 2 hours at 4°C with end-over-end mixing. The lysates were then centrifuged for 2 minutes at 15000 rpm at 4˚C and the supernatant was collected. From the pre-coupled MTALTND4 protein, 10 µg was added to 1 ml of the pre-cleared HeLa or HEK lysates and incubated overnight at 4°C. Samples were washed 6 times with lysing buffer. Samples were then sent to the proteomics platform of the McGill University Health Center (MUHC) Research Institute for mass spectrometry analysis. Data was analyzed with Mascot (Matrix Science) against a human database and a final report was generated using Scaffold5 (Proteome Software) (table S5).

The validation of a predicted interaction with the complement component 1q subcomponent binding protein (C1qbp) was done via Western blot as described above (primary antibody mouse anti-GC1q (1:1000; Abcam [ab24733]), was diluted in a PBS + 0.05% tween-20 solution and secondary antibody goat anti-mouse IgG (1:2000) coupled to the horseradish peroxidase (HRP) was diluted in a PBS + 0.05% tween-20 solution).

**Cell proliferation and viability**. HeLa or HEK-293T cells were treated with complete synthetic MTALTND4 (0,1μM, 10μM or 30 μM) or water (control) in DMEM low glucose buffer for 24, 48 and 72 hours. Cell proliferation (*n*=3) was evaluated directly by counting manually with a hemocytometer. To test for cell viability (*n*=3), 10 μL of 4% trypan blue exclusion dye were added to 10 μL of treated or untreated HeLa cells and examined under a microscope. Numbers of viable cells were estimated using a hemocytometer. Cell viability (n=5) was also determined by alamarBlue assay kit from ThemoFisher (DAL1025) following the manufacturer’s protocol.

**High resolution respirometry (HRR).** Cells (~80% confluency) were tripsinized (2.5%), washed, centrifuged, and finally resuspended in MiR05 mitochondrial respiratory buffer (110 mM D-sucrose, 60 mM lactobionic acid, 20 mM taurine, 20 mM HEPES, 10 mM KH_2_PO_4_, 3 mM MgCl_2_, 0.5 mM EGTA, BSA 1 g∙L^-1^). MiR05 was supplemented with 5 mM pyruvate (MiR05+P) as external energy source to support intact cell respiration (*39*). Cell viability (%) and concentration (Mx cells∙mL^−1^) were determined through a Neubauer hemocytometer and trypan blue staining.

Mitochondrial respiration has been characterized by high resolution respirometry (HRR), using a dedicated Oxygraph-2k with DatLab software v7.4 (Oroboros Instruments, Innsbruck, Austria). The respiratory chambers were preloaded with 2.1 mL MiR05+P respiratory medium and calibrated for oxygen saturation at 37°C. Collected cells were transferred into each respiratory chamber by replacing part of the chamber media, reaching 2.5 mL total chamber volume and cell concentration of 0.5-0.8 Mx cells∙mL^−1^, depending on the cell line and the specific experiment (*39, 40*). Cell suspensions were continuously stirred at 750 rpm and part of the solution (400 µL) was removed, sonicated (5 times 1 second, on ice), and stored at −80°C for protein determination. Chambers (final volume 2.1 mL) where then sealed, and oxygen consumption monitored. Data were corrected for instrumental background O_2_ flux measured at 37°C in a separate experiment (*39, 40*). Mitochondrial respiration was investigated in both intact (ce) and permeabilized (pce) cells.

***HRR in intact cells.*** Oxygen consumption in intact HeLa and HEK-293T cells has been monitored following sequential titration of MTALTND4 peptide. After recording the respiration of intact cells (ROUTINE, R-state) in the presence of 5 mM pyruvate (ce_R), increasing peptide concentrations were tested. This included 0.1 µM, 1 µM, 5 µM, 10 µM, 15 µM, 20 µM, 25 µM and 30 µM for *n* = 4 HeLa, and 0.1 µM, 1 µM, 5 µM, 10 µM, 15 µM, 20 µM, 25 µM, 30 µM, 35 µM, 40 µM and 45 µM for *n* = 3 HEK-293T. Simultaneous controls were treated with an equal volume of H_2_O titrated at the same time points. Respirometry data were normalized for an internal parameter, the ROUTINE (R) respiration - i.e. the respiration rate of intact cells before treatment - and expressed as flux control ratios (FCR). For each titration point, data are expressed as fraction of the control respiration measured in parallel at the same time. The exogenous impact of peptide addition was also assessed on intact HeLa (*n*=6-9) and HEK-293T (*n*=6) cells following a specific coupling control protocol (CCP). After the measurement of ROUTINE respiration in the presence of 5 mM pyruvate (ce_R), the exogenous impact of MTALTND4 was assessed at a final concentration of 10 µM (ce_pept_R). Addition of the ATP-synthase inhibitor oligomycin (15 nM) declined the respiration to the level of LEAK respiration (L, Leak state - state 4 or 2’) and allowed the measurement of the residual respiration due to the futile proton cycle (ce_pept_L). The maximal respiration was then stimulated by the titration of FCCP (0.125 µM each step) (ce_pept_E). Finally, the residual oxygen consumption (ROX) was determined in presence of 2.5 µM antimycin A (complex III inhibitor) and subtracted from the data. The ‘spare respiratory capacity’ (SRC) was obtained for both HeLa and HEK-293T cells by subtracting routine respiration from the maximal uncoupled respiration (E−R) (*n*=6-6). Data were normalized for the ROUTINE respiration and expressed as flux control ratios.

***HRR in permeabilized cells.*** HEK-293T cells (*n*=6) were added intact to each chamber and ROUTINE respiration (ce_R) was determined in presence of complex I (CI) substrates pyruvate (5 mM), malate (2 mM) and glutamate (10 mM). The plasma membrane of cells was then selectively permeabilized with 7.5 µg∙mL^−1^ digitonin, and Leak respiration (Leak, L-state - state 4 or 2’) in presence of CI-linked substrates measured (CI_L). The optimum digitonin concentration for complete plasma membrane permeabilization of cultured cells was determined empirically following existing protocols (*39-42*). Addition of 2.5 mM ADP stimulated mitochondrial respiration (OXPHOS, P-state or state 3) sustained by CI substrates (CI_P). To verify the intactness of mitochondrial membranes, cytochrome *c* (10 µM) was added to the chambers (c_P). Addition of complex II (CII) linked substrate succinate (10 mM) fueled the OXPHOS activity, now sustained by both CI and CII substrates (CI+II_P). This represents the maximal coupled respiration and data presented as flux control ratios have been normalized for it. Graphical representations of the data start from this point onward. The impact of MTALTND4 was tested against the CI+II sustained coupled respiration, at a final peptide concentration of 10 µM (CI+II-pept_P). Further addition of FCCP (titration, 0.125 µM each step) stimulated the maximal uncoupled respiration (ETS, E-state or state 3u) (CI+II-pept_E). Residual oxygen consumption (ROX) was achieved in presence of 0.5 µM rotenone (Rot - complex I inhibitor) and 2.5 µM antimycin A (complex III inhibitor), the latter subtracted from the data. Cytochrome *c* oxidase (complex IV, CIV) standalone activity was measured in presence of 2 mM ascorbate and 0.5 mM TMPD (CIV-pept_E). The residual oxygen consumption due to autoxidation was determined with 20 mM sodium azide (complex IV inhibitor) and subtracted from CIV activity.

**Hydrogen peroxide efflux.** The rate of hydrogen peroxide (H_2_O_2_) efflux from intact cells was determined using the Amplex® Red Hydrogen Peroxide Assay Kit (Invitrogen, A22188). HEK-293T cells (*n*=5) were collected and suspended in the respiratory buffer MiR05 supplemented with 5 mM pyruvate at a concentration of 5 Mx cells∙mL^−1^. Each sample was split in two (control and treatment). Before the treatment, homogeneous aliquots were collected, lysed through sonication (5 times 1 second strokes on ice) and kept frozen at −80°C for further protein determination. Intact cells were incubated with either 30 µM MTALTND4 (*n*=5) or the same volume of H_2_O for the paired controls (*n*=5) for 30 minutes at 37°C. After peptide exposition, samples were transferred to a black microplate and incubated in the dark for 30 minutes at 37°C in the presence of 0.01 mM Amplex® Red, 0.1 U∙mL^−1^ horseradish peroxidase (HRP), 50 mM sodium phosphate buffer, pH 7.4. The H_2_O_2_ efflux was then measured following the increase in fluorescence (excitation 560nm; emission 615nm) for 1.5 hours. Hydrogen peroxide efflux rate was determined with the use of a standard curve and normalized for the amount of proteins (pmol H_2_O_2_ ∙ mg proteins^−1^ ∙ minute^−1^).

**Enzymatic activities and ATP content.** HEK-293T cells (10 Mx cells∙mL^−1^) were incubated for 4 hours at 37 °C in MiR05 supplemented with 5 mM pyruvate and 30 µM MTALTND4 (*n*=5). Each sample was split and had its own control, with H_2_O used instead of the peptide (*n*=5). Cells were lysed (5 times 1 second strokes on ice) using a sonic dismembrator (Fisher) and kept frozen at −80°C for further analysis of enzymatic activities and ATP content.

The rates of enzymatic activity were measured spectrophotometrically with a Mithras LB940 microplate reader (Berthold technologies, Germany) and MikroWin 2010 software (Labsis Laborsysteme, Germany). Enzymatic capacities were expressed as µmol of substrate transformed to product per minute (Units - U) over mg of proteins. Enzymatic assays were performed in duplicates at 37°C in the following conditions:

***Catalase (CAT, EC 1.11.1.6).*** CAT activity was estimated as the total H_2_O_2_ scavenging capacity. It was measured at 240 nm following the disappearance of H_2_O_2_ (ε_240_ = 43.6 M^−1^∙cm^−1^) for 1 minute. The medium was composed of 50 mM potassium phosphate, 0.1% (v/v) triton X 100 and 60 mM H_2_O_2_, pH 7.0. The background activity in absence of sample was subtracted from the main results (*43, 44*).

***Lactate dehydrogenase (LDH, EC 1.1.1.27).*** LDH activity was determined by the rate of NADH oxidation (ε_340_ = 6.22 mM^−1^∙cm^−1^) following pyruvate reduction to lactate. The medium was composed of 50 mM potassium phosphate, 0.16 mM NADH and 0.4 mM pyruvate, pH 7.0. The specificity of the reaction was tested in the presence of 30 mM oxamate (LDH inhibitor) (*45*).

***ATP content.*** ATP was quantified using the ATP Determination Kit (Sigma A22066). The bioluminescence assay follows the ATP-dependent light production in a reaction involving a recombinant firefly luciferase and its substrate D-luciferin. The conditions were set as follow: 0.5 mM D-luciferin, 1.25 μg/mL firefly luciferase, 1 mM dithiothreitol, 5 mM MgSO_4_, 100 μM EDTA, 100 μM sodium azide, 25 mM tricine buffer, pH 7.8. The luminescence was measured in duplicates immediately after sample addition, in a luminometer (Mithras LB940, Berthold technologies, Germany) set at 28°C. The content of ATP in the sample was derived using a standard curve with known concentrations of ATP, to which the background luminescence recorded in absence of sample was subtracted. Data are presented as ATP over total protein content (nmol ATP ∙ mg proteins^−1^).

***Protein content.*** The total level of proteins (mg∙ml^−1^) of each sample was determined in duplicate through the Bicinchoninic Acid (BCA) assay (Bicinchoninic Acid Protein Assay Kit, Sigma BCA1) using a Bovine Serum Albumin (BSA) standard curve (*46*).

**Data analysis.** High resolution respirometry data were collected and preliminary analysed using DatLab v7.4 (Oroboros Instruments, Innsbruck, Austria), while enzymatic activities, ROS efflux rate, ATP and protein content with MikroWin 2010 software (Labsis Laborsysteme, Germany). Graphical representation and statistical analyses were performed using the R software (*47*) and ggplot2 package (*48*). The normality and homoscedasticity of data were verified with Shapiro and Levene’s tests respectively. The dose-dependent impact of peptide addition upon mitochondrial respiration was determined at each titration point using a one sample *t* test, comparing each treatment with its respective parallel control. For all the parameters included in the CCP and SUIT protocols, as well as for enzymatic activities (CAT and LDH), ATP content and H_2_O_2_ efflux rate, the effect of peptide addition was tested with the means of paired *t-*test or Welch-test. A linear mixed model was implemented for the parameters SRC, proliferation and viability. The factors ‘cell type’ and ‘treatment’ (effect of peptide addition) were considered as fixed effects when analysing HeLa and HEK-293T SRCs. The only factor ‘treatment’ was considered for HEK-293T cell proliferation and viability. In both cases, cell ID was implemented as random effect (paired samples). The significance of both factors and their possible interaction was determined through a Type III ANOVA and *post hoc* multi comparison. Data are graphically presented as mean ± standard error of the mean (s.e.m). Statistical significance was set at *p* ≤ 0.05 and *p*-values were corrected with Holm adjustment for multiple testing. Detailed summaries are provided in supplementary tables S6-9.

**Supplementary Text**

***Supplementary Text 1***

Mitochondrial genetic code

The codons AGA and AGG, which conventionally code for arginine, were thought to be recognized as termination codons in vertebrate mitochondria because of the absence of a mtDNA-encoded tRNA^ARG^ for these triplet codons, but it has now been shown that this is not the case at least in humans (*12, 13*). The only two human mitochondrial genes with AGA and AGG as putative terminators are *cox1* and *nad6* and it has been demonstrated that their two mRNAs would rather terminate at a UAG stop codon created because of a −1 frameshift of the mitoribosome (*12, 13*). If this was also the case for *mtaltnd4*, the translated peptide would be 59 amino-acids long with an expected size of 6.9 kDa. Alternatively, with AGA and AGG coding and a translation occurring either within the cytoplasm, as suggested for MOTS-c because of its tandem ATGAGG start and “putative stop” codons using the typical mitochondrial genetic code (*5*), or in mitochondria, as suggested for some of the SHLPs that contain putative AGA and AGG coding triplets (*6*), MTALTND4 consists of 99 amino acids with an expected size of 11,5 kDa and no predicted transmembrane helix. In accordance with some of the SHLPs (*6*), our results imply a possible import of the cytosolic tRNA^AGR^ into the mitochondria, an innate ability of mammalian mitochondria (*15, 16*). This scenario would also parsimoniously explain the previously reported intra-mitochondrial localization of endogenous MOTS-c (*5, 7*), suggesting that the peptide is translated in the mitochondria rather than imported after the translation in the cytoplasm of its exported mRNA. Further experiments, such as a treatment of cultured cells with chloramphenicol, will be needed to test this hypothesis for MOTS-c.

***Supplementary Text 2***

Conservation of MTALTND4

MTALTND4 is better conserved in primates (Fig. 2B) than in mammals in general (fig. S3). One could expect that an altORF of great importance to mitochondrial/cellular function would be highly conserved in animals. However, mitochondria-derived micropeptides such as Humanin, MOTS-c and SHLPs that appear to modulate mitochondrial and cellular biology, as well as global physiology, are not well conserved among mammals or vertebrates (*5, 49, 50*). For example, pseudogenization of the Humanin gene is common in the mtDNA of many vertebrates (*49*). Also, several subunits of the OXPHOS complexes encoded by the nucleus, although having key roles in the assembly or activity of the complex or in mitochondrial respiration, are not found in all animal species (*51, 52*).

The process leading to the creation of new proteins by mutations within a coding sequence that lead to the expression of a novel protein in another reading frame is called “overprinting” (*53, 54*). For mitochondria, with their reduced genome size and constraints on protein birth by duplication or fusion of existing coding sequences, overprinting maybe more frequent (*53, 54*).

**Supplementary Figures**

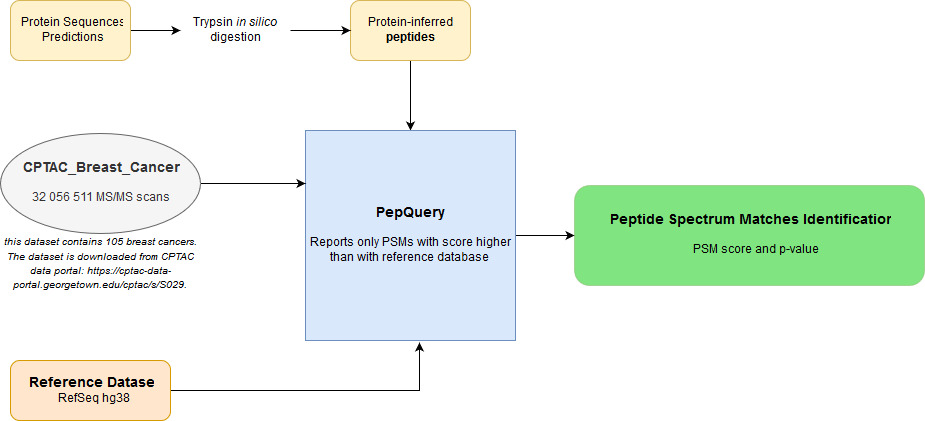

**Figure S1.** Using the PepQuery tool to interrogate spectrums from published proteome datasets.

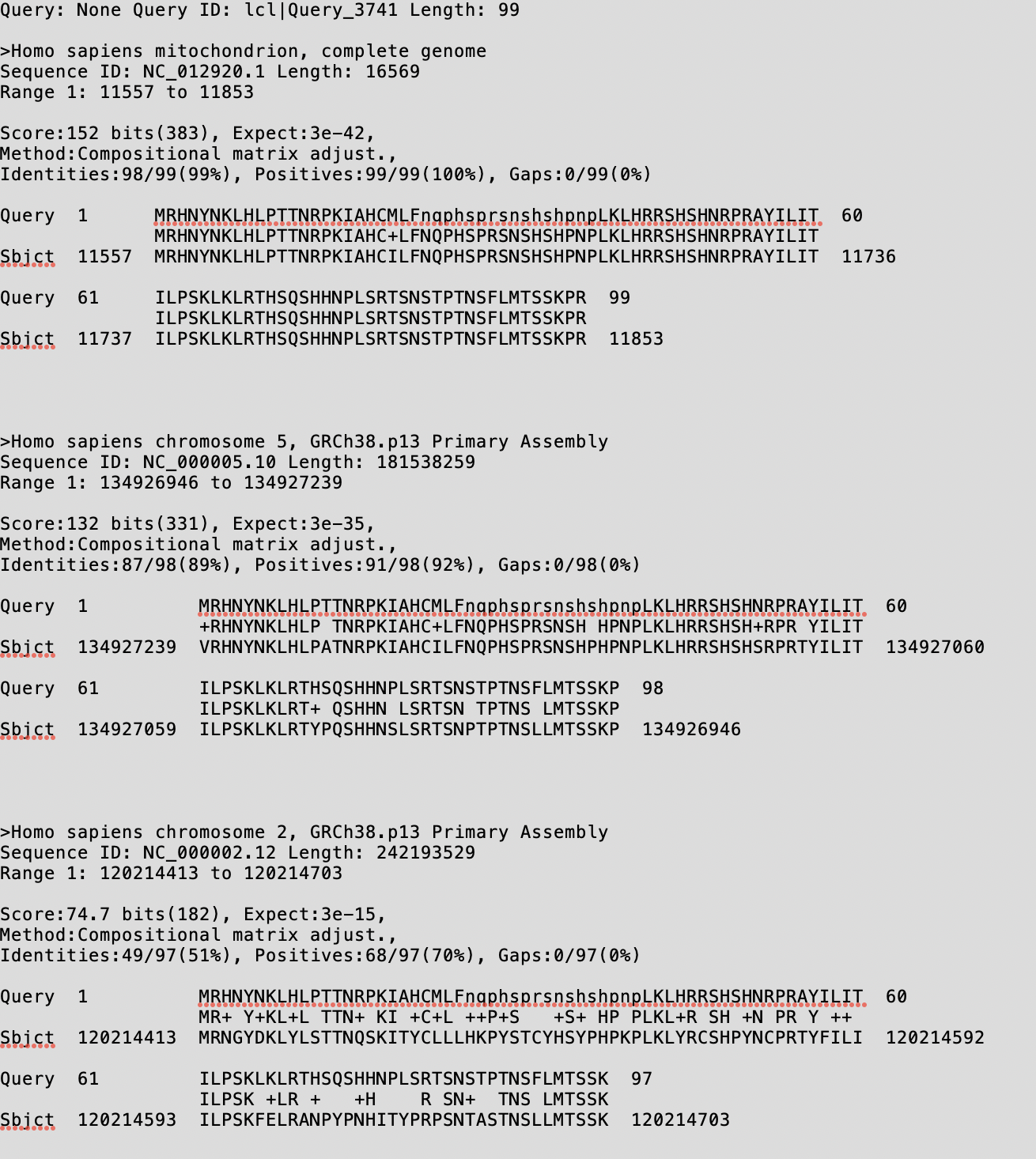

**Figure S2.** Comparison of MTALTND4 peptide encoded in mitochondrial DNA (mtDNA) and hypothetically in nuclear DNA that have been transferred from mtDNA (NUMT) through evolution. Only sequences with >50% identities and complete without stop codons are shown.

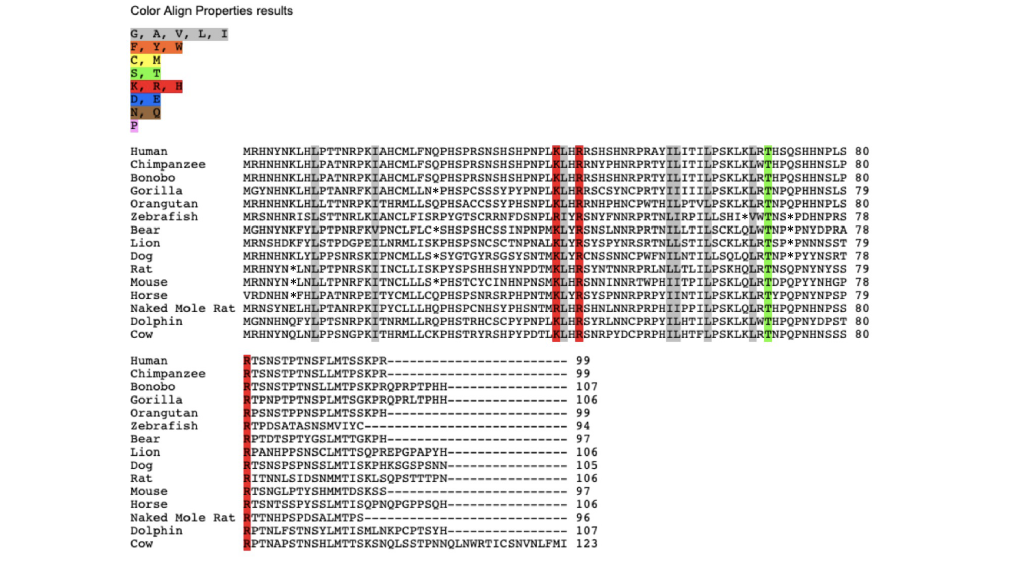

**Figure S3.** Multiple peptide sequence alignment in 15 mammal species: human (*Homo sapiens*), chimpanzee (*Pan troglodytes*), bonobo (*Pan paniscus*), gorilla (*Gorilla gorilla*), orangutan (*Pongo pygmaeus*), mouse (*Mus musculus*), rat (*Rattus norvegicus*), naked mole rat (*Heterocephalus glaber*), dog (*Canis lupus familiaris*), cow (*Bos taurus*), zebrafish (*Danio rerio*), lion (*Panthera leo*), bear (*Ursus arctos*), horse (*Equus caballus*), dolphin (*Tursiops truncatus*). *Indicates a stop codon.

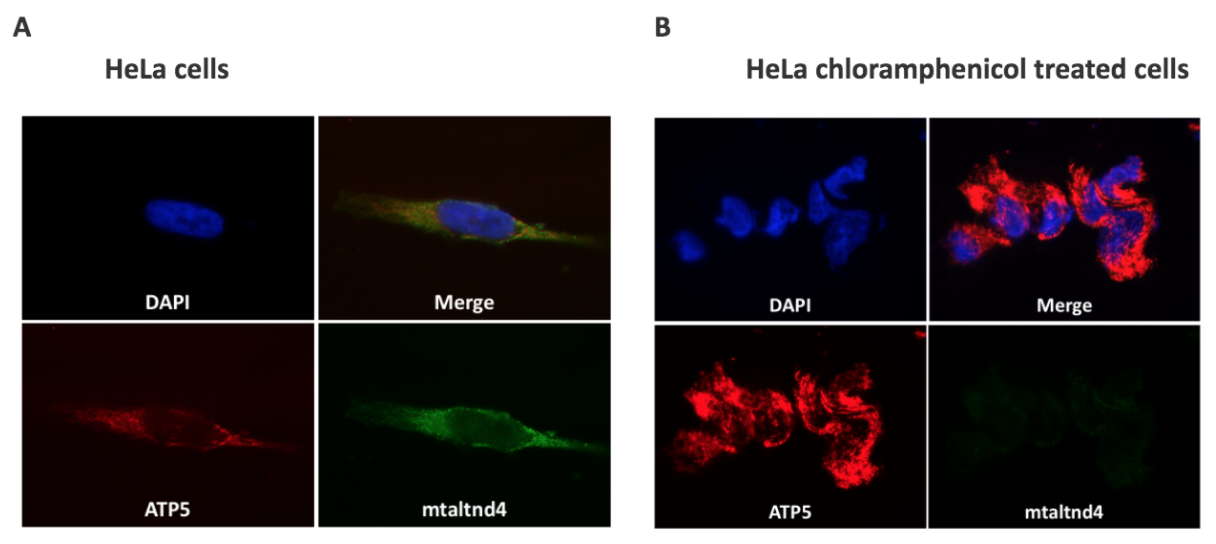

**A**

HeLa cells treated with chloramphenicol

**B**

HeLa cells

**Figure S4.** An alternative protein in the mtDNA-encoded *nd4* gene. *(A)* Detection of MTALTND4 and nuclear-encoded mitochondrial ATP5 by immunofluorescence in HeLa cells. *(B)* Detection of MTALTND4 and nuclear-encoded mitochondrial ATP5 by immunofluorescence in HeLa cells treated with chloramphenicol.

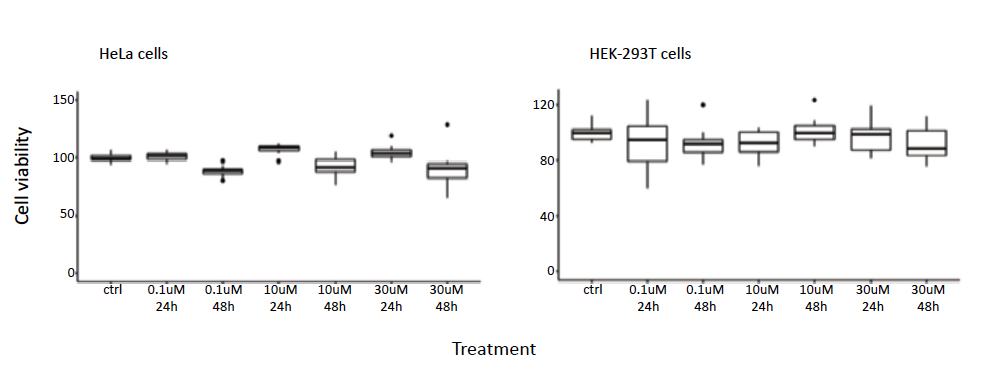

**Figure S5. Effect of MTALTND4 on cell viability.** HeLa and HEK-293T cells were cultured in low glucose DMEM with 0,1 uM, 10 uM or 30 uM MTALTND4 peptide or water (control) for 24h or 48h and assessed for cell viability using the alamarBlue assay (n = 3 for each cell line).

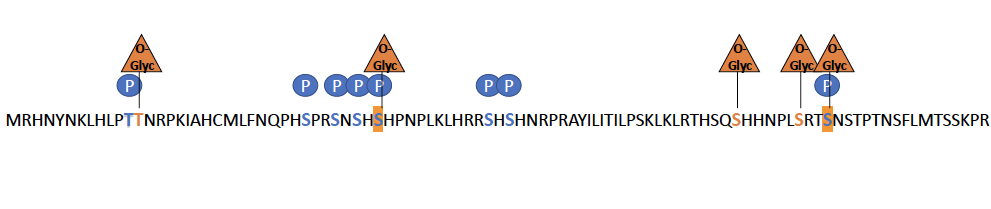

**Figure S6.** Putative post-translational modifications of MTALTND4. O-Glyc: putative O-glycosylation; P: putative phosphorylation. PTM prediction were done using webservers reviewed by (*35*), only scores with 80% probabilities were retained.

**A.	B.**

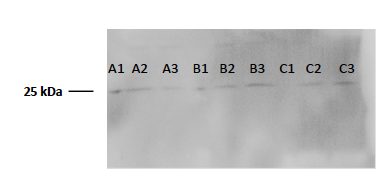

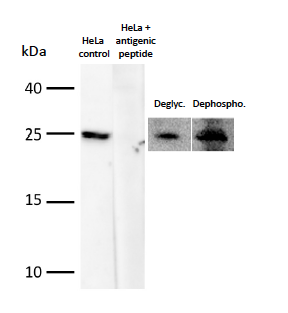

**Figure S7. A.** Specificity of the anti-MTALTND4 antibody and detection of putative post-translational modifications of MTALTND4. Deglyc: deglycosylation (whole cell lysates were treated with a complete protein deglycosylation mix II kit [NEB, Cat# P6044] following the manufacturer’s protocol); Dephosphos: dephosphorylation (whole cell lysates were treated with a lambda protein phosphatase [NEB, P0753S] following the manufacturer’s protocol). **B.** Apparent molecular weight of MTALTND4 exposed to different concentrations of denaturing agent 𝜷-Mercaptoéthanol and different heating times at 95˚C. A = 5 minutes, B = 30 minutes and C = 3 hours. 1 = 0% 𝜷-Mercaptoéthanol, 2 = 5% 𝜷-Mercaptoéthanol and 3 = 10% 𝜷-Mercaptoéthanol.

**Supplementary Tables**

**Table S1. Chosen candidates for antibody production.**

| Approach  used | Amino acid sequence (antigenic sequence in bold underlined) | Length | Predicted MW | Position in the reference human mtDNA |
| --- | --- | --- | --- | --- |
| In silico & Kozak | MGC**SGSSVSQCYRVHTPQTKMPNAWRAPVSG** | 31aa | 3,5kDa | non-coding region anti-sense  117 to 24 |
| OpenProt  IP_306387 | IFYLSRPRNKHASFYSSSNQKNKPSFHRSCHQVFPHASNRIHNPSNSYPLQQYTLRTMNHNQYYQSMLIINNHNSYSNKTRNSPLSL LSPRGYPRHPSDIRPASSHMTKTSP**HLNHMPNLSLTKRKP**SPHSLNLIHHSRQLRWIKPNPATQNLSMLLNYPHRMNNSSSTVQP  *The sequence is presented with AGA and AGG codons translated to arginine, and the sequence in OpenProt is in yellow (OpenProt considers AGA and AGG as stop codons for the mitogenome). | 172aa | 20kDa | within nd2  sense, frameshift  4547 to 5063 |
| OpenProt  IP_306389 | ICNNLLHSNTHHNRRLWQLTSSPNNRCPRYGVSPHKQHKLLTLTSLSPTPARICYSGGRSRNRLNSLPSLSRELLPPWSLRRPNHLLL TPSRCLLYLRGHQFHHNNYQYKTPCHNP**MPNAPLRLIRPNHSSPTSPISPSPSCWHHYTTN**RPQPQHHLLRPRRRRRPHSMPTPILI FRSPWSLYSYPTRLRNNLPYCNLLLRKKRTIWMHRYGLSYDINWLPRVYRVSTPYIYSRNRRRHTSMFHLRYHNHRYPHRRQSI  *The sequence is presented with AGA and AGG codons translated to arginine, and the sequence in OpenProt is in yellow (OpenProt considers AGA and AGG as stop codons for the mitogenome). | 259aa | 31,2kDa | within cox1  sense, frameshift  6089 to 6866 |
| OpenProt  IP_306398 | **MSSSKPHLSPPWLSSPDEATSQNAWTQAHTSYSTP**  *The sequence is presented with AGA and AGG codons translated to arginine, and the sequence in OpenProt is in yellow (OpenProt considers AGA and AGG as stop codons for the mitogenome). | 35aa | 3,8kDa | within nd4  sense, frameshift  11115 to 11219 |
| OpenProt  IP_306403 | MAESSATFTPMAPQYSLSASSYTS**GEAYITDHFSTQKPET**SALSSCLQL  *The sequence is presented with AGA and AGG codons translated to arginine, and the sequence in OpenProt is in yellow (OpenProt considers AGA and AGG as stop codons for the mitogenome). | 49aa | 5,2kDa | within cytb  sense, frameshift  14970 to 15116 |
| MS/MS | MGLSRIEGLFGQVVCGGLGMCFLVLHRAIIGMWLVCWLVGLVWGALWSG**SEITWLGRRSLGGLRGP**LLGVMGWVLLYDRHVIGGSLCVVVQVEAY | 95aa | 10,3kDa | cox3  anti-sense  9462 to 9177 |
| MS/MS | MWSLPR**RLPGWPSSARMRRLRAVPRTPAHAPNNRY**SVPMSLWFVENSQRSANISGGEVKWLSEALDCKSKDRG | 73aa | 8,3kDa | non-coding region  anti-sense  6061 to 5842 |
| MS/MS | IMRMTAPVKLQGVWMRMAVTTRAMWLIEEYAMSDFRSVCRRQ**MELVMIMPHRDSTRKG** | 57aa | 6,9kDa | nd4  anti-sense  11709 to 11535 |
| MS/MS | MRHNYNKLHLPTTNRPKIAHCMLFNQPHS**PRSNSHSHPNPLKLHRRSHSHNRPR**AYILITILPSKLKLRTHSQSHHNPLSRTSNSTPTNSFLMTSSKPR | 99aa | 11,5kDa | within nd4  sense, frameshift  11557 to 11854 |

**Table S2.1 Identification of mitochondrial smORFs and altORFs - in silico + kozak approach**

Vertebrate mitochondrial genetic code Start: ATG, ATT, ATA

Stop: TAA, TAG

AGA, AGG coding

| START | STOP | LONG. | CODON I | CONTEXT | Kozak -3 | Kozak +4 | CODON F | **FEATURE** | PROTEIN SEQUENCE |
| --- | --- | --- | --- | --- | --- | --- | --- | --- | --- |
| 49 | 199 | 50 | ATT | TCTCCATGC ATT G | T | G | TAA |  | IWYFRLGGMHAMALRDAGAGAPYVAVSVFDSC LILLFIAPTFNITGEHTY |
| 232 | 334 | 34 | ATA | TTGTAGGAC ATA T | G | T | TAA |  | MMMTIECLHSHFPHRHHNKKFPPNPPSPASGH ST |
| 448 | 832 | 128 | ATT | AACTAACAC ATT T | C | T | TAG |  | IIFPSHSHTTNLINTTPAHPTQHTHTAANPMPRT NQTPKTPPTVYVAYLLKAMHWKCLDGLTSPHKQ MGLVLAFLLALSKITHASIPVPVSSPSKSPRSKGTSI KHAAMQLKTLSLATPPRETAVINL |
| 967 | 1195 | 76 | ATA | CCCTCCCCA ATA A | C | A | TAG | **RNR1** | MKLKLTWVVKNSSWHKMDYESGFNMSEHTMA KTQTGIRYPTMLSPKPQQLNQQNCSPEHYEPQL KTQRTWRCFMSL |
| 1258 | 1426 | 56 | ATA | TCAGCCTAT ATA C | T | C | TAG | **RNR1** | MPPSSANPDEGYKVSASTHVKTLGQGVAHEVAR NGLHFLPQKTTMALMKLKGRRWI |
| 1540 | 1609 | 23 | ATT | CCCCTACGC ATT A | C | A | TAG | **RNR1** | IYMEETSRNMVSVLESALGRTRV |
| 1963 | 2038 | 25 | ATA | TGTAGCAAA ATA T | A | T | TAG | **RNR2** | MVGRFMGRGDKPTEPGDSWLSKMES |
| 2323 | 2563 | 80 | ATT | CATGAAAAC ATT T | A | T | TAA | **RNR2** | ILLRMSLRQIKTLNWQLTAQYLQSTNKSLLPSLST QHRHAHKERLKKVKGTRQILPRLFTKNITSSITSIR GTACPVTHV |
| 2617 | 2800 | 61 | ATA | GTTCCTTAA ATA G | T | G | TAA | **RNR2** | MGTCMNGSTRVQLSLTFNQWNWPAREEAGM TQQDEKTLWSFNLLMQTVPNKPTGPKLPNLH |
| 2893 | 3232 | 113 | ATT | CTATACTCA ATT A | T | A | TAA | **RNR2** | IDPMTWPTEQVTLGMTAQSYSRVHINNRVYDL DVGSGHPDGAAAIKGSFVQRLKSYVIWVQTGVI QVGFYLXSNSSLYERTREMRPTSQSAFPRKWYHL NLVLYPHPPKNRVC |

| 3289 | 4261 | 324 | ATT | AGAGGTTCA ATT C | T | C | TAA | **TRNL1** | IPLLNNMPMANLLLLIVPILIAMAFLMLTERKILGY MQLRKGPNVVGPYGLLQPFADAMKLFTKEPLKP ATSTITLYITAPTLALTIALLLWTPLPMPNPLVNLN LGLLFILATSSLAVYSILWSGWASNSNYALIGALRA VAQTISYEVTLAIILLSTLLMSGSFNLSTLITTQEHL WLLLPSWPLAMMWFISTLAETNRTPFDLAEGES ELVSGFNIEYAAGPFALFFMAEYTNIIMMNTLTTT IFLGTTYDALSPELYTTYFVTKTLLLTSLFLWIRTAY PRFRYDQLMHLLWKNFLPLTLALLMWYVSMPITI  SSIPPQT |
| --- | --- | --- | --- | --- | --- | --- | --- | --- | --- |
| 5425 | 5500 | 25 | ATA | AGTTTGAAC ATA A | A | A | TAA | **ND2** | MQNPPHSSPHSSPLPRYSYLSPLLY |
| 5857 | 5989 | 44 | ATT | TGTCTTTAG ATT A | T | A | TAG |  | IYSPMLHSAILPHPHWCSPTVDYSLQTTKTLEHYT  YYSAHELES |
| 7540 | 7738 | 66 | ATA | AACCATTTC ATA C | T | C | TAA | **TRND** | MTLSKLNYRLNPMYLNGTCSASRSTRRYFPYHRR  AYHLSWSRPHNHFPYLLPSPVCPFPNTHNKTN |
| 7894 | 8311 | 139 | ATG | TGGCCACCA ATG T | C | T | TAG | **COX2** | MVLNLRVHRLRRTNLQLLHTSPIIPRTRRPATPW RWQSSSTPDWSPHSYNNYITRRLALMSCPHIRLK NRCNSRTSKPNHFHRYTTGGMLRSMLWNLWSK PQFHAHRPRINSPKNLWNRARIYPMAPPLPPLEP  TVKLT |
| 8335 | 8506 | 57 | ATT | AAGTTAAAG ATT A | A | A | TAA | **TRNK** | IKRTNTSLQWNAPTKYYRMAHHNYPHTPYTIPH  HPTKNIKHKLPPTSLTKAHKNKKL |
| 8527 | 9205 | 226 | ATG | AGAACCAAA ATG A | A | A | TAA | **ATP8** | MNENLFASFIAPTILGLPAAVLIILFPPLLIPTSKYLI NNRLITTQQWLIKLTSKQMMTMHNTKGRTWSL MLVSLIIFIATTNLLGLLPHSFTPTTQLSMNLAMAI PLWAGTVIMGFRSKIKNALAHFLPQGTPTPLIPM LVIIETISLLIQPMALAVRLTANITAGHLLMHLIGSA TLAMSTINLPSTLIIFTILILLTILEIAVALIQAYVFTLL  VSLYLHDNT |
| 9721 | 9802 | 27 | ATT | TGGGTCTCT ATT T | T | T | TAG | **COX3** | ILPSYKPQSTSSLPSPFPTASTAQHFL |
| 9943 | 10003 | 20 | ATG | ATTTTGTAG ATG G | T | G | TAA | **COX3** | MWFDYFCMSPSIDEGLTLLV |
| 10789 | 10861 | 24 | ATT | AACAATTAT ATT C | T | C | TAG | **TRNC-comp** | ITTTDMTFQKTHNLNQHNHPQPNY |

| 11086 | 11410 | 108 | ATT | AATTATAAC ATT A | A | A | TAA | **ND4** | IHSHRTNHILYLLRNHTYPHLGYHHPMRQPARTP ERRHMLPILHPSRLPSPTHRTNLHSQHPRLTKHST THSHCPRTIKLLSQQLNMTSLHNSFYSKDTSLRTP  LMTP |
| --- | --- | --- | --- | --- | --- | --- | --- | --- | --- |
| 11557 | 11854 | 99 | ATG | ACTATCCCT ATG G | C | G | TAA | **ND4** | MRHNYNKLHLPTTNRPKIAHCMLFNQPHSPRSN SHSHPNPLKLHRRSHSHNRPRAYILITILPSKLKLRT  HSQSHHNPLSRTSNSTPTNSFLMTSSKPR |
| 12046 | 12139 | 31 | ATT | AAAACCCTC ATT A | C | A | TAA | **ND4** | IHTRKHPHVHTPIPHSPPIPQPRHHYRVFLL |
| 12310 | 14146 | 612 | ATT | GCCCCAAAA ATT T | A | T | TAA | **TRNL2** | ILVQLQMKVMTMHTTMTTLTLTSLIPPILTTLVNP NKKNSYPHYVKSIVASTFIISLFPTTMFMCLDQEVI ISNWHWATTQTTQLSLSFKLDYFSMMFIPVALFV TWSIMEFSLWYMNSDPNINQFFKYLLIFLITMLIL VTANNLFQLFIGWEGVGIMSFLLISWWYARADA NTAAIQAILYNRIGDIGFILALAWFILHSNSWDPQ QMALLNANPSLTPLLGLLLAAAGKSAQLGLHPW LPSAMEGPTPVSALLHSSTMVVAGIFLLIRFHPLA ENSPLIQTLTLCLGAITTLFAAVCALTQNDIKKIVAF STSSQLGLMMVTIGINQPHLAFLHICTHAFFKAM LFMCSGSIIHNLNNEQDIRKMGGLLKTMPLTSTS LTIGSLALAGMPFLTGFYSKDHIIETANMSYTNA WALSITLIATSLTSAYSTRMILLTLTGQPRFPTLTNI NENNPTLLNPIKRLAAGSLFAGFLITNNISPASPFQ TTIPLYLKLTALAVTFLGLLTALDLNYLTNKLKMKS PLCTFYFSNMLGFYPSITHRTIPYLGLLTSQNLPLLL LDLTWLEKLLPKTISQHQISTSIITSTQKGMIKLYFL  SFFFPLILTLLLIT |
| 14152 | 14218 | 22 | ATT | ACATAACCT ATT C | C | C | TAA |  | IPPSNLNYNMYTNKQCSTSNYY |
| 14230 | 14386 | 52 | ATA | TCAACGCCC ATA T | C | T | TAA |  | MIMQSPRTNRILPNQPWPLSFMNYSASYTIKVY  HNHHPIMLFHPQHQSYLHR |
| 14524 | 14584 | 20 | ATA | TAAACCCAT ATA C | C | C | TAA |  | MTSPKIQNNNTPDHTANNQY |
| 14665 | 14767 | 34 | ATA | AAACAAAGC ATA A | A | A | TAA |  | MHHYSRTDYNHDQWYEKPSLYFNYKNTNDPNT  QN |

| 14797 | 15394 | 199 | ATT | TAACCACTC ATT A | C | A | TAA | **CYTB** | IHRPPHPIQHLRMMKLRLTPWRLPDPPNHHRTI PSHALLTRRLNRLFINRPHHSRRKLWLNHPLPSR QWRLNILYLPLPTHRARPMLRIISLLRNLKHRHYP PACNYSNSLHRLCPPVRPNIILRGHSNYKLTIRHP MHWDRPSSMNLRRLLSRQSHPHTILYLSLHLALH  YCSPSNTPPPILARNGIKQPPRNHLPFR |
| --- | --- | --- | --- | --- | --- | --- | --- | --- | --- |
| 15469 | 15601 | 44 | ATT | CTTAATGAC ATT A | G | A | TAA | **CYTB** | INTILTRPPRRPRQLYPSQPLKHPSPHQARMMFP  IRLHNSPIRP |
| 15628 | 15895 | 89 | ATT | CCTTGCCCT ATT C | C | C | TAG | **CYTB** | ITIHPHPSNNPHPPYIQTTKHNISPTKPITLLTPSRR PPHSNLNRRTTSKLPFYHHWTSSIRTMLHNNPN  PNTNYLPNWKQNTQMGLSL |
| 15907 | 15997 | 30 | ATA | TATAAACTA ATA A | C | A | TAA | **TRNT** | MHQSCKPEMKTFFQGQIREKVFNSTISTQS |
| 16087 | 16315 | 76 | ATG | ACAACCGCT ATG A | G | A | TAA |  | MYFVHYCQPPWMLYGTMNTWPPVVHKNPIHI KTPSPCLQASTAINPQLSHINCNSKATPHPLGYQ  QTYPPLTVHST |
| 16333 | 16462 | 43 | ATA | TTACCGTAC ATA C | T | C | TAA |  | MAHYSQIPSRPHGWPPSDRGPLTTILREINIPHKS  ATLLAPGP |
| 116 | 209 | 31 | ATG | GAGCACCCT ATG C | C | C | TAA |  | MSQYLSLIPASSYYLSHLRSMLQANMLTKVC |
| 281 | 383 | 34 | ATA | ACAGACATC ATA C | A | C | TAA |  | MTKNFHQTPPPPLLATALKHISAKPQKQRTLTPA |
| 470 | 584 | 38 | ATA | CCCACTCCC ATA T | C | T | TAG |  | MLLISSMQPPPILPSTHTPLLTPYPEPTKPQRHPP  QFM |
| 695 | 794 | 33 | ATG | AGATTACAC ATG A | C | A | TAG | **RNR1** | MQASPFQWVHPLNHHDQKEQASSTQQCSSKRL  A |
| 860 | 968 | 36 | ATA | AACTAAGCT ATA T | G | T | TAA | **RNR1** | MLTPGLVNFVPATAVTRLTQVNRSRRKECFRSPP  PQ |
| 1751 | 1862 | 37 | ATA | GTATAGGCG ATA A | G | A | TAA | **RNR2** | MEIETWRNRYSTARERWKIMTKHNMARTNPYT  FCMMN |
| 1871 | 1934 | 21 | ATA | TAACTAGAA ATA C | G | C | TAA | **RNR2** | MTLQGEPKLRPPKPDELPKNS |
| 2192 | 2336 | 48 | ATT | CAGCCACCA ATT A | C | A | TAA | **RNR2** | IKKAFKLNTHYLKNPKHMTELLTPNWTNLSPYRR  TNVSMSNMKTFSSA |
| 2633 | 2705 | 24 | ATG | CCTGTATGA ATG C | T | C | TAA | **RNR2** | MAPRGFSCLLLLTSEIDLPVKRRA |
| 2798 | 2903 | 35 | ATT | CAAACCTGC ATT A | T | A | TAA | **RNR2** | IKNFGWGDLGAEPNLRAVHAKTSPVKANYYTQLI  Q |
| 2951 | 3050 | 33 | ATT | CGCAATCCT ATT T | C | T | TAA | **RNR2** | ILESMSTMGFTTSMLDQDIPMVQPLLKVRLFND |

| 3236 | 3347 | 37 | ATG | TTTGTTAAG ATG C | A | C | TAA | **TRNL1** | MAEPGNRMKLKTLQSEVQFLFLTTYPWPTSYSSL  YPF |
| --- | --- | --- | --- | --- | --- | --- | --- | --- | --- |
| 4217 | 4277 | 20 | ATG | TTATATGAT ATG C | G | C | TAA | **ND1** | MSPYPLQSPAFPLKPKKYVW |
| 4325 | 4418 | 31 | ATT | AACCCCCTT ATT C | C | C | TAA | **TRNI** | ISRTMRIEPIPENPKFSVPPITPHPKVRSAK |
| 4547 | 5063 | 172 | ATT | CTCGCACTG ATT T | C | T | TAA | **ND2** | IFYLSRPRNKHASFYSSSNQKNKPSFHRSCHQVFP HASNRIHNPSNSYPLQQYTLRTMNHNQYYQSM LIINNHNSYSNKTRNSPLSLLSPRGYPRHPSDIRPA SSHMTKTSPHLNHMPNLSLTKRKPSPHSLNLIHH SRQLRWIKPNPATQNLSMLLNYPHRMNNSSSTV  QP |
| 5114 | 5291 | 59 | ATT | TACTACCGC ATT C | C | C | TAG | **ND2** | IPTTQLKLQHHDPTTISHLKQANMTNTLNSIHPPL  PRRPAPANRLFAQMGHYRRIHKKQ |
| 5411 | 5519 | 36 | ATG | AAAAATAAA ATG C | A | C | TAG | **ND2** | MTVWTYKTHPIPPHTHRPYHATPTYLPFYTNNL  MEI |
| 5573 | 5642 | 23 | ATT | CAATACTTA ATT C | T | C | TAA | **TRNW** | ISVTAKDCKTPLCINWTQISHFN |
| 5807 | 6050 | 81 | ATG | CAATTCAAT ATG A | A | A | TAA |  | MKITSELVKRGLTPVFRFTVQCFTQPFYLTPTDVR RPLTILYKPQRHWNTMPIIRRMSWSPRHSSKPPY  SSRAGPARQPSR |
| 6089 | 6866 | 259 | ATT | AGCCCATGC ATT G | T | G | TAG | **COX1** | ICNNLLHSNTHHNRRLWQLTSSPNNRCPRYGVS PHKQHKLLTLTSLSPTPARICYSGGRSRNRLNSLPS LSRELLPPWSLRRPNHLLLTPSRCLLYLRGHQFHH NNYQYKTPCHNPMPNAPLRLIRPNHSSPTSPISP SPSCWHHYTTNRPQPQHHLLRPRRRRRPHSMP TPILIFRSPWSLYSYPTRLRNNLPYCNLLLRKKRTI WMHRYGLSYDINWLPRVYRVSTPYIYSRNRRRH  TSMFHLRYHNHRYPHRRQSI |
| 6902 | 7442 | 180 | ATG | CAATATGAA ATG T | G | T | TAG | **COX1** | MICCSALSPRIHLSFHRRWPDWHCISKLITRHRTT RHVLRCSPLPLCPINRSCICHHRRLHSLISPILRLHP RPNLRQNPFHYHIHRRKSNFLPTTLSRPIRNAPTL LGLPRCMHHMKHPIICRLIHFSNSSNINNFHDLRS LRFEAKSPNSRRTLHKPGVTMWMPPTLPHIRRT  RMHKI |

| 7586 | 8267 | 227 | ATG | TATATCTTA ATG C | T | C | TAG | **COX2** | MAHAAQVGLQDATSPIMEELITFHDHALMIIFLI CFLVLYALFLTLTTKLTNTNISDAQEMETVWTILP AIILVLIALPSLRILYMTDEVNDPSLTIKSIGHQWY WTYEYTDYGGLIFNSYMLPPLFLEPGDLRLLDVD NRVVLPIEAPIRMMITSQDVLHSWAVPTLGLKTD AIPGRLNQTTFTATRPGVYYGQCSEICGANHSFM  PIVLELIPLKIFEMGPVFTL |
| --- | --- | --- | --- | --- | --- | --- | --- | --- | --- |
| 8366 | 8570 | 68 | ATG | TACAGTGAA ATG C | G | C | TAG | **ATP8** | MPQLNTTVWPTMITPMLLTLFLITQLKMLNTNY HLPPSPKPMKMKNYNKPWEPKWTKICSLHSLPP  QS |
| 9251 | 9653 | 134 | ATG | ACCCAGCCC ATG C | C | C | TAG | **COX3** | MTPNRGPLSPPNDLRPSHVISLPLHNAPHTRPTN QHTNHMPMMARCNTRKHMPRPPHTTCPKRPS MRDNPIYYLRSFFLRRIFLSLLPLQPSPYPPIRRALA  PNRHHPAKSPRSPTPKHIRITRIRSINHLSSP |
| 9794 | 9998 | 68 | ATT | CGGCTCAAC ATT T | A | T | TAG | **COX3** | IFCSHRLPRTSRHYWLNFPHYLLHPPTNISLYIQTS  LWLRSRRLMLAFCRCGLTISVCLHLLMRVLLF |
| 10121 | 10304 | 61 | ATT | AATTATTAC ATT T | T | T | TAA | **ND3** | ILTTTTQRLHRKIHPLRVRLRPYIPRPRPFLHKILLSS  YYLLIIWSRNCPPFTPTMSPTNN |
| 10337 | 10412 | 25 | ATT | ATCCCTCTT ATT A | C | A | TAG | **ND3** | INHHPSPKSGLWVTTKRIRLNRIGM |
| 10502 | 10742 | 80 | ATT | TATACTAGC ATT A | A | A | TAA | **ND4L** | IYHLTSRNTSMSLTPHILPTMPRRNNTIAVHYSYS HNPQHPLPLSQYCAYCHTSLCRLRSSGGPSPTSL  NLQHMWPRLRT |

| 10760 | 12278 | 506 | ATG | CCTACTCCA ATG T | C | T | TAA | **TRNC-comp** | MLKLIVPTIMLLPLTWLSKKHMIWINTTTHSLIISII PLLFFNQINNNLFSCSPTFSSDPLTTPLLMLTTWLL PLTIMASQRHLSSEPLSRKKLYLSMLISLQISLIMTF TATELIMFYIFFETTLIPTLAIITRWGNQPERLNAG TYFLFYTLVGSLPLLIALIYTHNTLGSLNILLLTLTAQ ELSNSWANNLMWLAYTMAFMVKMPLYGLHL WLPKAHVEAPIAGSMVLAAVLLKLGGYGMMRL TLILNPLTKHMAYPFLVLSLWGMIMTSSICLRQTD LKSLIAYSSISHMALVVTAILIQTPWSFTGAVILMI AHGLTSSLLFCLANSNYERTHSRIMILSQGLQTLLP LMAFWWLLASLANLALPPTINLLGELSVLVTTFS WSNITLLLTGLNMLVTALYSLYMFTTTQWGSLTH HINNMKPSFTRENTLMFMHLSPILLLSLNPDIITG FSSCKYSLTKTSDCESDNRGLRPLIYRESSQELLTH  APMSNNMAFSTFKG |
| --- | --- | --- | --- | --- | --- | --- | --- | --- | --- |
| 12824 | 12893 | 23 | ATG | CCCGAGCAG ATG C | C | C | TAG | **ND5** | MPTQQPFKQSYTTVSAMSVSSSP |
| 14171 | 14231 | 20 | ATT | GCAATCTCA ATT C | T | C | TAA |  | ITMYTPTNNVQPVTTTNQRP |
| 14297 | 14498 | 67 | ATT | CCTTCATAA ATT T | T | T | TAA |  | IIQLPTLLKFTTTTTPSYSFTHSTNPTSIANPTKTLTK  TSTPDPHASGYSSMAIAVVYPKTTIIPPK |
| 14597 | 15905 | 436 | ATA | CCCCCATAA ATA G | T | G | TAA |  | MGEGLEENPTNPITKPTLNRNKAYIIILARTTTTTN DMKNHRCISTTRTPMTPMRKTNPLMKLINHSFI DLPTPSNISAWWNFGSLLGACLILQITTGLFLAM HYSPDASTAFSSIAHITRDVNYGWIIRYLHANGAS MFFICLFLHIGRGLYYGSFLYSETWNIGIILLLATM ATAFMGYVLPWGQMSFWGATVITNLLSAIPYIG TDLVQWIWGGYSVDSPTLTRFFTFHFILPFIIAALA TLHLLFLHETGSNNPLGITSHSDKITFHPYYTIKDAL GLLLFLLSLMTLTLFSPDLLGDPDNYTLANPLNTPP HIKPEWYFLFAYTILRSVPNKLGGVLALLLSILILAM IPILHMSKQQSMMFRPLSQSLYWLLAADLLILTW IGGQPVSYPFTIIGQVASVLYFTTILILMPTISLIENK  MLKWACPCSMN |
| 16016 | 16136 | 40 | ATT | ATTTAAACT ATT T | A | T | TAA |  | ILCSFMGKQIWVPPKYWLTHQQPLCISYITASHH  EYCTVP |

| 16160 | 16271 | 37 | ATA | CTGTAGTAC ATA A | T | A | TAG |  | MKTQSTSKPPPHAYKQVQQSTLNYHTSTATPKP  PLTH |
| --- | --- | --- | --- | --- | --- | --- | --- | --- | --- |
| 16340 | 16475 | 45 | ATT | ACATAGCAC ATT C | C | C | TAG |  | ITVKSLLVPMDDPPQMGVPWPPSSVKSMSRTR  VLLSSLRAHNTWG |
| 21 | 213 | 64 | ATT | TATCACCCT ATT A | C | A | TAA |  | INHSRELSMHLVFSSGGYARDSIARRWSRSTLCRS  ICLWFLPHPIIYRTYVQYYRRTYLLKCVN |
| 219 | 282 | 21 | ATG | AATTAATTA ATG T | T | T | TAA |  | MLVGHNNNNWMSAQPLSTQTS |
| 291 | 372 | 27 | ATT | TAACAAAAA ATT C | A | C | TAA |  | ISTKPPLPRFWPQHLNTSLPNPKNKEP |
| 390 | 477 | 29 | ATT | CCTAACCAG ATT C | C | C | TAA |  | ISNFIFWRYALLTVTPQLTHYFPLPLPYY |
| 924 | 1035 | 37 | ATA | CCCAAGTCA ATA A | T | A | TAA | **RNR1** | MEAGVKSVLDHPLPNKAKTHLSCKKLQLTQNRL  RKWL |
| 1218 | 1407 | 63 | ATA | CTGTAATCG ATA A | T | A | TAA | **RNR1** | MNPDQPHHLLLSLYTAIFSKPWWRLQSKRKYPR  KDVRSRCSPWGGKKWATFSTPENYDSPYET |
| 2052 | 2226 | 58 | ATT | CAACTTTAA ATT G | T | G | TAA | **RNR2** | ICPQNPLNPLVNLTVSPKRNSSLDTRKKPCRESKK  FNTHSRPKSSHQLRKRSSSTPTT |
| 2370 | 2445 | 25 | ATT | GAACTGACA ATT A | A | A | TAA | **RNR2** | INSPMSTINQQVIITLTVNPTQACS |
| 2598 | 2667 | 23 | ATA | AAAGGTAGC ATA T | A | T | TAA | **RNR2** | MITCSLNRDLYEWLHEGSAVSYF |
| 2805 | 2955 | 50 | ATT | GCATTAAAA ATT C | A | C | TAG | **RNR2** | ISVGATSEQNPTSEQYMLRLHQSKRTTMLNWSN  NLTNGTSYPRDNSAILF |
| 3048 | 3174 | 42 | ATT | TGTTCAACG ATT A | A | A | TAA | **RNR2** | IKVLRDLSSDRSNPGRFLSXFKFLPVRKDKRNKAY  FTKRLPP |
| 3177 | 3258 | 27 | ATG | CCCCCGTAA ATG T | T | T | TAA | **RNR2** | MMSSQLSIMPTPTQEQGLLRWQSPVIA |
| 3360 | 3633 | 91 | ATT | CGCAATGGC ATT C | G | C | TAG | **ND1** | IPNAYRTKNSRLYTTTQRPQRCRPLRATTTLRWR HKTLHQRAPKTRHIYHHPLHHRPDLSSHHRSSTM  NPPPHTQPPGQPQPRPPIYSSHL |
| 3825 | 4299 | 158 | ATT | ACACCTCTG ATT C | C | C | TAA | **ND1** | ITPAIMTLGHNMIYLHTSRDQPNPLRPCRRGVRT SLRLQHRMRRRPLRPILHSRMHKHYYNKHPHHY NLPRNNMWRTLPWTLHNMFCHQDPTSNLPVL MNSNSMPPIPLRPTHTPPMKKLPTTHPSITYMM  CLHTHYNLQHSPSNLRNMSDKRVTLME |

| 4470 | 5511 | 347 | ATT | CCCGTACTA ATT A | C | A | TAG | **ND2** | INPLAQPVIYSTIFAGTLITALSSHWFFTWVGLEM NMLAFIPVLTKKMNPRSTEAAIKYFLTQATASMIL LMAILFNNMLSGQWTMTNTTNQYSSLMIMMA MAMKLGMAPFHFWVPEVTQGTPLTSGLLLLTW QKLAPISIMYQISPSLNVSLLLTLSILSIMAGSWGG LNQTQLRKILAYSSITHMGWMMAVLPYNPNMT ILNLTIYIILTTTAFLLLNLNSSTTTLLLSRTWNKLTW LTPLIPSTLLSLGGLPPLTGFLPKWAIIEEFTKNNSLI IPTIMATITLLNLYFYLRLIYSTSITLLPMSNNVKMK  WQFEHTKPTPFLPTLIALTTLLLPISPFMLMIL |
| --- | --- | --- | --- | --- | --- | --- | --- | --- | --- |
| 5904 | 7563 | 553 | ATG | CCCCCACTG ATG T | C | T | TAG | **COX1** | MFADRWLFSTNHKDIGTLYLLFGAWAGVLGTAL SLLIRAELGQPGNLLGNDHIYNVIVTAHAFVMIFF MVMPIMIGGFGNWLVPLMIGAPDMAFPRMN NMSFWLLPPSLLLLLASAMVEAGAGTGWTVYPP LAGNYSHPGASVDLTIFSLHLAGVSSILGAINFITTI INMKPPAMTQYQTPLFVWSVLITAVLLLLSLPVLA AGITMLLTDRNLNTTFFDPAGGGDPILYQHLFWF FGHPEVYILILPGFGMISHIVTYYSGKKEPFGYMG MVWAMMSIGFLGFIVWAHHMFTVGMDVDTR AYFTSATMIIAIPTGVKVFSWLATLHGSNMKWSA AVLWALGFIFLFTVGGLTGIVLANSSLDIVLHDTYY VVAHFHYVLSMGAVFAIMGGFIHWFPLFSGYTL DQTYAKIHFTIMFIGVNLTFFPQHFLGLSGMPRR YSDYPDAYTTWNILSSVGSFISLTAVMLMIFMIW EAFASKRKVLMVEEPSMNLEWLYGCPPPYHTFE EPVYMKSRQKRKESNPPKLVSSQPHGLHDFFKKV  LEKPFHNFVKVKL |
| 8502 | 8715 | 71 | ATT | AAATAAAAA ATT T | A | T | TAA | **ATP8** | IMTNPENQNERKSVRFIHCPHNPRPTRRSTDHSI SPSIDPHLQMSHQQPTNHHPTMTNQTNLKTND  NHTQH |
| 8799 | 8883 | 28 | ATT | GCCTCACTC ATT A | C | A | TAA | **ATP6** | IYTNHPTIYKPSHGHPLMSGHSDYRLSL |

| 9204 | 10764 | 520 | ATA | CGACAACAC ATA T | C | T | TAA | **ATP6** | MMTHQSHAYHMVKPSPWPLTGALSALLMTSGL AMWFHFHSMTLLMLGLLTNTLTMYQWWRDV TRESTYQGHHTPPVQKGLRYGMILFITSEVFFFAG FFWAFYHSSLAPTPQLGGHWPPTGITPLNPLEVP LLNTSVLLASGVSITWAHHSLMENNRNQMIQAL LITILLGLYFTLLQASEYFESPFTISDGIYGSTFFVAT GFHGLHVIIGSTFLTICFIRQLMFHFTSKHHFGFEA AAWYWHFVDVVWLFLYVSIYWWGSYSFSMNS TVNFQLTSFDNIQKRVMNFALILMINTLLALLLMII TFWLPQLNGYMEKSTPYECGFDPMSPARVPFS MKFFLVAITFLLFDLEIALLLPLPWALQTTNLPLMV MSSLLLIIILALSLAYEWLQKGLDWTELVYSLNKTN DFDSLNYDNHIYQMPLIYMNIMLAFTISLLGMLV YRSHLMSSLLCLEGMMLSLFIMATLMTLNTHSLL ANIVPIAMLVFAACEAAVGLALLVSISNTYGLDYV  HNLNLLQC |
| --- | --- | --- | --- | --- | --- | --- | --- | --- | --- |
| 11115 | 11220 | 35 | ATA | TCATATTTT ATA C | T | C | TAG | **ND4** | MSSSKPHLSPPWLSSPDEATSQNAWTQAHTSYS  TP |
| 12165 | 12228 | 21 | ATT | AAACATCAG ATT T | C | T | TAA | **TRNH** | IVNLTTEAYDPLFTEKAHKNC |
| 12234 | 12408 | 58 | ATG | TGCTAACTC ATG C | C | C | TAA | **TRNQ-comp** | MPPCLTTWLSQLLKDNSYPLVLGPKNFGATPNKS  NNHAHYYNHPNPDFPNSPHPYHPR |
| 12429 | 12678 | 83 | ATA | AAAAAACTC ATA C | C | C | TAA | **ND5** | MPPLCKIHCRIHLYYQSLPHNNIHVPRPRSYYLELT LSHNPNNPALPKLQTRLLLHNIHPCSIVRYMVHH  RILTVMYKLRPKH |
| 12747 | 12954 | 69 | ATT | TAACAACCT ATT C | C | C | TAA | **ND5** | IPTVHRLRGRRNYILLAHQLMMRPSRCQHSSHSS NPMQPYRRYRFHPRLSMIYPTLQLMRPTTNSPS  KR |
| 13014 | 13143 | 43 | ATT | ATCAGCCCA ATT G | C | G | TAG | **ND5** | IRSPPLTPLSHRRPHPSLSPTPLKHYSCSRNLLTHPL  PPPSRK |
| 13170 | 13386 | 72 | ATG | TCTAACACT ATG T | A | T | TAA | **ND5** | MLRRYHHSVRSSLRPYTKWHQKNRSLLHFKSTRT HNSYNRHQPTTPSIPAHLYPRLLQSHTIYVLRVHH  PQP |
| 13473 | 13602 | 43 | ATT | CAGCCTAGC ATT G | A | G | TAG | **ND5** | ISRNTFPHRFLLQRPHHRNRKHIMHKRLSPIYYSH  RYLPDKRL |
| 14145 | 14208 | 21 | ATA | CCTAATCAC ATA C | C | C | TAA | **ND5** | MTYSPEQSQLQYMHQQTMFNQ |

| 14703 | 14775 | 24 | ATG | CCACGACCA ATG T | C | T | TAA |  | MMWKTIVVFQLQEHQWPQYAKLTP |
| --- | --- | --- | --- | --- | --- | --- | --- | --- | --- |
| 14967 | 15117 | 50 | ATT | GAGACGTAA ATT T | T | T | TAG | **CYTB** | IMAESSATFTPMAPQYSLSASSYTSGEAYITDHFS  TQKPETSALSSCLQL |
| 15387 | 15462 | 25 | ATT | TCACCTCCC ATT C | C | C | TAA | **CYTB** | IPMKSPSTLTTQSKTPSAYFSSFSP |
| 16062 | 16161 | 33 | ATT | CACCCAAGT ATT A | A | A | TAA |  | IDSPINNRYVFRTLLPATMNIVRYHKYLTTCST |
| 16194 | 16308 | 38 | ATG | CCCCTCCCC ATG T | C | T | TAG |  | MLTSKYSNQPSTITHQLQLQSHPSPTRMPTNLPT  LNST |
| 16314 | 16386 | 24 | ATA | ACATAGTAC ATA A | T | A | TAG |  | MKPFTVHSTLQSNPFSSPWMTPLR |
| -16315 | -16087 | 76 | ATG | AATGGCTTT ATG A | T | A | TAG |  | MYYVLLRVGRFVGILVGEGWLWSCSWCVMVEG WLLYLLVSMGRGFWCGLGFYVLQVVKYLWYRT  MFMVAGSNVRNT |
| -16075 | -16006 | 23 | ATG | CGGTTGTTG ATG G | T | G | TAG |  | MGESMLGWYPNLLPHERTENSLN |
| -15139 | -15052 | 29 | ATA | CGGGAGGAC ATA C | G | C | TAG | **CYTB-comp** | MAYEGCCYSCKQEDNADVSGFWVEKWSVM |
| -15001 | -14899 | 34 | ATT | TGAGGCGCC ATT G | G | G | TAG | **CYTB-comp** | IGVKVADDSAMIYVSSDVGDWWKGGWGVWW  VVHG |
| -14719 | -14611 | 36 | ATG | AATACAACG ATG T | A | T | TAA |  | MVFHIIGRGCSPCENNDVCFVSVECGFSNGVCG  VFF |
| -14470 | -14320 | 50 | ATA | TGTCTTTGG ATA A | T | A | TAA |  | MYYSDGYWGVSWGMGVRGWGLGECFSGVSD  GGRIGAVGERVWWGGGCGKL |
| -14221 | -14152 | 23 | ATT | TGGGCGTTG ATT G | T | G | TAG |  | ISSSYWLNIVCWCMYCNWDCSGE |
| -13564 | -13387 | 59 | ATA | GAGTAATAG ATA G | T | G | TAA | **ND5-comp** | MGLRRLCMMCLRFRWCGLWSRNLWGKVFLLM  LGCQWWGRLKWEVWFWVVLLFFEYLVHC |
| -13354 | -13282 | 24 | ATA | AGCACATAA ATA T | T | T | TAA | **ND5-comp** | MVWLWRRRGYRCAGMLGVVGWCRL |
| -13171 | -13090 | 27 | ATA | CGCCTAAGC ATA T | A | T | TAG | **ND5-comp** | MVLEFGLVGYFLLGGGSGWVRRFLLQL |
| -13015 | -12955 | 20 | ATT | GGAGACCTA ATT G | C | G | TAG | **ND5-comp** | IGLICLLLLGGGLVVGWGLD |
| -12898 | -12784 | 38 | ATG | GGATAAATC ATG T | A | T | TAA | **ND5-comp** | MLRRGWNRYRRYGCMGLLEWLLCWHLLGRIIN  WWARRM |
| -12616 | -12523 | 31 | ATG | TAACGAACA ATG T | A | T | TAA | **ND5-comp** | MLQGWMLWRSSLVWSLGRAGLFGLWLSVSSR |
| -10711 | -10483 | 76 | ATT | GTGTTGGAG ATT A | G | A | TAA | **TRNY** | IETSRARPTAASQAAKTSMAMGTMLAKREWVL RVMRVAMMNSDSIIPSRHSREDMRCERYTSIPR  SEMVNASMMFM |
| -10348 | -10168 | 60 | ATG | AGGGCTAGG ATG T | A | T | TAA | **ND3-comp** | MMINKRDDMTISGRLVVCRAHGRGKRRAISRSN  NKKVMATKKNFMEKGTRAGDMGSKPHS |

| -10117 | -9991 | 42 | ATA | CAAAATGTA ATA T | G | T | TAA | **ND3-comp** | MIISSKARRVLIIKIKAKFITLFWMLSKLVNWKLTV  LFMLKE |
| --- | --- | --- | --- | --- | --- | --- | --- | --- | --- |
| -9973 | -9784 | 63 | ATG | CATCAATAG ATG A | T | A | TAG | **COX3-comp** | METYRNSQTTSTKCQYQAAASKPKWCLDVKWN ISWRMKQMVRKVEPMMTWSPWKPVATKNVE  P |
| -9781 | -9721 | 20 | ATG | GAGCCGTAG ATG C | T | C | TAG | **COX3-comp** | MPSEMVKGDSKYSEACRRVK |
| -9706 | -9511 | 65 | ATT | CCCAGTAAA ATT T | A | T | TAA | **COX3-comp** | IVMSSAWIIWFRLFSIRLWWAQVIDTPDASNTD VFRSGTSRGFSGVMPVGGQCPPNWGVGARLE  W |
| -9466 | -9229 | 79 | ATA | TCTGAGGTA ATA A | G | A | TAG | **COX3-comp** | MNRIIPYRRPFWTGGVWWPWYVLSRVTSRHH WYMVSVLVSRPSMRSVMEWKWNHMARPEVI  RRAERAPVRGHGLGFTMW |
| -9208 | -9127 | 27 | ATT | TGGTGGGTC ATT T | G | T | TAG | **COX3-comp** | IMCCRAGRGLLEVWKRRLGLRRQRFLG |
| -8668 | -8470 | 66 | ATT | TGATTAGTC ATT T | G | T | TAG | **ATP6-comp** | IVGWWLVGCWWDIWRWGSMEGEMEWSVLR RVGLGLWGQWMKRTDFRSFWFSGFVMIFYFYG  LWWGR |
| -7372 | -7231 | 47 | ATG | TCCAGGTTT ATG A | T | A | TAG | **COX1-comp** | MEGSSTIRTFRFEAKASQIMKIINITAVREMNEPT  DDRMFHVVYASG |
| -7210 | -7129 | 27 | ATT | CGTCGGGGC ATT C | G | C | TAG | **COX1-comp** | IPDRPRKCCGKKVRFTPMNMMVKWILA |
| -6997 | -6685 | 104 | ATG | TGTAGTACG ATG C | A | C | TAG | **COX1-comp** | MSSDEFANTMPVRPPTVKRKMNPRAQSTAADH FMLLPWSVASQLNTLTPVGMAMIMVAEVKYAR VSTSIPTVNMWCAHTMNPRKPIDIMAQTMPM  YPNGSFFPE |
| -6589 | -6310 | 93 | ATG | TGGTATAGA ATG G | A | G | TAG | **COX1-comp** | MGSPPPAGSKKVVLRLRSVSSMVMPAARTGRD RRSRTAVIRTDQTKRGVWYWVMAGGFMLMIV  VMKLMAPKMEETPARCKEKMVRSTEAPGWE |
| -6253 | -6064 | 63 | ATA | GCCTCCACT ATA C | A | C | TAG | **COX1-comp** | MADASRSRREGGKSQKLMLFMRGNAMSGAPII  RGTSQLPKPPIMMGITMKKIITNAWAVTMTL |
| -6061 | -5842 | 73 | ATG | ACGTTGTAG ATG G | T | G | TAG | **COX1-comp** | MWSLPRRLPGWPSSARMRRLRAVPRTPAHAPN NRYSVPMSLWFVENSQRSANISGGEVKWLSEAL  DCKSKDRG |
| -5806 | -5710 | 32 | ATT | GATTTTCAT ATT A | C | A | TAG |  | IELQIRRSSFKPAGASPAFFPGGGRSRLKPVD |
| -5641 | -5572 | 23 | ATT | CTTAGCTTA ATT A | T | A | TAA |  | IKVADLRSVDAEWGFAVLSCYRN |

| -5506 | -5362 | 48 | ATT | TTCTATAAG ATT T | A | T | TAG | **ND2-comp** | IISMKGEMGRSSVVRAMSVGRNGVGFVCSNCH  FIFTLLDMGSSVIEVE |
| --- | --- | --- | --- | --- | --- | --- | --- | --- | --- |
| -5323 | -5092 | 77 | ATG | AGGAGGGTG ATG T | G | T | TAA | **ND2-comp** | MVAMMVGMMRLLFFVNSSMMAHLGKKPVSG GRPPRERRVDGIKGVSHVSLFQVRDSSRVVVLEF  KLSSRNAVVVRMM |
| -5089 | -4498 | 197 | ATA | ATAATATAA ATA T | T | T | TAG | **ND2-comp** | MVKLRMVMLGLYGRTAIIHPMWVIEEYAKILRS WVWFNPPQLPAMMDKIERVRRRLTFSEGEIWY MIEMGASFCHVRRSRPDVRGVPWVTSGTQKW KGAIPSFIAMAIMIINDEYWLVVLVMVHCPESML LKRMAIRRIMDAVACVRKYLMAASVERGFIFLVR TGMKASMFISRPTQVKNQCELSAVMSVPAKMV  E |
| -4396 | -4315 | 27 | ATG | TACTTTAGG ATG G | A | G | TAA |  | MGCDRWHGEFWILRDGFDSHSPRNKGV |
| -4306 | -4234 | 24 | ATT | TAAGCTCCT ATT T | C | T | TAA | **TRNI-comp** | IIYSIKVTLLSDMFLRFEGECWRL |
| -4204 | -4024 | 60 | ATG | ATATAAGTA ATG T | G | T | TAG | **ND1-comp** | MLGWVVGSFFMGGVWVGRSGIGGMLFEFMR  TGRLEVGSWWQNMLCRVQGRVRHMLFLGRL |
| -3988 | -3865 | 41 | ATT | TGTTTGTGT ATT G | T | G | TAA | **ND1-comp** | IRLWRMGRRGLRRIRCWSLRLVRTPLRQGRRGF  GWSLLVWR |
| -3766 | -3634 | 44 | ATG | TTATTAGTA ATG T | G | T | TAG | **ND1-comp** | MLMVEWWLGWLHMRLFGLLLAVRRSGRSLSL  MLTLIRGLSKRLG |
| -3616 | -3505 | 37 | ATA | CTAGAATAA ATA G | T | G | TAG | **ND1-comp** | MGGLGWGWPGGWVWGGGFMVEERWWELR  SGRWCRGWW |
| -3502 | -3373 | 43 | ATG | TGATGGTAG ATG G | T | G | TAA | **ND1-comp** | MWRVLGALWWRVLWRQRRVVVARRGLQRW  GLCVVVYSLEFFVR |
| -3361 | -3301 | 20 | ATG | GCATTAGGA ATG C | G | C | TAA | **ND1-comp** | MPLRLEWVQWGVGGWPWVCC |
| -3253 | -3151 | 34 | ATT | TTTTATGCG ATT C | G | C | TAG | **TRNL1-comp** | ITGLCHLNKPCSWVGVGMMLSWDDIIYGGRRFV  K |
| -2887 | -2782 | 35 | ATA | CAATTGAGT ATA T | A | T | TAG | **RNR2-comp** | MVVRFDWWSLSMYCSEVGFCSEVAPTEIFNAGL  VV |
| -2731 | -2668 | 21 | ATA | TAAAGCTCC ATA G | T | G | TAA | **RNR2-comp** | MGSSRLAVLCPPLHGQVNFTG |
| -1861 | -1705 | 52 | ATT | TTCTAGTTA ATT A | T | A | TAG | **RNR2-comp** | IHYAEGMGVSPCYIMLGYNFSSFPCGTMSIAPGF  NFYRLYFIWVNGLAKVVW |
| -1267 | -1159 | 36 | ATG | TTGCTGAAG ATG C | A | C | TAA | **RNR1-comp** | MAVYRLSKRWWGWSGFIDYRTGSSRGMWSTA  RSFEF |

| -673 | -532 | 47 | ATA | TAAGAGCTA ATA A | C | A | TAG | **RNR1-comp** | MERLGPNLFVYGVMWARLNIFSVLLWGGKLHKL  WGVSLGFGWFGVWG |
| --- | --- | --- | --- | --- | --- | --- | --- | --- | --- |
| -292 | -214 | 26 | ATT | TTGGTGGAA ATT T | G | T | TAA |  | IFCYDVCVESGCADIQLLLLCPTSIN |
| -157 | -13 | 48 | ATA | GTGCGATAA ATA T | T | T | TAG |  | MMGWGRNQRQMLRHRVLRLQRLAMLSRAYP  PDENTKCMESSREWLMGW |
| -16500 | -16326 | 58 | ATG | AGGAACCAG ATG C | C | C | TAA |  | MSDTVHFSYPQVLWARSEESSTLVRDIDFTEDGG  QGTPIWGGSSMGTRRDLTVMCYVR |
| -16161 | -16011 | 50 | ATG | TGGGTTTTT ATG A | T | A | TAA |  | MYYRWSSIYGTVQYSWWLAVMYEMHSGCWW  VSQYLGGTQICFPMKEQRMV |
| -16008 | -15906 | 34 | ATT | ATAGTTTAA ATT G | T | G | TAG |  | IRILALGANGGVKDFFSDLSLEKGFHLRFTRLVY |
| -15900 | -15780 | 40 | ATA | TATTAGTTT ATA T | T | T | TAG | **TRNT-comp** | MLQGQAHLSILFSIREMVGIRIRIVVKYSTDATCP  MMVKG |
| -15693 | -15579 | 38 | ATT | GGGCGAAAT ATT T | A | T | TAG | **CYTB-comp** | IMLCCLDMWRMGIIARMRMDSNRARTPPSLLG  TDRRIV |
| -15399 | -15249 | 50 | ATT | TGGAAGGTG ATT T | G | T | TAG | **CYTB-comp** | ILSEWEVIPRGLFDPVSCKNRRWSVARAAMMKG  KMKWKVKNRVRVGLSTE |
| -15237 | -15138 | 33 | ATT | CCTCCTCAG ATT A | C | A | TAG | **CYTB-comp** | IHWTRSVPMYGMADSKFVITVAPQNDIWPHGR  T |
| -15132 | -15072 | 20 | ATG | ACATAGCCT ATG A | C | A | TAG | **CYTB-comp** | MKAVAMVASRRMMPMFQVSE |
| -14904 | -14139 | 255 | ATG | GAGTAGTGC ATG C | T | C | TAG | **CYTB-comp** | MARNSPVVIWRIRQAPRSEPKFHHAEMLDGVG RSMNEWLINFIRGLVLRIGVIGVLVVEMQRWFF MSLVVVVVRARMMMYALFLLSVGLVMGFVGFS SKPSPIYGGLVLIVSGVVGCVIILNFGGGYMGLMV FLIYLGGMMVVFGYTTAMAIEEYPEAWGSGVEV LVSVLVGLAMEVGLVLWVKEYDGVVVVVNFNSV GSWMIYEGEGSGLIREDPIGAGALYDYGRWLVV  VTGWTLFVGVYIVIEIARGNRLCD |
| -13953 | -13887 | 22 | ATA | AAGGCCTAG ATA G | T | G | TAG | **ND5-comp** | MGDCAVCDARVESEYVGEMKCA |
| -12696 | -12582 | 38 | ATA | GATGAGTAG ATA T | T | T | TAG | **ND5-comp** | MFEELINVWVWVYMSQWEFYDGPCNEQCYRD  EYYGEVV |
| -12441 | -12381 | 20 | ATA | GGATTTTAC ATA T | T | T | TAG | **ND5-comp** | MMGVWVFFVRVNEGGKDGGN |
| -12291 | -12204 | 29 | ATG | CTAAGACCA ATG A | C | A | TAA | **TRNL2-comp** | MDSCYPLKVEKAMLLDMGAWVSSSCELSR |

| -12201 | -11985 | 72 | ATA | TCTCGGTAA ATA G | T | G | TAG | **TRNQ** | MRGRKPLLSDSQSDVLVKLYLQEENPVMMSGLR DRRRMGDRCMNMRVFSRVNEGFMLLMWWV  SEPHCVVVNM |
| --- | --- | --- | --- | --- | --- | --- | --- | --- | --- |
| -11958 | -11760 | 66 | ATG | GTGACTAGT ATG T | A | T | TAG | **ND4-comp** | MLSPVSRRVMFDQENVVTSTESSPSRLMVGGKA RLARLARSHQKAISGSRVWSPWERIMMRLWVR  S |
| -11709 | -11535 | 58 | ATT | CCGTGGGCG ATT T | G | T | TAG | **ND4-comp** | IMRMTAPVKLQGVWMRMAVTTRAMWLIEEYA  MSDFRSVCRRQMELVMIMPHRDSTRKG |
| -11343 | -11256 | 29 | ATT | GCTAGTCAT ATT A | C | A | TAA | **ND4-comp** | IKLLAQEFDSSWAVRVSSRMFSEPRVLWV |
| -11109 | -11043 | 22 | ATG | ATATAAAAT ATG T | A | T | TAG | **ND4-comp** | MISSVAVNVMIKEICREISMER |
| -10992 | -10650 | 114 | ATG | TGGCTTGCC ATG T | G | T | TAG | **ND4-comp** | MIVRGRSQVVSIRRGVVRGSEEKVGEQLNRLLLI WLKNSRGMMLMIRLWVVVLIQIMCFLESHVSG SNMIVGTISFSIGVGLGYVRSLGHMCWRLRLVGL  GPPLLRRRQRLVWQ |
| -10584 | -10506 | 26 | ATA | TGAACAGCG ATA T | G | T | TAA | **ND4L-comp** | MVLFLLGMVGRMWGVSDMLVFLEVRW |
| -10443 | -10323 | 40 | ATG | CATAATTTA ATG G | T | G | TAA | **TRNR-comp** | MSRNHSFCLNYMPIRFSLILFVVTHRPDLGLGW  WLMRGMT |
| -9957 | -9888 | 23 | ATA | CATACAGAA ATA T | G | T | TAA | **COX3-comp** | MVKPHLQNASIRRRLRSQSDVWM |
| -9795 | -9726 | 23 | ATG | CTACAAAAA ATG T | A | T | TAA | **COX3-comp** | MLSRRCRRKWWRETRSTLRLVGG |
| -9633 | -9531 | 34 | ATA | AGGTGATTG ATA T | T | T | TAG | **COX3-comp** | MLLMRVMRMCLGVGLLGDLAGWCLLGASALLI  GG |
| -9462 | -9177 | 95 | ATA | AGGTAATAA ATA G | T | G | TAG | **COX3-comp** | MGLSRIEGLFGQVVCGGLGMCFLVLHRAIIGMW LVCWLVGLVWGALWSGSEITWLGRRSLGGLRG  PLLGVMGWVLLYDRHVIGGSLCVVVQVEAY |
| -8448 | -8358 | 30 | ATT | GTGTTTAAT ATT T | A | T | TAA | **ATP8-comp** | IFSWVMRNSVRSMGVIMVGHTVVFSWGISL |
| -8247 | -8163 | 28 | ATT | ACGGGCCCT ATT C | C | C | TAG | **COX2-comp** | ISKIFRGINSRTMGMKLWFAPQISEHWP |
| -8109 | -7947 | 54 | ATT | CGTCCGGGA ATT C | G | C | TAG | **COX2-comp** | IASVFKPNVGTAHECKTSCDVIIMRMGASIGSTTR  LSTSRSRRSPGSRNNGGSM |
| -7833 | -7569 | 88 | ATG | ATGTAAAGG ATG G | A | G | TAG | **COX2-comp** | MRRDGRAMRTRMMAGRMVQTVSISWASEML VLVSFVVSVRKRAYRTRKQMRKMIMRAWSWKV  MSSSMMGEVASCRPTCAACAIKMYRI |
| -7536 | -7443 | 31 | ATG | AGTTATGAA ATG T | G | T | TAG | **TRND-comp** | MVFLMPFWKSHGGHGVGLKPALGGSIPSFFV |

| -7440 | -7149 | 97 | ATT | TTTGTCTAG ATT T | T | T | TAG | **COX1-comp** | ILCMRVLRMCGRVGGIHMVTPGLWRVLLLLGLF ASKRRLLKSWKLLMLLLLEKWMSLQMMGCFM WCMHRGSPSNVGAFRMGRESVVGRKLDLRRW  MW |
| --- | --- | --- | --- | --- | --- | --- | --- | --- | --- |
| -7101 | -6867 | 78 | ATA | AGCCTGAGA ATA G | A | G | TAA | **COX1-comp** | MGEISEWSLLWWQMQLLLMGHSGSGLQRSTC RVVRCLVMSLLMQCQSGHLRWKERWILGLRAL  QQIISYCFRGVWRVS |
| -6705 | -6642 | 21 | ATG | TGTATCCAA ATG T | C | T | TAA | **COX1-comp** | MVLFFRSSKLQYGRLFRSLVG |
| -6606 | -6525 | 27 | ATA | AAAATCAGA ATA G | A | G | TAG | **COX1-comp** | MGVGMEWGLLLRRGRRRWCWGCGLLVV |
| -6495 | -6408 | 29 | ATA | CTGGGAGAG ATA G | G | G | TAA | **COX1-comp** | MGEVGLLWLGRIRRRGAFGIGLWQGVLYW |
| -6246 | -6072 | 58 | ATG | CTATAGCAG ATG G | C | G | TAA | **COX1-comp** | MRAGVGEREVRVRSLCCLCGETPYRGHRLLGELV  SCQSLRLWWVLLWRRLLQMHGLWR |
| -5814 | -5694 | 40 | ATT | TCCGAGGTG ATT T | G | T | TAA |  | IFMLNCKFEEAASNLPGLLPPFFPAAGEVDWSQL  IRVLSC |
| -5628 | -5520 | 36 | ATT | AAGTGGCTG ATT G | C | G | TAA |  | ICVQLMQSGVLQSLAVTEIKYCNLLRALKALGLYL  T |
| -5259 | -5109 | 50 | ATT | TAATGGCCC ATT G | C | G | TAG | **ND2-comp** | IWAKSRLAGAGLLGRGGWMELRVLVMLACFRC  EMVVGSWCWSLSWVVGMR |
| -5010 | -4824 | 62 | ATG | TTGAGGAGT ATG T | A | T | TAA | **ND2-comp** | MLRFCVAGFGLIHLNCLLWWMRLREWGEGLRL  VRERFGMWLRWGLVFVMWEEAGRMSEGCLG |
| -4668 | -4590 | 26 | ATG | GGATTATGG ATG G | T | G | TAA | **ND2-comp** | MRLLAWGNTWWQLLWNEGLFFWLELE |
| -4434 | -4287 | 49 | ATG | TTTCGGGGT ATG G | G | G | TAA | **TRNM-comp** | MGPMAYLADLTLGWGVMGGTENFGFSGMGSI  LMVLEMRGFKLLLFTLSK |
| -3732 | -3639 | 31 | ATA | GGTGACTTC ATA G | T | G | TAG | **ND1-comp** | MWDCLGYCSQCADQGVVWVWCSPWSEDWV  NG |
| -3186 | -3105 | 27 | ATG | TAAGTTGAG ATG T | G | T | TAG | **RNR2-comp** | MMSFTGEGALWSRPYFSCPFVQGGIWX |
| -3006 | -2931 | 25 | ATG | ACCATCGGG ATG C | G | C | TAG | **RNR2-comp** | MSWSNIEVVNPIVDMDSRMGLRCYP |
| -2679 | -2487 | 64 | ATT | GGCAGGTCA ATT C | T | C | TAA | **RNR2-comp** | ISLVKSKRQLNPRGAIHTGPYLRNKWLCYLCTVRV  PRPLNMCHWAGGASNTGDARGDVFGKQAG |
| -1824 | -1731 | 31 | ATA | GTCCTTGCT ATA T | G | T | TAA | **RNR2-comp** | MLCLVMIFHLSLAVLYLLRQVSISIAYTLFG |
| -1506 | -1374 | 44 | ATA | CTTTGAAGT ATA T | A | T | TAG | **RNR1-comp** | MLEEGDGRCVRASGPCSTKHSTLSLLLNPPSTLKF  HKGYRSFLG |
| -1257 | -1116 | 47 | ATA | TGGCGGTAT ATA G | T | G | TAA | **RNR1-comp** | MGWARGGEVDRGLSITEQAPLEGYEAPPGPLSF  KLWLVVFWRAVLLI |

| -582 | -402 | 60 | ATA | GTAAGCTAC ATA A | T | A | TAA | **TRNF-comp** | MNCGGCLWGLVGSGYGVSSGVCVLGRMGGGC  IDEISSMGVGGENNVLVGGWLLKVHTAKR |
| --- | --- | --- | --- | --- | --- | --- | --- | --- | --- |
| -342 | -210 | 44 | ATG | TTGGCAGAG ATG G | G | G | TAA |  | MCLSAVARSGGGGVWWKFFVMMSVWKVAVQ  TFNCYYYVLQALIN |
| -117 | -24 | 31 | ATA | TACTGCGAC ATA G | G | G | TAA |  | MGCSGSSVSQCYRVHTPQTKMPNAWRAPVSG |
| -16562 | -16343 | 73 | ATG | TCCATCGTG ATG C | G | C | TAA |  | MSYLRGTCGLFRLYDPEVGTRCRMQFTLATPKCY GPGARRVALLCGMLISRRMVVKGPLSEGGHPW  GREGIWL |
| -16238 | -16103 | 45 | ATG | TTGCAGTTG ATG G | T | G | TAA |  | MCDSWGLIAVLACKHGEGVLMWIGFLCTTGGQ  VFMVPYNIHGGWQ |
| -16088 | -16016 | 24 | ATA | ACGAAATAC ATA C | T | C | TAG |  | MAVVDGWVNTWVVPKSASPWKNRE |
| -15983 | -15851 | 44 | ATG | TGGGTGCTA ATG T | C | T | TAG |  | MVELKTFSLICPWKKVFISGLQDWCISLYYKDRPI  WVFCFQLGR |
| -15713 | -15602 | 37 | ATT | AATAAAGTG ATT G | G | G | TAG | **CYTB-comp** | IGLVGEMLCFVVWMYGGWGLLLGWGWMVM  GQGRLLVC |
| -15572 | -15311 | 87 | ATA | TGTAGGCGA ATA G | C | G | TAA | **CYTB-comp** | MGNIIRAWCGEGCLRGWLGYNCLGRLGGLVRM VLMSLRREGREVSRGRLWLCSKGGRWFYRNGR  WFLGGCLIPFRARMGGGVLLGLQ |
| -15014 | -14789 | 75 | ATA | AGATAAAGA ATA T | A | T | TAA | **CYTB-comp** | MLRRHWREGSGWFSHNLRLEWCGRLMKRRLR RLVSSAWLGMVLWWFGGSGRRQGVSRSFIMR  RCWMGWGGRWMSG |
| -14468 | -14396 | 24 | ATA | TCTTTGGAT ATA T | G | T | TAG |  | MLQRWLLRSILRHGGQGLRSWWVF |
| -14294 | -14219 | 25 | ATG | GAATAATTT ATG A | T | A | TAG |  | MKERGQGWFGRILLVRGLCMIMGVD |
| -14087 | -13952 | 45 | ATT | TAAAGTTTA ATT T | T | T | TAG | **ND5-comp** | IMPFWVEVMMEVEIWCCEIVLGNSFSSQVRSRR  SRGRFWLVRRPR |
| -13781 | -13601 | 60 | ATT | TAGAGGGGG ATT T | G | T | TAG | **ND5-comp** | IVVWKGDAGEMLLVMRNPANRLPAARRLMGFS  RVGLFSLMLVRVGKRGWPVRVRRIIRVL |
| -13577 | -13505 | 24 | ATG | GAGGTAGCG ATG G | G | G | TAG | **ND5-comp** | MRVMDRAQAFVYDMFAVSMMWSLE |

| -13484 | -12587 | 299 | ATT | AGGAAAGGT ATT C | G | C | TAG | **ND5-comp** | IPANARLPMVREVEVRGMVLSSPPIFRMSCSLLR LWMMDPEHMNSMALKKAWVQMCRNARCG WLMPIVTIMSPSWLEVEKATIFLMSFCVRAQTA ANRVVMAPKHSVRVWISGLFSARGWKRMSKKI PATTMVLEWSRAETGVGPSMAEGSQGWRPN WADLPAAARRRPSSGVRLGLAFRRAICCGSHELE CRMNHAKARMKPMSPMRLYRIAWMAAVLASA RAYHQLMSKKDMIPTPSQPMNSWNRLLAVTKIS MVIRKMSRYLKNWLMFGSEFMYHSENSMMDH  VTNNATGMNIMEK |
| --- | --- | --- | --- | --- | --- | --- | --- | --- | --- |
| -12524 | -12440 | 28 | ATA | CAGTTCGAG ATA T | G | T | TAA | **ND5-comp** | MMTSWSRHMNIVVGKRLMMKVDATMDFT |
| -12392 | -12299 | 31 | ATG | GTGGTAAGG ATG G | A | G | TAA | **ND5-comp** | MGGIREVRVRVVMVVCMVITFIWSCTKIFGA |
| -12101 | -12029 | 24 | ATA | GGTTGAGGG ATA G | G | G | TAA | **ND4-comp** | MGGEWGMGVWTWGCFLVWMRVLCC |
| -11558 | -11411 | 49 | ATA | TTATGCCTC ATA G | C | G | TAG | **ND4-comp** | MGMVQGRGRLCVLSGGWEWVWGVLYHSRLV  LRVLRQVLLTQRWGLRHGL |
| -11345 | -11153 | 64 | ATA | AAGCTAGTC ATA T | G | T | TAG | **ND4-comp** | MLSCWLRSLMVLGQWEWVVECLVSLGCCECKL VRWVGEGSLLGCRMGSMCLRSGVLAGCLIGW  W |
| -11027 | -10967 | 20 | ATA | TTTTTCGTG ATA T | G | T | TAG | **ND4-comp** | MVVHWMSGVGLPWLWGVGVR |
| -9935 | -9875 | 20 | ATG | ATCTACAAA ATG C | A | C | TAG | **COX3-comp** | MPVSGGGFEAKVMFGCKVKY |
| -9152 | -8420 | 244 | ATT | TAGGCTTGG ATT A | T | A | TAA | **ATP6-comp** | IKATAISRMVSRIRIVKMMSVEGRLMVDIARVAL PIRCMSRWPAVMLAVRRTARAIGWMSRLMVS MMTSMGMRGVGVPCGKKWARAFLILERKPMI TVPAHKGMAMARFMDSWVVGVNEWGRSPRR LVVAMKMIKDTSMRDQVRPLVLCMVIICFEVSLI SHCWVVISRLLMRYLEVGINRGGNRMISTAAGR PRIVGAMNEANRFSFILVLRVCYNFLFLWALVRE  VGGSLCLMFLVGWWGMV |
| -8258 | -8144 | 38 | ATA | ATAGGGTAA ATA G | T | G | TAG | **COX2-comp** | MRALFQRFLGELILGRWAWNCGLLHRFQSIDRS  MPPVV |
| -7964 | -7871 | 31 | ATA | GTTCTAGGA ATA T | G | T | TAA | **COX2-comp** | MMGEVCRSWRLVRRSRCTRRFSTIGGQLIWW |
| -7505 | -7388 | 39 | ATG | GAAAAAGTC ATG A | G | A | TAG |  | MEAMGLAWNQLWGVRFLPFLSRFYVYGFFECV  VGWGASM |

| -6896 | -6752 | 48 | ATT | TCATTTCAT ATT C | C | C | TAG | **COX1-comp** | IASVECGESAKYFDAGGDSDDYGSGGEMCSCVY  VYSYCKYMVCSHDKP |
| --- | --- | --- | --- | --- | --- | --- | --- | --- | --- |
| -5801 | -5729 | 24 | ATT | TCATATTGA ATT C | T | C | TAG |  | IANSKKQLQTCRGFSRLFSRRREK |
| -4742 | -4682 | 20 | ATT | TGAGTATTG ATT G | T | G | TAG | **ND2-comp** | IGSIGYGSLSGEYIVEEDSY |
| -4247 | -4151 | 32 | ATG | TGAGGGGGA ATG T | G | T | TAG | **ND1-comp** | MLEIVMGMETYHMSNARVSGRKFFHRRCMSW  S |
| -4028 | -3686 | 114 | ATT | CCTAGGAAG ATT T | A | T | TAG | **ND1-comp** | IVVVRVFIMMMFVYSAMKNRAKGPAAYSMLKP ETSSDSPSARSKGVRLVSASVEMNHIMAKGHDG RSNQRCSCVVMRVERLKEPLISNVDSRMMARVT  SYEIVWATARSAPIRA |
| -3620 | -3518 | 34 | ATA | GTGGCTAGA ATA A | A | A | TAG | **ND1-comp** | MNRRPRLRLTRGLGMGRGVHSRRAMVRAKVG  AVM |
| -3509 | -3434 | 25 | ATG | TAGAGGGTG ATG T | G | T | TAG | **ND1-comp** | MVDVAGFRGSLVKSFMASAKGCSSP |
| -3386 | -3266 | 40 | ATT | TAGCCTAGA ATT T | A | T | TAA | **ND1-comp** | IFRSVSIRNAIAIRMGTMRSRRLAMGMLLRRGIE  PLTVKF |
| -2750 | -2615 | 45 | ATT | ACTGTTTGC ATT A | T | A | TAA | **RNR2-comp** | INKLKLHRVFSSCCVMPASSRAGQFHWLKVRDS  WTLVEPFMQVPI |
| -2531 | -2303 | 76 | ATA | GTGCCTCTA ATA T | C | T | TAA | **RNR2-comp** | MLVMLEVMFLVNRRGKICRVPFTFFNLSLWACL CWVDSEGNNDLLVDCRYWAVNCQFSVLIWRRL  MRRRMFSCYLY |
| -2279 | -2195 | 28 | ATA | TATAGGGTG ATA A | G | A | TAA | **RNR2-comp** | MDWSNWVWGVQLYVWDFLGSGCWAWTLS |
| -2168 | -2090 | 26 | ATG | AGGCCTACT ATG G | A | G | TAA | **RNR2-comp** | MGVKFFTLSTRFFPSVQRAVPLWTNS |
| -1520 | -1427 | 31 | ATG | TTTAGTTAA ATG C | T | C | TAA | **RNR1-comp** | MSFEVYLRRVTGGVYALQGPVQLSTLLLVYC |
| -1400 | -1292 | 36 | ATA | TTAAGTTTC ATA G | T | G | TAG | **RNR1-comp** | MRAIVVFWGRKCSPFLATSWATPWPNVFTWVL  ALTL |
| -1091 | -1022 | 23 | ATA | GGGCTAAGC ATA T | A | T | TAG | **RNR1-comp** | MVGYLIPVWVLAIVCSDMLKPLS |
| -1016 | -950 | 22 | ATT | TCGTAGTCT ATT T | T | T | TAA | **RNR1-comp** | ILCQLEFFTTQVSFSFIGEGVI |
| -773 | -590 | 61 | ATT | TTGAGCTGC ATT C | T | C | TAA | **RNR1-comp** | IAACLMLVPFDRGDLEGELTGTGMLACVILLRAN  RKARTKPICLWGDVSPSKHFQCIALRR |
| -539 | -443 | 32 | ATG | GTTCGGGGT ATG G | G | G | TAG |  | MGLAAVCVCWVGWAGVVLMRLVVWEWEGK  MMC |
| -416 | -335 | 27 | ATA | TAAAAGTGC ATA C | T | C | TAA |  | MPPKDKIWNLVRLVLGFFVFGVWQRCV |
| -179 | -116 | 21 | ATA | TCGCCTGTA ATA T | G | T | TAG |  | MLNVGAMNNRMRQESKTDTAT |

**Table S2.2 14 chosen candidates from the kozak approach**

| **PROTEIN SEQUENCE** | **FEATURE** | **tBLASTn**  **excluding Homo/Pan/Gori lla** | **tBLASTn**  **against Mus musculus** | **Nuclear hits with**  **>70%**  **similarity** | **Antibody comments** |
| --- | --- | --- | --- | --- | --- |
| MRIEPIPENPKFSVPPITPHPKVRSAK | Non-coding region + strand | yes | no | 25 |  |
| MWRTLPWTLHNMFCHQDPTSNLPVLMNS NSMPPIPLRPTHTPPMKKLPTTHPSITYMMC LHTHYNLQHSPSNLRNMSDKRVTLME | Within nd1 + strand | yes | no | 3 |  |
| MGSPPPAGSKKVVLRLRSVSSMVMPAA | Cox1 complementary strand | no | no | 2 |  |
| MSVGRNGVGFVCSNCHFIFTLLDMGSSVIEV E | Nd2 complementary strand | yes | no | 6 | Too hydrophobic for immunisation |
| MGEISEWSLLWWQMQLLLMGHSGSGLQR STCRVVRCLVMSLLMQCQSGHLRWKERWIL GLRALQQIISYCFRGVWRVS | Cox1 complementary strand | yes | no | 3 |  |
| MGVGMEWGLLLRRGRRRWCWGCGLLVV | Cox1 complementary strand | no | no | 2 |  |
| MGEVGLLWLGRIRRRGAFGIGLWQGVLYW | Cox1 complementary strand | no | no | 3 |  |
| MGPMAYLADLTLGWGVMGGTENFGFSG MGSILMVLEMRGFKLLLFTLSK | trnM complementary strand | yes | yes | 14 | Too hydrophobic for immunisation |
| MGGGCIDEISSMGVGGENNVLVGGWLLKV HTAKR | Non-coding - strand | no | no | 0 |  |

| MCLSAVARSGGGGVWWKFFVMMSVWKV AVQTFNCYYYVLQALIN | Non-coding - strand | no | no | 1 | Too hydrophobic for  immunisation |
| --- | --- | --- | --- | --- | --- |
| **MGCSGSSVSQCYRVHTPQTKMPNAWRAP**  **VSGR** | **Non-coding - strand** | **yes** | **no** | **1** |  |
| MGGEWGMGVWTWGCFLVWMRVLCC | Nd4 complementary  strand | no | no | 1 |  |
| MGVKFFTLSTRFFPSVQRAVPLWTNS | 16Srna complementary strand SHLP2 (Cobb  et al. 2016) | yes | no | SHLP2 |  |
| MGLAAVCVCWVGWAGVVLMRLVVWEWE  GKMMC | Non-coding - strand | no | no | 0 |  |

**ANTIBODY PRODUCTION**

| START | LONG. | CODON I CO | NTEXT | Kozak -3 | Kozak +4 | CODON F | **FEATURE** |
| --- | --- | --- | --- | --- | --- | --- | --- |
| -117 | 31 | ATA | TACTGCG AC ATA G | G | G | TAA | MGCSGSSVS QCYRVHTPQT KMPNAWRAP VSGR |

**Table S3. Identification of mitochondrial smORFs and altORFs - OpenProt approach and sequences selected for antibody production**

Genomic regions without an associated transcript will not be in OpenProt (i.e. regions with no gene name, or -comp)

| **Genes** | **Present_in_OP** |
| --- | --- |
| ATP6 | 1 |
| ATP8 | 1 |
| COX1 | 1 |
| COX2 | 1 |
| COX3 | 1 |
| CYTB | 1 |
| ND1 | 1 |
| ND2 | 1 |
| ND3 | 1 |
| ND4 | 1 |
| ND4L | 1 |
| ND5 | 1 |
| RNR1 | 1 |
| RNR2 | 1 |
| TRND | 1 |
| TRNH | 1 |
| TRNI | 1 |
| TRNK | 1 |
| TRNL1 | 1 |
| TRNL2 | 1 |
| TRNQ | 1 |
| TRNS1 | 1 |
| TRNT | 1 |
| TRNW | 1 |
| TRNY | 1 |

**OpenProt predictions for genes above - AltProt (novel protein from non canonical ORF) are in green - sorted by MS detection - more info on each protein on OpenProt website Identification of mitochondrial smORFs and altORFs - OpenProt approach**

\# OpenProt release 1.3 - August 30 2018

| **Protein accession** | **Protein Type** | **Species** | **Protein length (a.a.)** | **Molecular weight (kDa)** | **Isoelectric point** | **Gene symbol** | **Transcript accession** | **Type** | **MS**  **score** | **TE**  **score** | **Domains** | **Orthology Across 10 Species = Species name : id**  **%** |
| --- | --- | --- | --- | --- | --- | --- | --- | --- | --- | --- | --- | --- |
|  |  |  |  |  |  |  |  |  |  |  |  | SC:44.09,DR:67.73, |
|  |  |  |  |  |  |  |  |  |  |  |  | MM:73.64,RN: |
| P00403 | RefProt | Homo sapiens | 227 | 25.56 | 4.44 | COX2 | COX2 | CDS | 335 | 0 | 25 | 74.09,DM:58.45,CE: |
|  |  |  |  |  |  |  |  |  |  |  |  | 44.5,BT:74.55,PT: |
|  |  |  |  |  |  |  |  |  |  |  |  | 97.8,OA:73.64 |
|  |  |  |  |  |  |  |  |  |  |  |  | DR:55.18,MM: |
|  |  |  |  |  |  |  |  |  |  |  |  | 66.13,RN:64.68, |
| P03915 | RefProt | Homo sapiens | 603 | 67.03 | 9.32 | ND5 | ND5 | CDS | 110 | 0 | 62 | DM:37.2,CE:35.31, |
|  |  |  |  |  |  |  |  |  |  |  |  | BT:71.23,PT:93.27, |
|  |  |  |  |  |  |  |  |  |  |  |  | OA:71.6 |
|  |  |  |  |  |  |  |  |  |  |  |  | MM:46.27,RN: |
| P03928 | RefProt | Homo sapiens | 68 | 7.99 | 10.56 | ATP8 | ATP8 | CDS | 88 | 0 | 7 | 47.76,BT:55.17,PT: |
|  |  |  |  |  |  |  |  |  |  |  |  | 94.12,OA:49.23 |
|  |  |  |  |  |  |  |  |  |  |  |  | DR:60.22,MM: |
|  |  |  |  |  |  |  |  |  |  |  |  | 67.03,RN:67.9,DM: |
| YP_003024035.1 | RefProt | Homo sapiens | 459 | 51.58 | 9.67 | ND4 | ND4 | CDS | 72 | 0 | 45 | 42.96,CE:32.93,BT: |
|  |  |  |  |  |  |  |  |  |  |  |  | 74.56,PT:94.99,OA: |
|  |  |  |  |  |  |  |  |  |  |  |  | 75.66 |
|  |  |  |  |  |  |  |  |  |  |  |  | DR:67.2,MM:78.33, |
|  |  |  |  |  |  |  |  |  |  |  |  | RN:77.89,DM: |
| YP_003024026.1 | RefProt | Homo sapiens | 318 | 35.66 | 6.53 | ND1 | ND1 | CDS | 71 | 0 | 32 | 49.34,CE:35.33,BT: |
|  |  |  |  |  |  |  |  |  |  |  |  | 78.55,PT:94.65,OA: |
|  |  |  |  |  |  |  |  |  |  |  |  | 77.99 |
|  |  |  |  |  |  |  |  |  |  |  |  | SC:66.05,DR:85.57, |
|  |  |  |  |  |  |  |  |  |  |  |  | MM:91.02,RN: |
| P00395 | RefProt | Homo sapiens | 513 | 57.04 | 6.7 | COX1 | COX1 | CDS | 66 | 0 | 56 | 90.62,DM:76.24,CE: |
|  |  |  |  |  |  |  |  |  |  |  |  | 60.67,BT:90.82,PT: |
|  |  |  |  |  |  |  |  |  |  |  |  | 98.83,OA:91.99 |
|  |  |  |  |  |  |  |  |  |  |  |  | SC:35.58,DR:52.86, |
|  |  |  |  |  |  |  |  |  |  |  |  | MM:75.66,RN: |
| YP_003024031.1 | RefProt | Homo sapiens | 226 | 24.82 | 10.68 | ATP6 | ATP6 | CDS | 57 | 0 | 30 | 75.66,DM:37.95,CE: |
|  |  |  |  |  |  |  |  |  |  |  |  | 32.69,BT:77.88,PT: |
|  |  |  |  |  |  |  |  |  |  |  |  | 94.25,OA:77.43 |
|  |  |  |  |  |  |  |  |  |  |  |  | SC:43.85,DR:80.84, |
|  |  |  |  |  |  |  |  |  |  |  |  | MM:86.97,RN: |
| YP_003024032.1 | RefProt | Homo sapiens | 261 | 29.95 | 7.34 | COX3 | COX3 | CDS | 49 | 0 | 28 | 87.74,DM:65.25,CE: |
|  |  |  |  |  |  |  |  |  |  |  |  | 43.31,BT:87.69,PT: |
|  |  |  |  |  |  |  |  |  |  |  |  | 97.32,OA:86.97 |

| YP_003024027.1 | RefProt | Homo sapiens | 347 | 38.96 | 10.3 | ND2 | ND2 | CDS | 37 | 0 | 39 | DR:44.8,MM:58.06, RN:57.76,DM: 41.64,BT:63.93,PT:  96.81,OA:63.85 |
| --- | --- | --- | --- | --- | --- | --- | --- | --- | --- | --- | --- | --- |
| YP_003024038.1 | RefProt | Homo sapiens | 380 | 42.72 | 8.22 | CYTB | CYTB | CDS | 28 | 0 | 37 | SC:50.27,DR:70.26, MM:78.57,RN: 78.63,DM:62.81,CE:  44.87,BT:78.89,PT:  93.67,OA:77.84 |
| YP_003024033.1 | RefProt | Homo sapiens | 115 | 13.19 | 4.08 | ND3 | ND3 | CDS | 17 | 0 | 13 | DR:57.52,MM:  70.79,RN:73.03,  DM:47.62,BT:73.91, PT:94.78,OA:73.04 |
| IP_306405 | AltProt | Homo sapiens | 61 | 6.38 | 11.88 | RNR1 | RNR1 | rRNA | 1 | 0 | 8 | PT:90 |
| IP_306387 | AltProt | Homo sapiens | 32 | 3.66 | 11.93 | ND2 | ND2 | CDS | 0 | 0 | 0 | PT:65.62 |
| IP_306389 | AltProt | Homo sapiens | 33 | 3.7 | 9.51 | COX1 | COX1 | CDS | 0 | 0 | 0 |  |
| IP_306392 | AltProt | Homo sapiens | 39 | 4.56 | 10.98 | ATP8 | ATP8 | CDS | 0 | 0 | 0 | PT:79.49 |
| IP_306398 | AltProt | Homo sapiens | 35 | 3.82 | 6.5 | ND4 | ND4 | CDS | 0 | 0 | 0 | PT:85.71 |
| IP_306399 | AltProt | Homo sapiens | 59 | 6.91 | 12.81 | ND4 | ND4 | CDS | 0 | 0 | 0 | PT:82.76 |
| IP_306403 | AltProt | Homo sapiens | 49 | 5.22 | 4.14 | CYTB | CYTB | CDS | 0 | 0 | 0 | PT:74.29 |
| IP_306404 | AltProt | Homo sapiens | 31 | 3.43 | 10.22 | RNR1 | RNR1 | rRNA | 0 | 0 | 0 |  |
| IP_306406 | AltProt | Homo sapiens | 37 | 4.35 | 9.95 | RNR1 | RNR1 | rRNA | 0 | 0 | 0 | PT:86.49 |
| IP_306407 | AltProt | Homo sapiens | 32 | 3.27 | 6.5 | RNR1 | RNR1 | rRNA | 0 | 0 | 0 | PT:93.75 |
| IP_306408 | AltProt | Homo sapiens | 33 | 3.81 | 8.56 | RNR1 | RNR1 | rRNA | 0 | 0 | 0 |  |
| IP_306409 | AltProt | Homo sapiens | 32 | 3.71 | 9.44 | RNR2 | RNR2 | rRNA | 0 | 0 | 0 |  |
| IP_306410 | AltProt | Homo sapiens | 33 | 3.83 | 10.9 | RNR2 | RNR2 | rRNA | 0 | 0 | 0 |  |
| IP_306411 | AltProt | Homo sapiens | 29 | 3.33 | 4.31 | RNR2 | RNR2 | rRNA | 0 | 0 | 0 | BT:93.1,PT:100,OA:  93.1 |
| YP_003024034.1 | RefProt | Homo sapiens | 98 | 10.74 | 6.2 | ND4L | ND4L | CDS | 0 | 0 | 12 | DR:55.68,MM:  66.33,RN:69.15,  DM:36.67,BT:73.47, PT:98.98,OA:76.53 |

**ANTIBODY PRODUCTION (antigen in red)**

| IP_306387 | AltProt | Homo sapiens | 32 | 3.66 | 11.93 | ND2 | ND2 | CDS | 0 | 0 | 0 | PT:65.62 | **MTKTSPHLNHMPNLSLTKRKPSPHSL**  **NLIHHS** |
| --- | --- | --- | --- | --- | --- | --- | --- | --- | --- | --- | --- | --- | --- |
| IP_306389 | AltProt | Homo sapiens | 33 | 3.7 | 9.51 | COX1 | COX1 | CDS | 0 | 0 | 0 |  | **MPNAPLRLIRPNHSSPTSPISPSPSCWHHYTTN** |
| IP_306398 | AltProt | Homo sapiens | 35 | 3.82 | 6.5 | ND4 | ND4 | CDS | 0 | 0 | 0 | PT:85.71 | **MSSSKPHLSPPWLSSPDEATSQNAWTQAHTSYSTP** |
| IP_306403 | AltProt | Homo sapiens | 49 | 5.22 | 4.14 | CYTB | CYTB | CDS | 0 | 0 | 0 | PT:74.29 | **MAESSATFTPMAPQYSLSASSYTSGEAYITDHFSTQK PETSALSSCLQL** |

### Table S4. Identification of mitochondrial smORFs and altORFs - MS approach and sequences selected for antibody production

**Legend**

### Tables: PepQuery outputs

**Prot_Accession**	Accession number for the protein (unique ID)

**Description**	Accession number for the protein (output by pepquery)

**Prot_Seq**	Protein sequence

**PepQuery_hits**	Number of peptide spectrum matches (PSMs) reported by PepQuery = number of PSMs with higher score than with any reference peptides **without** PTMs **Confident_hits**	Number of confident PSMs reported by PepQuery = number of PSMs with higher score than with any reference peptides **with or without any PTMs Best_Score**	Score of the best confident PSM

**pvalue**	*p-*value of the best confident PSM

**Peptide_Sequence**	Peptide sequence yielding the best PSM

**Peptide_Check_Mass**	Theoretical mass of the peptide *(may vary from experimental mass based on precursor charge, PTMs, and error)*

**Exp_Pep_Mass**	Observed mass of the peptide (experimental)

**Error_mass_ppm**	Error of the peptide mass in ppm

**Validation	#N/A** if no confident PSMs; **No** if no confident PSMs with an error less than |4,5| ppm; **Yes** if confident PSM with an error less than |4,5| ppm

**Best PSM for each protein**

| **Prot_Acces**  **sion** | **Description** | **Prot_Seq** | **PepQuery_**  **hits** | **Confident_**  **hits** | **Best_Score** | **pvalue** | **Peptide_Sequenc**  **e** | **Peptide_Check_M**  **ass** | **Exp_Pep_Mass** | **Error_mass_p**  **pm** | **Validation** |
| --- | --- | --- | --- | --- | --- | --- | --- | --- | --- | --- | --- |
| SB_0001 | SB_0001 | IWYFRLGGMHA MALRDAGAGAPY VAVSVFDSCLILLFI  APTFNITGEHTY | 37 | 4 | 18.28136369 | 0.001020408163 | LGGMHAMALR | 1055.536813 | 1215.632296 | 1.251744129 | Yes |
| SB_0002 | SB_0002 | MMMTIECLHSHF PHRHHNKKFPPN PPSPASGHST | 31 | 0 | 0 | #N/A | #N/A | #N/A | #N/A | #N/A | #N/A |
| SB_0003 | SB_0003 | IIFPSHSHTTNLINT TPAHPTQHTHTA ANPMPRTNQTPK TPPTVYVAYLLKA MHWKCLDGLTSP HKQMGLVLAFLLA LSKITHASIPVPVSS PSKSPRSKGTSIKH AAMQLKTLSLATP  PRETAVINL | 52 | 3 | 21.5925048 | 0.008743169399 | TLSLATPPR | 954.5498034 | 1098.655792 | -3.548999013 | Yes |
| SB_0004 | SB_0004 | MKLKLTWVVKNS SWHKMDYESGFN MSEHTMAKTQTG IRYPTMLSPKPQQ LNQQNCSPEHYEP QLKTQRTWRCFM SL | 23 | 1 | 12.50724417 | 0.000999000999 | MDYESGFNMSEH TMAKTQTGIR | 2533.103525 | 2869.269804 | 7.891735204 | No |
| SB_0005 | SB_0005 | MPPSSANPDEGY KVSASTHVKTLGQ GVAHEVARNGLH FLPQKTTMALMK LKGRRWI | 111 | 8 | 31.85069597 | 0.001998001998 | TLGQGVAHEVAR | 1236.657452 | 1380.764859 | -3.848103668 | Yes |
| SB_0006 | SB_0006 | IYMEETSRNMVSV LESALGRTRV | 55 | 4 | 22.97199697 | 0.000999000999 | IYMEETSRNMVSV LESALGR | 2284.119098 | 2572.347638 | -9.47025908 | No |
| SB_0007 | SB_0007 | MVGRFMGRGDK PTEPGDSWLSKM ES | 54 | 12 | 27.52984605 | 0.001998001998 | MVGRFMGRGDKP TEPGDSWLSK | 2450.183426 | 2914.478541 | 0.3292530069 | Yes |
| SB_0008 | SB_0008 | ILLRMSLRQIKTLN WQLTAQYLQSTN KSLLPSLSTQHRHA HKERLKKVKGTRQ ILPRLFTKNITSSITS  IRGTACPVTHV | 43 | 1 | 23.99545008 | 0.000999000999 | LFTKNITSSITSIR | 1579.893313 | 1868.103082 | -2.998073063 | Yes |

| SB_0009 | SB_0009 | MGTCMNGSTRV QLSLTFNQWNWP AREEAGMTQQDE KTLWSFNLLMQT VPNKPTGPKLPNL H | 20 | 0 | 0 | #N/A | #N/A | #N/A | #N/A | #N/A | #N/A |
| --- | --- | --- | --- | --- | --- | --- | --- | --- | --- | --- | --- |
| SB_0010 | SB_0010 | IDPMTWPTEQVT LGMTAQSYSRVHI NNRVYDLDVGSG HPDGAAAIKGSFV QRLKSYVIWVQTG VIQVGFYLXSNSSL YERTREMRPTSQS AFPRKWYHLNLVL YPHPPKNRVC | 53 | 1 | 20.4256365 | 0.000999000999 | EMRPTSQSAFPR | 1405.6772 | 1565.764758 | 6.03635534 | No |
| SB_0011 | SB_0011 | IPLLNNMPMANLL LLIVPILIAMAFLM LTERKILGYMQLR KGPNVVGPYGLL QPFADAMKLFTKE PLKPATSTITLYITA PTLALTIALLLWTP LPMPNPLVNLNL GLLFILATSSLAVYS ILWSGWASNSNY ALIGALRAVAQTIS YEVTLAIILLSTLLM SGSFNLSTLITTQE HLWLLLPSWPLA MMWFISTLAETN RTPFDLAEGESELV SGFNIEYAAGPFAL FFMAEYTNIIMM NTLTTTIFLGTTYD ALSPELYTTYFVTK TLLLTSLFLWIRTA YPRFRYDQLMHLL WKNFLPLTLALLM WYVSMPITISSIPP QT | 97 | 75 | 90.43099497 | 0.000999000999 | KGPNVVGPYGLLQ PFADAMK | 2101.102977 | 2533.421241 | -4.745353471 | No |
| SB_0012 | SB_0012 | MQNPPHSSPHSS PLPRYSYLSPLLY | 8 | 0 | 0 | #N/A | #N/A | #N/A | #N/A | #N/A | #N/A |
| SB_0013 | SB_0013 | IYSPMLHSAILPHP HWCSPTVDYSLQ TTKTLEHYTYYSAH ELES | 1 | 0 | 0 | #N/A | #N/A | #N/A | #N/A | #N/A | #N/A |

| SB_0014 | SB_0014 | MTLSKLNYRLNP MYLNGTCSASRST RRYFPYHRRAYHL SWSRPHNHFPYLL PSPVCPFPNTHNK TN | 38 | 0 | 0 | #N/A | #N/A | #N/A | #N/A | #N/A | #N/A |
| --- | --- | --- | --- | --- | --- | --- | --- | --- | --- | --- | --- |
| SB_0015 | SB_0015 | MVLNLRVHRLRRT NLQLLHTSPIIPRT RRPATPWRWQSS STPDWSPHSYNN YITRRLALMSCPHI RLKNRCNSRTSKP NHFHRYTTGGML RSMLWNLWSKP QFHAHRPRINSPK NLWNRARIYPMA PPLPPLEPTVKLT | 49 | 2 | 19.26859725 | 0.008849557522 | NLWNRAR | 928.4991084 | 1072.603732 | -2.365401482 | Yes |
| SB_0016 | SB_0016 | IKRTNTSLQWNAP TKYYRMAHHNYP HTPYTIPHHPTKNI KHKLPPTSLTKAH KNKKL | 29 | 0 | 0 | #N/A | #N/A | #N/A | #N/A | #N/A | #N/A |
| SB_0017 | SB_0017 | MNENLFASFIAPTI LGLPAAVLIILFPPL LIPTSKYLINNRLIT TQQWLIKLTSKQ MMTMHNTKGRT WSLMLVSLIIFIAT TNLLGLLPHSFTPT TQLSMNLAMAIPL WAGTVIMGFRSKI KNALAHFLPQGTP TPLIPMLVIIETISLL IQPMALAVRLTAN ITAGHLLMHLIGS ATLAMSTINLPSTL IIFTILILLTILEIAVA LIQAYVFTLLVSLYL HDNT | 130 | 89 | 51.46183017 | 0.001020408163 | LITTQQWLIK | 1242.733572 | 1530.93992 | -1.428598097 | Yes |
| SB_0018 | SB_0018 | ILPSYKPQSTSSLPS PFPTASTAQHFL | 0 | 0 | 0 | #N/A | #N/A | #N/A | #N/A | #N/A | #N/A |
| SB_0019 | SB_0019 | MWFDYFCMSPSI DEGLTLLV | 2 | 0 | 0 | #N/A | #N/A | #N/A | #N/A | #N/A | #N/A |
| SB_0020 | SB_0020 | ITTTDMTFQKTHN LNQHNHPQPNY | 9 | 1 | 21.06164271 | 0.00515995872 | ITTTDMTFQK | 1184.574696 | 1488.764668 | 6.110094777 | No |

| SB_0021 | SB_0021 | IHSHRTNHILYLLR NHTYPHLGYHHP MRQPARTPERRH MLPILHPSRLPSPT HRTNLHSQHPRLT KHSTTHSHCPRTIK LLSQQLNMTSLHN SFYSKDTSLRTPLM TP | 37 | 0 | 0 | #N/A | #N/A | #N/A | #N/A | #N/A | #N/A |
| --- | --- | --- | --- | --- | --- | --- | --- | --- | --- | --- | --- |
| SB_0022 | SB_0022 | MRHNYNKLHLPT TNRPKIAHCMLFN QPHSPRSNSHSHP NPLKLHRRSHSHN RPRAYILITILPSKLK LRTHSQSHHNPLS RTSNSTPTNSFLM TSSKPR | 62 | 11 | 19.17011656 | 0.002018163471 | LHLPTTNRPK | 1175.677457 | 1463.877578 | 2.758316996 | Yes |
| SB_0023 | SB_0023 | IHTRKHPHVHTPIP HSPPIPQPRHHYR VFLL | 42 | 1 | 14.03856455 | 0.002421307506 | HHYRVFLL | 1083.597754 | 1227.709476 | -7.84342575 | No |

| SB_0024 | SB_0024 | ILVQLQMKVMTM HTTMTTLTLTSLIP PILTTLVNPNKKNS YPHYVKSIVASTFII SLFPTTMFMCLD QEVIISNWHWAT TQTTQLSLSFKLDY FSMMFIPVALFVT WSIMEFSLWYMN SDPNINQFFKYLLI FLITMLILVTANNL FQLFIGWEGVGI MSFLLISWWYAR ADANTAAIQAILY NRIGDIGFILALAW FILHSNSWDPQQ MALLNANPSLTPL LGLLLAAAGKSAQ LGLHPWLPSAME GPTPVSALLHSST MVVAGIFLLIRFHP LAENSPLIQTLTLC LGAITTLFAAVCAL TQNDIKKIVAFSTS SQLGLMMVTIGIN QPHLAFLHICTHA FFKAMLFMCSGSII HNLNNEQDIRKM GGLLKTMPLTSTS LTIGSLALAGMPFL TGFYSKDHIIETAN MSYTNAWALSITL IATSLTSAYSTRMI LLTLTGQPRFPTLT NINENNPTLLNPIK RLAAGSLFAGFLIT NNISPASPFQTTIP LYLKLTALAVTFLG LLTALDLNYLTNKL KMKSPLCTFYFSN MLGFYPSITHRTIP YLGLLTSQNLPLLL LDLTWLEKLLPKTI SQHQISTSIITSTQ KGMIKLYFLSFFFP LILTLLLIT | 113 | 63 | 76.17222727 | 0.000999000999 | FPTLTNINENNPTL LNPIKR | 2308.253868 | 2596.46168 | -1.396327951 | Yes |
| --- | --- | --- | --- | --- | --- | --- | --- | --- | --- | --- | --- |

| SB_0025 | SB_0025 | IPPSNLNYNMYTN KQCSTSNYY | 41 | 0 | 0 | #N/A | #N/A | #N/A | #N/A | #N/A | #N/A |
| --- | --- | --- | --- | --- | --- | --- | --- | --- | --- | --- | --- |
| SB_0026 | SB_0026 | MIMQSPRTNRILP NQPWPLSFMNYS ASYTIKVYHNHHPI MLFHPQHQSYLH R | 34 | 0 | 0 | #N/A | #N/A | #N/A | #N/A | #N/A | #N/A |
| SB_0027 | SB_0027 | MTSPKIQNNNTP DHTANNQY | 6 | 0 | 0 | #N/A | #N/A | #N/A | #N/A | #N/A | #N/A |
| SB_0028 | SB_0028 | MHHYSRTDYNHD QWYEKPSLYFNYK NTNDPNTQN | 2 | 0 | 0 | #N/A | #N/A | #N/A | #N/A | #N/A | #N/A |
| SB_0029 | SB_0029 | IHRPPHPIQHLRM MKLRLTPWRLPD PPNHHRTIPSHAL LTRRLNRLFINRPH HSRRKLWLNHPLP SRQWRLNILYLPL PTHRARPMLRIISL LRNLKHRHYPPAC NYSNSLHRLCPPV RPNIILRGHSNYKL TIRHPMHWDRPS SMNLRRLLSRQSH PHTILYLSLHLALH YCSPSNTPPPILAR  NGIKQPPRNHLPF R | 121 | 2 | 13.82029738 | 0.004424778761 | LRLTPWR | 940.5606422 | 1084.660857 | 1.728571636 | Yes |
| SB_0030 | SB_0030 | INTILTRPPRRPRQ LYPSQPLKHPSPH QARMMFPIRLHN  SPIRP | 30 | 0 | 0 | #N/A | #N/A | #N/A | #N/A | #N/A | #N/A |
| SB_0031 | SB_0031 | ITIHPHPSNNPHPP YIQTTKHNISPTKPI TLLTPSRRPPHSNL NRRTTSKLPFYHH WTSSIRTMLHNN PNPNTNYLPNWK QNTQMGLSL | 49 | 1 | 16.8933154 | 0.003144654088 | RPPHSNLNR | 1089.579146 | 1233.689229 | -6.478610557 | No |
| SB_0032 | SB_0032 | MHQSCKPEMKTF FQGQIREKVFNSTI STQS | 20 | 2 | 13.83337106 | 0.003003003003 | MHQSCKPEMK | 1217.535493 | 1722.853313 | 2.770027249 | Yes |

| SB_0033 | SB_0033 | MYFVHYCQPPW MLYGTMNTWPP VVHKNPIHIKTPSP CLQASTAINPQLS HINCNSKATPHPL GYQQTYPPLTVHS T | 22 | 0 | 0 | #N/A | #N/A | #N/A | #N/A | #N/A | #N/A |
| --- | --- | --- | --- | --- | --- | --- | --- | --- | --- | --- | --- |
| SB_0034 | SB_0034 | MAHYSQIPSRPH GWPPSDRGPLTTI LREINIPHKSATLL APGP | 10 | 0 | 0 | #N/A | #N/A | #N/A | #N/A | #N/A | #N/A |
| SB_0035 | SB_0035 | MSQYLSLIPASSYY LSHLRSMLQANM LTKVC | 14 | 0 | 0 | #N/A | #N/A | #N/A | #N/A | #N/A | #N/A |
| SB_0036 | SB_0036 | MTKNFHQTPPPP LLATALKHISAKPQ  KQRTLTPA | 3 | 0 | 0 | #N/A | #N/A | #N/A | #N/A | #N/A | #N/A |
| SB_0037 | SB_0037 | MLLISSMQPPPILP STHTPLLTPYPEPT KPQRHPPQFM | 0 | 0 | 0 | #N/A | #N/A | #N/A | #N/A | #N/A | #N/A |
| SB_0038 | SB_0038 | MQASPFQWVHP LNHHDQKEQASS  TQQCSSKRLA | 12 | 0 | 0 | #N/A | #N/A | #N/A | #N/A | #N/A | #N/A |
| SB_0039 | SB_0039 | MLTPGLVNFVPAT AVTRLTQVNRSRR KECFRSPPPQ | 10 | 0 | 0 | #N/A | #N/A | #N/A | #N/A | #N/A | #N/A |
| SB_0040 | SB_0040 | MEIETWRNRYST ARERWKIMTKHN MARTNPYTFCM  MN | 33 | 0 | 0 | #N/A | #N/A | #N/A | #N/A | #N/A | #N/A |
| SB_0041 | SB_0041 | MTLQGEPKLRPPK PDELPKNS | 10 | 0 | 0 | #N/A | #N/A | #N/A | #N/A | #N/A | #N/A |
| SB_0042 | SB_0042 | IKKAFKLNTHYLKN PKHMTELLTPNW TNLSPYRRTNVSM SNMKTFSSA | 71 | 8 | 16.0339017 | 0.008743169399 | TNVSMSNMK | 1010.452479 | 1330.657952 | -8.639537899 | No |
| SB_0043 | SB_0043 | MAPRGFSCLLLLT SEIDLPVKRRA | 4 | 0 | 0 | #N/A | #N/A | #N/A | #N/A | #N/A | #N/A |
| SB_0044 | SB_0044 | IKNFGWGDLGAE PNLRAVHAKTSPV KANYYTQLIQ | 20 | 1 | 23.17994279 | 0.000999000999 | NFGWGDLGAEPN LR | 1544.737155 | 1688.850186 | -6.473704492 | No |
| SB_0045 | SB_0045 | ILESMSTMGFTTS MLDQDIPMVQPL  LKVRLFND | 1 | 0 | 0 | #N/A | #N/A | #N/A | #N/A | #N/A | #N/A |

| SB_0046 | SB_0046 | MAEPGNRMKLKT LQSEVQFLFLTTYP WPTSYSSLYPF | 18 | 3 | 13.93237678 | 0.008869179601 | MAEPGNR | 773.3490056 | 917.4533302 | -2.447302268 | Yes |
| --- | --- | --- | --- | --- | --- | --- | --- | --- | --- | --- | --- |
| SB_0047 | SB_0047 | MSPYPLQSPAFPL  KPKKYVW | 0 | 0 | 0 | #N/A | #N/A | #N/A | #N/A | #N/A | #N/A |
| SB_0048 | SB_0048 | ISRTMRIEPIPENP KFSVPPITPHPKVR SAK | 10 | 0 | 0 | #N/A | #N/A | #N/A | #N/A | #N/A | #N/A |
| SB_0049 | SB_0049 | IFYLSRPRNKHASF YSSSNQKNKPSFH RSCHQVFPHASN RIHNPSNSYPLQQ YTLRTMNHNQYY QSMLIINNHNSYS NKTRNSPLSLLSPR GYPRHPSDIRPASS HMTKTSPHLNHM PNLSLTKRKPSPHS LNLIHHSRQLRWI KPNPATQNLSMLL NYPHRMNNSSST  VQP | 122 | 4 | 12.05265959 | 0.007658643326 | IFYLSRPR | 1050.59742 | 1338.79671 | 3.632896529 | Yes |
| SB_0050 | SB_0050 | IPTTQLKLQHHDP TTISHLKQANMTN TLNSIHPPLPRRPA PANRLFAQMGHY RRIHKKQ | 41 | 1 | 20.32219288 | 0.000999000999 | QANMTNTLNSIHP PLPR | 1902.973367 | 2047.067215 | 4.036097747 | Yes |
| SB_0051 | SB_0051 | MTVWTYKTHPIP PHTHRPYHATPTY LPFYTNNLMEI | 0 | 0 | 0 | #N/A | #N/A | #N/A | #N/A | #N/A | #N/A |
| SB_0052 | SB_0052 | ISVTAKDCKTPLCI  NWTQISHFN | 2 | 0 | 0 | #N/A | #N/A | #N/A | #N/A | #N/A | #N/A |
| SB_0053 | SB_0053 | MKITSELVKRGLTP VFRFTVQCFTQPF YLTPTDVRRPLTIL YKPQRHWNTMPI IRRMSWSPRHSSK PPYSSRAGPARQP SR | 31 | 2 | 17.19087616 | 0.001001001001 | HWNTMPIIRR | 1322.702959 | 1466.810724 | -3.863706822 | Yes |

| SB_0054 | SB_0054 | ICNNLLHSNTHHN RRLWQLTSSPNN RCPRYGVSPHKQ HKLLTLTSLSPTPA RICYSGGRSRNRL NSLPSLSRELLPPW SLRRPNHLLLTPSR CLLYLRGHQFHHN NYQYKTPCHNPM PNAPLRLIRPNHSS PTSPISPSPSCWH HYTTNRPQPQHH LLRPRRRRRPHSM PTPILIFRSPWSLYS YPTRLRNNLPYCN LLLRKKRTIWMHR YGLSYDINWLPRV YRVSTPYIYSRNRR RHTSMFHLRYHN HRYPHRRQSI | 156 | 8 | 27.54786054 | 0.001154734411 | ELLPPWSLR | 1109.623299 | 1253.713095 | 9.8081714 | No |
| --- | --- | --- | --- | --- | --- | --- | --- | --- | --- | --- | --- |
| SB_0055 | SB_0055 | MICCSALSPRIHLS FHRRWPDWHCIS KLITRHRTTRHVLR CSPLPLCPINRSCIC HHRRLHSLISPILRL HPRPNLRQNPFH YHIHRRKSNFLPTT LSRPIRNAPTLLGL PRCMHHMKHPII CRLIHFSNSSNINN FHDLRSLRFEAKSP NSRRTLHKPGVT MWMPPTLPHIRR TRMHKI | 99 | 8 | 26.39510071 | 0.000999000999 | TTRHVLRCSPLPLC PINR | 2075.124397 | 2333.278658 | -3.949405601 | Yes |

| SB_0056 | SB_0056 | MAHAAQVGLQD ATSPIMEELITFHD HALMIIFLICFLVLY ALFLTLTTKLTNTN ISDAQEMETVWTI LPAIILVLIALPSLRI LYMTDEVNDPSLT IKSIGHQWYWTYE YTDYGGLIFNSYM LPPLFLEPGDLRLL DVDNRVVLPIEAPI RMMITSQDVLHS WAVPTLGLKTDAI PGRLNQTTFTATR PGVYYGQCSEICG ANHSFMPIVLELIP LKIFEMGPVFTL | 779 | 656 | 76.27685888 | 0.000999000999 | MMITSQDVLHSW AVPTLGLK | 2226.154024 | 2514.353794 | 1.755039781 | Yes |
| --- | --- | --- | --- | --- | --- | --- | --- | --- | --- | --- | --- |
| SB_0057 | SB_0057 | MPQLNTTVWPT MITPMLLTLFLITQ LKMLNTNYHLPPS PKPMKMKNYNKP WEPKWTKICSLHS LPPQS | 328 | 277 | 64.10598268 | 0.000999000999 | MLNTNYHLPPSPK PMK | 1866.948398 | 2299.260345 | -2.484358335 | Yes |
| SB_0058 | SB_0058 | MTPNRGPLSPPN DLRPSHVISLPLHN APHTRPTNQHTN HMPMMARCNTR KHMPRPPHTTCP KRPSMRDNPIYYL RSFFLRRIFLSLLPL QPSPYPPIRRALAP NRHHPAKSPRSPT PKHIRITRIRSINHL SSP | 35 | 2 | 24.59374409 | 0.000999000999 | RPSMRDNPIYYLR | 1679.856552 | 1968.066962 | -3.172199407 | Yes |
| SB_0059 | SB_0059 | IFCSHRLPRTSRHY WLNFPHYLLHPPT NISLYIQTSLWLRS RRLMLAFCRCGLT ISVCLHLLMRVLLF | 10 | 0 | 0 | #N/A | #N/A | #N/A | #N/A | #N/A | #N/A |
| SB_0060 | SB_0060 | ILTTTTQRLHRKIH PLRVRLRPYIPRPR PFLHKILLSSYYLLII WSRNCPPFTPTM SPTNN | 18 | 2 | 21.63645375 | 0.001005025126 | ILTTTTQRLHR | 1338.773144 | 1482.86694 | 5.600029957 | No |
| SB_0061 | SB_0061 | INHHPSPKSGLWV TTKRIRLNRIGM | 7 | 0 | 0 | #N/A | #N/A | #N/A | #N/A | #N/A | #N/A |

| SB_0062 | SB_0062 | IYHLTSRNTSMSLT PHILPTMPRRNNT IAVHYSYSHNPQH PLPLSQYCAYCHTS LCRLRSSGGPSPTS LNLQHMWPRLRT | 64 | 0 | 0 | #N/A | #N/A | #N/A | #N/A | #N/A | #N/A |
| --- | --- | --- | --- | --- | --- | --- | --- | --- | --- | --- | --- |
| SB_0063 | SB_0063 | MLKLIVPTIMLLPL TWLSKKHMIWIN TTTHSLIISIIPLLFF NQINNNLFSCSPT FSSDPLTTPLLMLT TWLLPLTIMASQR HLSSEPLSRKKLYL SMLISLQISLIMTF TATELIMFYIFFETT LIPTLAIITRWGNQ PERLNAGTYFLFYT LVGSLPLLIALIYTH NTLGSLNILLLTLTA QELSNSWANNLM WLAYTMAFMVK MPLYGLHLWLPK AHVEAPIAGSMVL AAVLLKLGGYGM MRLTLILNPLTKH MAYPFLVLSLWG MIMTSSICLRQTD LKSLIAYSSISHMA LVVTAILIQTPWSF TGAVILMIAHGLT SSLLFCLANSNYER THSRIMILSQGLQ TLLPLMAFWWLL ASLANLALPPTINL LGELSVLVTTFSW SNITLLLTGLNMLV TALYSLYMFTTTQ WGSLTHHINNMK PSFTRENTLMFM HLSPILLLSLNPDIIT GFSSCKYSLTKTSD CESDNRGLRPLIYR ESSQELLTHAPMS NNMAFSTFKG | 87 | 8 | 84.24539063 | 0.000999000999 | AHVEAPIAGSMVL AAVLLK | 1889.080786 | 2177.288583 | -1.661575297 | Yes |
| SB_0064 | SB_0064 | MPTQQPFKQSYT TVSAMSVSSSP | 8 | 0 | 0 | #N/A | #N/A | #N/A | #N/A | #N/A | #N/A |

| SB_0065 | SB_0065 | ITMYTPTNNVQPV TTTNQRP | 0 | 0 | 0 | #N/A | #N/A | #N/A | #N/A | #N/A | #N/A |
| --- | --- | --- | --- | --- | --- | --- | --- | --- | --- | --- | --- |
| SB_0066 | SB_0066 | IIQLPTLLKFTTTTT PSYSFTHSTNPTSI ANPTKTLTKTSTP DPHASGYSSMAIA VVYPKTTIIPPK | 16 | 0 | 0 | #N/A | #N/A | #N/A | #N/A | #N/A | #N/A |
| SB_0067 | SB_0067 | MGEGLEENPTNPI TKPTLNRNKAYIIIL ARTTTTTNDMKN HRCISTTRTPMTP MRKTNPLMKLIN HSFIDLPTPSNISA WWNFGSLLGACL ILQITTGLFLAMHY SPDASTAFSSIAHI TRDVNYGWIIRYL HANGASMFFICLF LHIGRGLYYGSFLY SETWNIGIILLLAT MATAFMGYVLP WGQMSFWGATV ITNLLSAIPYIGTDL VQWIWGGYSVDS PTLTRFFTFHFILPF IIAALATLHLLFLHE TGSNNPLGITSHS DKITFHPYYTIKDA LGLLLFLLSLMTLTL FSPDLLGDPDNYT LANPLNTPPHIKPE WYFLFAYTILRSVP NKLGGVLALLLSILI LAMIPILHMSKQQ SMMFRPLSQSLY WLLAADLLILTWI GGQPVSYPFTIIG QVASVLYFTTILIL MPTISLIENKMLK WACPCSMN | 63 | 7 | 21.38188719 | 0.002212389381 | KTNPLMK | 830.4683856 | 1262.772811 | 1.41407854 | Yes |
| SB_0068 | SB_0068 | ILCSFMGKQIWVP PKYWLTHQQPLCI SYITASHHEYCTVP | 13 | 4 | 16.55336317 | 0.001092896175 | ILCSFMGK | 897.4452082 | 1242.673333 | -2.022001366 | Yes |
| SB_0069 | SB_0069 | MKTQSTSKPPPH AYKQVQQSTLNY  HTSTATPKPPLTH | 5 | 0 | 0 | #N/A | #N/A | #N/A | #N/A | #N/A | #N/A |

| SB_0070 | SB_0070 | ITVKSLLVPMDDP PQMGVPWPPSSV KSMSRTRVLLSSLR AHNTWG | 3 | 0 | 0 | #N/A | #N/A | #N/A | #N/A | #N/A | #N/A |
| --- | --- | --- | --- | --- | --- | --- | --- | --- | --- | --- | --- |
| SB_0071 | SB_0071 | INHSRELSMHLVF SSGGYARDSIARR WSRSTLCRSICLW FLPHPIIYRTYVQY YRRTYLLKCVN | 60 | 0 | 0 | #N/A | #N/A | #N/A | #N/A | #N/A | #N/A |
| SB_0072 | SB_0072 | MLVGHNNNNW MSAQPLSTQTS | 0 | 0 | 0 | #N/A | #N/A | #N/A | #N/A | #N/A | #N/A |
| SB_0073 | SB_0073 | ISTKPPLPRFWPQ HLNTSLPNPKNKE P | 4 | 0 | 0 | #N/A | #N/A | #N/A | #N/A | #N/A | #N/A |
| SB_0074 | SB_0074 | ISNFIFWRYALLTV  TPQLTHYFPLPLPY Y | 2 | 0 | 0 | #N/A | #N/A | #N/A | #N/A | #N/A | #N/A |
| SB_0075 | SB_0075 | MEAGVKSVLDHP LPNKAKTHLSCKKL  QLTQNRLRKWL | 5 | 0 | 0 | #N/A | #N/A | #N/A | #N/A | #N/A | #N/A |
| SB_0076 | SB_0076 | MNPDQPHHLLLS LYTAIFSKPWWRL QSKRKYPRKDVRS RCSPWGGKKWAT FSTPENYDSPYET | 18 | 0 | 0 | #N/A | #N/A | #N/A | #N/A | #N/A | #N/A |
| SB_0077 | SB_0077 | ICPQNPLNPLVNL TVSPKRNSSLDTR KKPCRESKKFNTH SRPKSSHQLRKRSS STPTT | 17 | 0 | 0 | #N/A | #N/A | #N/A | #N/A | #N/A | #N/A |
| SB_0078 | SB_0078 | INSPMSTINQQVII TLTVNPTQACS | 0 | 0 | 0 | #N/A | #N/A | #N/A | #N/A | #N/A | #N/A |
| SB_0079 | SB_0079 | MITCSLNRDLYEW LHEGSAVSYF | 10 | 1 | 12.12041551 | 0.009836065574 | MITCSLNR | 936.4520848 | 1153.56304 | 6.508987843 | No |
| SB_0080 | SB_0080 | ISVGATSEQNPTSE QYMLRLHQSKRTT MLNWSNNLTNG TSYPRDNSAILF | 47 | 0 | 0 | #N/A | #N/A | #N/A | #N/A | #N/A | #N/A |
| SB_0081 | SB_0081 | IKVLRDLSSDRSNP GRFLSXFKFLPVRK DKRNKAYFTKRLP P | 33 | 1 | 22.79775127 | 0.003058103976 | RNKAYFTKR | #N/A | 1614.978081 | -6.018171773 | No |
| SB_0082 | SB_0082 | MMSSQLSIMPTP TQEQGLLRWQSP VIA | 19 | 0 | 0 | #N/A | #N/A | #N/A | #N/A | #N/A | #N/A |

| SB_0083 | SB_0083 | IPNAYRTKNSRLYT TTQRPQRCRPLRA TTTLRWRHKTLHQ RAPKTRHIYHHPL HHRPDLSSHHRSS TMNPPPHTQPPG QPQPRPPIYSSHL | 34 | 0 | 0 | #N/A | #N/A | #N/A | #N/A | #N/A | #N/A |
| --- | --- | --- | --- | --- | --- | --- | --- | --- | --- | --- | --- |
| SB_0084 | SB_0084 | ITPAIMTLGHNMI YLHTSRDQPNPLR PCRRGVRTSLRLQ HRMRRRPLRPILH SRMHKHYYNKHP HHYNLPRNNMW RTLPWTLHNMFC HQDPTSNLPVLM NSNSMPPIPLRPT HTPPMKKLPTTHP SITYMMCLHTHY NLQHSPSNLRNM SDKRVTLME | 101 | 4 | 24.34214933 | 0.000999000999 | NMSDKRVTLME | 1322.632225 | 1626.841966 | -6.55855142 | No |
| SB_0085 | SB_0085 | INPLAQPVIYSTIFA GTLITALSSHWFFT WVGLEMNMLAFI PVLTKKMNPRSTE AAIKYFLTQATAS MILLMAILFNNML SGQWTMTNTTN QYSSLMIMMAM AMKLGMAPFHF WVPEVTQGTPLTS GLLLLTWQKLAPIS IMYQISPSLNVSLL LTLSILSIMAGSW GGLNQTQLRKILA YSSITHMGWMM AVLPYNPNMTILN LTIYIILTTTAFLLLN LNSSTTTLLLSRTW NKLTWLTPLIPSTL LSLGGLPPLTGFLP KWAIIEEFTKNNSL IIPTIMATITLLNLY FYLRLIYSTSITLLP MSNNVKMKWQF EHTKPTPFLPTLIAL TTLLLPISPFMLMI L | 68 | 17 | 30.98510606 | 0.001048218029 | WAIIEEFTK | 1135.591332 | 1423.796681 | -0.8391981488 | Yes |

| SB_0086 | SB_0086 | MFADRWLFSTNH KDIGTLYLLFGAW AGVLGTALSLLIRA ELGQPGNLLGND HIYNVIVTAHAFV MIFFMVMPIMIG GFGNWLVPLMIG APDMAFPRMNN MSFWLLPPSLLLLL ASAMVEAGAGTG WTVYPPLAGNYS HPGASVDLTIFSLH LAGVSSILGAINFIT TIINMKPPAMTQY QTPLFVWSVLITA VLLLLSLPVLAAGIT MLLTDRNLNTTFF DPAGGGDPILYQH LFWFFGHPEVYILI LPGFGMISHIVTYY SGKKEPFGYMGM VWAMMSIGFLGF IVWAHHMFTVG MDVDTRAYFTSA TMIIAIPTGVKVFS WLATLHGSNMK WSAAVLWALGFIF LFTVGGLTGIVLAN SSLDIVLHDTYYVV AHFHYVLSMGAV FAIMGGFIHWFPL FSGYTLDQTYAKIH FTIMFIGVNLTFFP QHFLGLSGMPRR YSDYPDAYTTWNI LSSVGSFISLTAVM LMIFMIWEAFASK RKVLMVEEPSMN LEWLYGCPPPYHT FEEPVYMKSRQKR KESNPPKLVSSQP HGLHDFFKKVLEK PFHNFVKVKL | 151 | 91 | 90.04728257 | 0.000999000999 | VLMVEEPSMNLE WLYGCPPPYHTFE EPVYMK | 3727.711022 | 4088.940668 | -2.215073813 | Yes |
| --- | --- | --- | --- | --- | --- | --- | --- | --- | --- | --- | --- |

| SB_0087 | SB_0087 | IMTNPENQNERK SVRFIHCPHNPRP TRRSTDHSISPSID PHLQMSHQQPTN HHPTMTNQTNLK TNDNHTQH | 31 | 0 | 0 | #N/A | #N/A | #N/A | #N/A | #N/A | #N/A |
| --- | --- | --- | --- | --- | --- | --- | --- | --- | --- | --- | --- |
| SB_0088 | SB_0088 | IYTNHPTIYKPSHG HPLMSGHSDYRLS L | 3 | 0 | 0 | #N/A | #N/A | #N/A | #N/A | #N/A | #N/A |

| SB_0089 | SB_0089 | MMTHQSHAYHM VKPSPWPLTGALS ALLMTSGLAMWF HFHSMTLLMLGLL TNTLTMYQWWR DVTRESTYQGHHT PPVQKGLRYGMIL FITSEVFFFAGFFW AFYHSSLAPTPQL GGHWPPTGITPL NPLEVPLLNTSVLL ASGVSITWAHHSL MENNRNQMIQA LLITILLGLYFTLLQ ASEYFESPFTISDGI YGSTFFVATGFHG LHVIIGSTFLTICFIR QLMFHFTSKHHF GFEAAAWYWHF VDVVWLFLYVSIY WWGSYSFSMNST VNFQLTSFDNIQK RVMNFALILMINT LLALLLMIITFWLP QLNGYMEKSTPYE CGFDPMSPARVP FSMKFFLVAITFLL FDLEIALLLPLPWA LQTTNLPLMVMS SLLLIIILALSLAYE WLQKGLDWTELV YSLNKTNDFDSLN YDNHIYQMPLIYM NIMLAFTISLLGML VYRSHLMSSLLCLE GMMLSLFIMATL MTLNTHSLLANIV PIAMLVFAACEAA VGLALLVSISNTYG LDYVHNLNLLQC | 111 | 69 | 55.11127489 | 0.000999000999 | ESTYQGHHTPPVQ K | 1607.769182 | 1895.97657 | -1.701906999 | Yes |
| --- | --- | --- | --- | --- | --- | --- | --- | --- | --- | --- | --- |
| SB_0090 | SB_0090 | MSSSKPHLSPPWL SSPDEATSQNAW TQAHTSYSTP | 0 | 0 | 0 | #N/A | #N/A | #N/A | #N/A | #N/A | #N/A |
| SB_0091 | SB_0091 | IVNLTTEAYDPLFT  EKAHKNC | 2 | 0 | 0 | #N/A | #N/A | #N/A | #N/A | #N/A | #N/A |

| SB_0092 | SB_0092 | MPPCLTTWLSQLL KDNSYPLVLGPKN FGATPNKSNNHA HYYNHPNPDFPN SPHPYHPR | 28 | 4 | 20.47544064 | 0.001998001998 | DNSYPLVLGPK | 1201.634257 | 1489.846087 | -5.150590851 | No |
| --- | --- | --- | --- | --- | --- | --- | --- | --- | --- | --- | --- |
| SB_0093 | SB_0093 | MPPLCKIHCRIHLY YQSLPHNNIHVPR PRSYYLELTLSHNP NNPALPKLQTRLL LHNIHPCSIVRYM VHHRILTVMYKLR PKH | 33 | 0 | 0 | #N/A | #N/A | #N/A | #N/A | #N/A | #N/A |
| SB_0094 | SB_0094 | IPTVHRLRGRRNYI LLAHQLMMRPSR CQHSSHSSNPMQ PYRRYRFHPRLSM IYPTLQLMRPTTN SPSKR | 40 | 0 | 0 | #N/A | #N/A | #N/A | #N/A | #N/A | #N/A |
| SB_0095 | SB_0095 | IRSPPLTPLSHRRP HPSLSPTPLKHYSC SRNLLTHPLPPPSR K | 14 | 0 | 0 | #N/A | #N/A | #N/A | #N/A | #N/A | #N/A |
| SB_0096 | SB_0096 | MLRRYHHSVRSSL RPYTKWHQKNRS LLHFKSTRTHNSY NRHQPTTPSIPAH LYPRLLQSHTIYVL RVHHPQP | 14 | 0 | 0 | #N/A | #N/A | #N/A | #N/A | #N/A | #N/A |
| SB_0097 | SB_0097 | ISRNTFPHRFLLQR PHHRNRKHIMHK RLSPIYYSHRYLPD KRL | 24 | 1 | 17.01163818 | 0.004842615012 | NRKHIMHK | 1062.586873 | 1494.885741 | 4.915416916 | No |
| SB_0098 | SB_0098 | MTYSPEQSQLQY MHQQTMFNQ | 4 | 0 | 0 | #N/A | #N/A | #N/A | #N/A | #N/A | #N/A |
| SB_0099 | SB_0099 | MMWKTIVVFQL QEHQWPQYAKLT P | 2 | 0 | 0 | #N/A | #N/A | #N/A | #N/A | #N/A | #N/A |
| SB_0100 | SB_0100 | IMAESSATFTPMA PQYSLSASSYTSGE AYITDHFSTQKPET  SALSSCLQL | 0 | 0 | 0 | #N/A | #N/A | #N/A | #N/A | #N/A | #N/A |
| SB_0101 | SB_0101 | IPMKSPSTLTTQSK TPSAYFSSFSP | 2 | 0 | 0 | #N/A | #N/A | #N/A | #N/A | #N/A | #N/A |
| SB_0102 | SB_0102 | IDSPINNRYVFRTL LPATMNIVRYHKY LTTCST | 11 | 0 | 0 | #N/A | #N/A | #N/A | #N/A | #N/A | #N/A |

| SB_0103 | SB_0103 | MLTSKYSNQPSTI THQLQLQSHPSPT RMPTNLPTLNST | 125 | 3 | 11.44115982 | 0.002997002997 | MPTNLPTLNST | 1187.585595 | 1347.676777 | 4.321420165 | Yes |
| --- | --- | --- | --- | --- | --- | --- | --- | --- | --- | --- | --- |
| SB_0104 | SB_0104 | MKPFTVHSTLQSN  PFSSPWMTPLR | 3 | 0 | 0 | #N/A | #N/A | #N/A | #N/A | #N/A | #N/A |
| SB_0105 | SB_0105 | MYYVLLRVGRFVG ILVGEGWLWSCS WCVMVEGWLLYL LVSMGRGFWCGL GFYVLQVVKYLWY RTMFMVAGSNVR NT | 151 | 5 | 21.9103728 | 0.000999000999 | TMFMVAGSNVR | 1211.57907 | 1355.671422 | 7.18439158 | No |
| SB_0106 | SB_0106 | MGESMLGWYPN LLPHERTENSLN | 4 | 0 | 0 | #N/A | #N/A | #N/A | #N/A | #N/A | #N/A |
| SB_0107 | SB_0107 | MAYEGCCYSCKQ EDNADVSGFWVE KWSVM | 5 | 0 | 0 | #N/A | #N/A | #N/A | #N/A | #N/A | #N/A |
| SB_0108 | SB_0108 | IGVKVADDSAMIY  VSSDVGDWWKG GWGVWWVVHG | 3 | 0 | 0 | #N/A | #N/A | #N/A | #N/A | #N/A | #N/A |
| SB_0109 | SB_0109 | MVFHIIGRGCSPC ENNDVCFVSVECG  FSNGVCGVFF | 2 | 1 | 13.14643154 | 0.006053268765 | MVFHIIGR | 971.5374648 | 1131.640437 | -5.273572206 | No |
| SB_0110 | SB_0110 | MYYSDGYWGVS WGMGVRGWGL GECFSGVSDGGRI GAVGERVWWGG GCGKL | 54 | 4 | 28.02515 | 0.000999000999 | IGAVGERVWWGG GCGK | 1630.803791 | 1976.033302 | -1.964915008 | Yes |
| SB_0111 | SB_0111 | ISSSYWLNIVCWC MYCNWDCSGE | 0 | 0 | 0 | #N/A | #N/A | #N/A | #N/A | #N/A | #N/A |
| SB_0112 | SB_0112 | MGLRRLCMMCLR FRWCGLWSRNL WGKVFLLMLGCQ WWGRLKWEVWF WVVLLFFEYLVHC | 23 | 0 | 0 | #N/A | #N/A | #N/A | #N/A | #N/A | #N/A |
| SB_0113 | SB_0113 | MVWLWRRRGYR CAGMLGVVGWC RL | 28 | 0 | 0 | #N/A | #N/A | #N/A | #N/A | #N/A | #N/A |
| SB_0114 | SB_0114 | MVLEFGLVGYFLL  GGGSGWVRRFLL QL | 8 | 0 | 0 | #N/A | #N/A | #N/A | #N/A | #N/A | #N/A |
| SB_0115 | SB_0115 | IGLICLLLLGGGLV  VGWGLD | 2 | 0 | 0 | #N/A | #N/A | #N/A | #N/A | #N/A | #N/A |

| SB_0116 | SB_0116 | MLRRGWNRYRRY GCMGLLEWLLCW HLLGRIINWWAR RM | 32 | 1 | 19.24635622 | 0.008474576271 | IINWWAR | 957.5184452 | 1101.610498 | 9.109554067 | No |
| --- | --- | --- | --- | --- | --- | --- | --- | --- | --- | --- | --- |
| SB_0117 | SB_0117 | MLQGWMLWRSS LVWSLGRAGLFGL WLSVSSR | 21 | 1 | 12.74455536 | 0.003996003996 | AGLFGLWLSVSSR | 1391.756098 | 1535.870039 | -7.710232259 | No |
| SB_0118 | SB_0118 | IETSRARPTAASQ AAKTSMAMGTM LAKREWVLRVMR VAMMNSDSIIPSR HSREDMRCERYTS IPRSEMVNASMM  FM | 146 | 2 | 20.03051746 | 0.000999000999 | VAMMNSDSIIPSR HSR | 1799.877029 | 1959.982881 | -4.506118554 | No |
| SB_0119 | SB_0119 | MMINKRDDMTIS GRLVVCRAHGRG KRRAISRSNNKKV MATKKNFMEKGT RAGDMGSKPHS | 64 | 4 | 19.56720182 | 0.002424242424 | DDMTISGR | 893.3912624 | 1053.484827 | 3.257502508 | Yes |
| SB_0120 | SB_0120 | MIISSKARRVLIIKI KAKFITLFWMLSK LVNWKLTVLFMLK E | 24 | 0 | 0 | #N/A | #N/A | #N/A | #N/A | #N/A | #N/A |
| SB_0121 | SB_0121 | METYRNSQTTSTK CQYQAAASKPKW CLDVKWNISWRM KQMVRKVEPMM TWSPWKPVATKN VEP | 24 | 0 | 0 | #N/A | #N/A | #N/A | #N/A | #N/A | #N/A |
| SB_0122 | SB_0122 | MPSEMVKGDSKY SEACRRVK | 32 | 1 | 13.0533318 | 0.006651884701 | YSEACRR | 883.3970162 | 1084.531222 | -9.830153414 | No |
| SB_0123 | SB_0123 | IVMSSAWIIWFRL FSIRLWWAQVIDT PDASNTDVFRSGT SRGFSGVMPVGG QCPPNWGVGARL EW | 10 | 0 | 0 | #N/A | #N/A | #N/A | #N/A | #N/A | #N/A |
| SB_0124 | SB_0124 | MNRIIPYRRPFWT GGVWWPWYVLS RVTSRHHWYMVS VLVSRPSMRSVM EWKWNHMARPE VIRRAERAPVRGH GLGFTMW | 113 | 4 | 19.15712093 | 0.000999000999 | HHWYMVSVLVSR PSMR | 1983.992327 | 2128.084213 | 4.805330945 | No |

| SB_0125 | SB_0125 | IMCCRAGRGLLEV WKRRLGLRRQRFL G | 15 | 1 | 25.62950155 | 0.000999000999 | IMCCRAGRGLLEV WK | 1733.889112 | 2152.138023 | -3.205659403 | Yes |
| --- | --- | --- | --- | --- | --- | --- | --- | --- | --- | --- | --- |
| SB_0126 | SB_0126 | IVGWWLVGCWW DIWRWGSMEGE MEWSVLRRVGLG LWGQWMKRTDF RSFWFSGFVMIFY  FYGLWWGR | 33 | 0 | 0 | #N/A | #N/A | #N/A | #N/A | #N/A | #N/A |
| SB_0127 | SB_0127 | MEGSSTIRTFRFEA KASQIMKIINITAV REMNEPTDDRMF HVVYASG | 89 | 4 | 17.13238898 | 0.009523809524 | IINITAVR | 898.5599734 | 1042.671773 | -9.311726812 | No |
| SB_0128 | SB_0128 | IPDRPRKCCGKKV RFTPMNMMVKW ILA | 8 | 0 | 0 | #N/A | #N/A | #N/A | #N/A | #N/A | #N/A |
| SB_0129 | SB_0129 | MSSDEFANTMPV RPPTVKRKMNPR AQSTAADHFMLL PWSVASQLNTLTP VGMAMIMVAEV KYARVSTSIPTVN MWCAHTMNPRK PIDIMAQTMPMY  PNGSFFPE | 14 | 0 | 0 | #N/A | #N/A | #N/A | #N/A | #N/A | #N/A |
| SB_0130 | SB_0130 | MGSPPPAGSKKV VLRLRSVSSMVM PAARTGRDRRSRT AVIRTDQTKRGV WYWVMAGGFM LMIVVMKLMAPK MEETPARCKEKM VRSTEAPGWE | 401 | 9 | 31.99770905 | 0.000999000999 | TDQTKRGVWYWV MAGGFMLMIVV MK | 2946.478001 | 3410.753217 | 6.120421135 | No |
| SB_0131 | SB_0131 | MADASRSRREGG KSQKLMLFMRGN AMSGAPIIRGTSQ LPKPPIMMGITMK KIITNAWAVTMTL | 53 | 3 | 11.91927806 | 0.009900990099 | MADASRSR | 892.418479 | 1036.514518 | 5.830421843 | No |
| SB_0132 | SB_0132 | MWSLPRRLPGWP SSARMRRLRAVPR TPAHAPNNRYSVP MSLWFVENSQRS ANISGGEVKWLSE ALDCKSKDRG | 107 | 5 | 24.68961083 | 0.002040816327 | WLSEALDCK | 1063.500807 | 1408.724693 | 1.226049523 | Yes |

| SB_0133 | SB_0133 | IELQIRRSSFKPAG ASPAFFPGGGRSR LKPVD | 4 | 0 | 0 | #N/A | #N/A | #N/A | #N/A | #N/A | #N/A |
| --- | --- | --- | --- | --- | --- | --- | --- | --- | --- | --- | --- |
| SB_0134 | SB_0134 | IKVADLRSVDAEW  GFAVLSCYRN | 9 | 0 | 0 | #N/A | #N/A | #N/A | #N/A | #N/A | #N/A |
| SB_0135 | SB_0135 | IISMKGEMGRSSV VRAMSVGRNGV GFVCSNCHFIFTLL DMGSSVIEVE | 38 | 1 | 10.14925663 | 0.007063572149 | IISMKGEMGR | 1120.573256 | 1408.789118 | -8.310300416 | No |
| SB_0136 | SB_0136 | MVAMMVGMMR LLFFVNSSMMAH LGKKPVSGGRPPR ERRVDGIKGVSHV SLFQVRDSSRVVV LEFKLSSRNAVVV RMM | 90 | 2 | 22.19791862 | 0.002688172043 | MVAMMVGMMR | 1155.509478 | 1347.595171 | 0.8468832668 | Yes |
| SB_0137 | SB_0137 | MVKLRMVMLGLY GRTAIIHPMWVIE EYAKILRSWVWFN PPQLPAMMDKIE RVRRRLTFSEGEI WYMIEMGASFCH VRRSRPDVRGVP WVTSGTQKWKG AIPSFIAMAIMIIN DEYWLVVLVMVH CPESMLLKRMAIR RIMDAVACVRKYL MAASVERGFIFLV RTGMKASMFISRP TQVKNQCELSAV MSVPAKMVE | 231 | 23 | 24.95221347 | 0.000999000999 | SWVWFNPPQLPA MMDK | 1945.921851 | 2250.102287 | 8.288127825 | No |
| SB_0138 | SB_0138 | MGCDRWHGEFW ILRDGFDSHSPRN KGV | 20 | 5 | 18.53962327 | 0.003144654088 | DGFDSHSPR | 1016.431152 | 1160.534518 | -1.106494702 | Yes |
| SB_0139 | SB_0139 | IIYSIKVTLLSDMFL  RFEGECWRL | 13 | 0 | 0 | #N/A | #N/A | #N/A | #N/A | #N/A | #N/A |
| SB_0140 | SB_0140 | MLGWVVGSFFM GGVWVGRSGIGG MLFEFMRTGRLE VGSWWQNMLCR VQGRVRHMLFLG RL | 23 | 0 | 0 | #N/A | #N/A | #N/A | #N/A | #N/A | #N/A |

| SB_0141 | SB_0141 | IRLWRMGRRGLR RIRCWSLRLVRTPL RQGRRGFGWSLL VWR | 34 | 5 | 24.18824918 | 0.008869179601 | TPLRQGR | 826.4773112 | 970.5764446 | 3.041597953 | Yes |
| --- | --- | --- | --- | --- | --- | --- | --- | --- | --- | --- | --- |
| SB_0142 | SB_0142 | MLMVEWWLGW LHMRLFGLLLAVR RSGRSLSLMLTLIR GLSKRLG | 10 | 0 | 0 | #N/A | #N/A | #N/A | #N/A | #N/A | #N/A |
| SB_0143 | SB_0143 | MGGLGWGWPG GWVWGGGFMV EERWWELRSGR WCRGWW | 12 | 0 | 0 | #N/A | #N/A | #N/A | #N/A | #N/A | #N/A |
| SB_0144 | SB_0144 | MWRVLGALWW RVLWRQRRVVVA RRGLQRWGLCVV VYSLEFFVR | 24 | 1 | 13.52718761 | 0.003816793893 | QRRVVVAR | 982.6148008 | 1126.70881 | 7.174189307 | No |
| SB_0145 | SB_0145 | MPLRLEWVQWG VGGWPWVCC | 0 | 0 | 0 | #N/A | #N/A | #N/A | #N/A | #N/A | #N/A |
| SB_0146 | SB_0146 | ITGLCHLNKPCSW VGVGMMLSWDD IIYGGRRFVK | 3 | 0 | 0 | #N/A | #N/A | #N/A | #N/A | #N/A | #N/A |
| SB_0147 | SB_0147 | MVVRFDWWSLS MYCSEVGFCSEVA  PTEIFNAGLVV | 0 | 0 | 0 | #N/A | #N/A | #N/A | #N/A | #N/A | #N/A |
| SB_0148 | SB_0148 | MGSSRLAVLCPPL HGQVNFTG | 1 | 0 | 0 | #N/A | #N/A | #N/A | #N/A | #N/A | #N/A |
| SB_0149 | SB_0149 | IHYAEGMGVSPCY IMLGYNFSSFPCG TMSIAPGFNFYRL YFIWVNGLAKVV W | 6 | 0 | 0 | #N/A | #N/A | #N/A | #N/A | #N/A | #N/A |
| SB_0150 | SB_0150 | MAVYRLSKRWW GWSGFIDYRTGSS RGMWSTARSFEF | 30 | 0 | 0 | #N/A | #N/A | #N/A | #N/A | #N/A | #N/A |
| SB_0151 | SB_0151 | MERLGPNLFVYG VMWARLNIFSVLL WGGKLHKLWGVS  LGFGWFGVWG | 24 | 1 | 11.74932761 | 0.000999000999 | MERLGPNLFVYGV MWAR | 2038.028042 | 2198.13785 | -5.814048991 | No |
| SB_0152 | SB_0152 | IFCYDVCVESGCA DIQLLLLCPTSIN | 0 | 0 | 0 | #N/A | #N/A | #N/A | #N/A | #N/A | #N/A |
| SB_0153 | SB_0153 | MMGWGRNQRQ MLRHRVLRLQRLA MLSRAYPPDENTK CMESSREWLMG W | 61 | 4 | 16.473808 | 0.004197271773 | MMGWGRNQR | 1134.517476 | 1310.619018 | -7.343252657 | No |

| SB_0154 | SB_0154 | MSDTVHFSYPQVL WARSEESSTLVRD IDFTEDGGQGTPI WGGSSMGTRRDL TVMCYVR | 75 | 3 | 25.31221444 | 0.001998001998 | DIDFTEDGGQGTPI WGGSSMGTR | 2383.038604 | 2527.126774 | 5.516739491 | No |
| --- | --- | --- | --- | --- | --- | --- | --- | --- | --- | --- | --- |
| SB_0155 | SB_0155 | MYYRWSSIYGTV QYSWWLAVMYE MHSGCWWVSQY LGGTQICFPMKEQ RMV | 1 | 0 | 0 | #N/A | #N/A | #N/A | #N/A | #N/A | #N/A |
| SB_0156 | SB_0156 | IRILALGANGGVK DFFSDLSLEKGFHL RFTRLVY | 22 | 1 | 24.23521817 | 0.001020408163 | DFFSDLSLEK | 1199.570992 | 1487.783158 | -5.386567532 | No |
| SB_0157 | SB_0157 | MLQGQAHLSILFSI REMVGIRIRIVVKY  STDATCPMMVKG | 99 | 1 | 16.12499284 | 0.003631961259 | EMVGIRIR | 972.5538426 | 1132.64155 | 8.208387675 | No |
| SB_0158 | SB_0158 | IMLCCLDMWRM GIIARMRMDSNR ARTPPSLLGTDRRI V | 81 | 12 | 20.26730827 | 0.001048218029 | MRMDSNRAR | 1135.533854 | 1295.622706 | 6.291403817 | No |
| SB_0159 | SB_0159 | ILSEWEVIPRGLFD PVSCKNRRWSVA RAAMMKGKMK WKVKNRVRVGLS TE | 33 | 0 | 0 | #N/A | #N/A | #N/A | #N/A | #N/A | #N/A |
| SB_0160 | SB_0160 | IHWTRSVPMYGM ADSKFVITVAPQN DIWPHGRT | 35 | 10 | 18.13464371 | 0.002997002997 | SVPMYGMADSK | 1184.520555 | 1488.711045 | 5.760069176 | No |
| SB_0161 | SB_0161 | MKAVAMVASRR  MMPMFQVSE | 21 | 2 | 20.80739396 | 0.008083140878 | AVAMVASRR | 959.5334428 | 1103.624632 | 9.876398944 | No |

| SB_0162 | SB_0162 | MARNSPVVIWRI RQAPRSEPKFHHA EMLDGVGRSMNE WLINFIRGLVLRIG VIGVLVVEMQRW FFMSLVVVVVRAR MMMYALFLLSVG LVMGFVGFSSKPS PIYGGLVLIVSGVV GCVIILNFGGGYM GLMVFLIYLGGM MVVFGYTTAMAI EEYPEAWGSGVE VLVSVLVGLAMEV GLVLWVKEYDGV VVVVNFNSVGSW MIYEGEGSGLIRE DPIGAGALYDYGR WLVVVTGWTLFV GVYIVIEIARGNRL CD | 88 | 2 | 17.30034468 | 0.000999000999 | FHHAEMLDGVGR | 1367.640422 | 1511.729762 | 8.436715496 | No |
| --- | --- | --- | --- | --- | --- | --- | --- | --- | --- | --- | --- |
| SB_0163 | SB_0163 | MGDCAVCDARVE SEYVGEMKCA | 15 | 1 | 16.17840283 | 0.005076142132 | VESEYVGEMK | 1169.527414 | 1617.826131 | 1.490421419 | Yes |
| SB_0164 | SB_0164 | MFEELINVWVWV YMSQWEFYDGPC NEQCYRDEYYGEV V | 0 | 0 | 0 | #N/A | #N/A | #N/A | #N/A | #N/A | #N/A |
| SB_0165 | SB_0165 | MMGVWVFFVRV NEGGKDGGN | 50 | 0 | 0 | #N/A | #N/A | #N/A | #N/A | #N/A | #N/A |
| SB_0166 | SB_0166 | MDSCYPLKVEKA MLLDMGAWVSS SCELSR | 11 | 0 | 0 | #N/A | #N/A | #N/A | #N/A | #N/A | #N/A |
| SB_0167 | SB_0167 | MRGRKPLLSDSQS DVLVKLYLQEENP VMMSGLRDRRR MGDRCMNMRVF SRVNEGFMLLM WWVSEPHCVVV  NM | 45 | 2 | 14.61955031 | 0.002997002997 | MGDRCMNMRVF SR | 1601.704692 | 1802.814079 | 7.862381391 | No |
| SB_0168 | SB_0168 | MLSPVSRRVMFD QENVVTSTESSPS RLMVGGKARLAR LARSHQKAISGSR VWSPWERIMMR LWVRS | 63 | 2 | 19.0610428 | 0.008484848485 | IMMRLWVR | 1103.60958 | 1263.71823 | -9.212354895 | No |

| SB_0169 | SB_0169 | IMRMTAPVKLQG VWMRMAVTTRA MWLIEEYAMSDF RSVCRRQMELVM IMPHRDSTRKG | 408 | 14 | 24.68111573 | 0.000999000999 | RQMELVMIMPHR | 1539.78359 | 1683.892152 | -3.836329152 | Yes |
| --- | --- | --- | --- | --- | --- | --- | --- | --- | --- | --- | --- |
| SB_0170 | SB_0170 | IKLLAQEFDSSWA VRVSSRMFSEPRV LWV | 18 | 0 | 0 | #N/A | #N/A | #N/A | #N/A | #N/A | #N/A |
| SB_0171 | SB_0171 | MISSVAVNVMIKE  ICREISMER | 12 | 0 | 0 | #N/A | #N/A | #N/A | #N/A | #N/A | #N/A |
| SB_0172 | SB_0172 | MIVRGRSQVVSIR RGVVRGSEEKVGE QLNRLLLIWLKNS RGMMLMIRLWV VVLIQIMCFLESHV SGSNMIVGTISFSI GVGLGYVRSLGH MCWRLRLVGLGP PLLRRRQRLVWQ | 155 | 1 | 12.06194623 | 0.009433962264 | GRSQVVSIR | 1000.577749 | 1144.676438 | 2.971509154 | Yes |
| SB_0173 | SB_0173 | MVLFLLGMVGR MWGVSDMLVFL EVRW | 12 | 0 | 0 | #N/A | #N/A | #N/A | #N/A | #N/A | #N/A |
| SB_0174 | SB_0174 | MSRNHSFCLNYM PIRFSLILFVVTHRP DLGLGWWLMRG  MT | 22 | 0 | 0 | #N/A | #N/A | #N/A | #N/A | #N/A | #N/A |
| SB_0175 | SB_0175 | MVKPHLQNASIRR RLRSQSDVWM | 6 | 1 | 14.2310863 | 0.000999000999 | MVKPHLQNASIRR R | 1704.968163 | 2009.156981 | 5.11088456 | No |
| SB_0176 | SB_0176 | MLSRRCRRKWW RETRSTLRLVGG | 1 | 0 | 0 | #N/A | #N/A | #N/A | #N/A | #N/A | #N/A |
| SB_0177 | SB_0177 | MLLMRVMRMCL GVGLLGDLAGWC LLGASALLIGG | 9 | 0 | 0 | #N/A | #N/A | #N/A | #N/A | #N/A | #N/A |
| SB_0178 | SB_0178 | MGLSRIEGLFGQV VCGGLGMCFLVL HRAIIGMWLVCW LVGLVWGALWSG SEITWLGRRSLGG LRGPLLGVMGWV LLYDRHVIGGSLCV  VVQVEAY | 48 | 10 | 26.94818027 | 0.008474576271 | RSLGGLR | 757.4558484 | 901.5583248 | -0.4346533091 | Yes |
| SB_0179 | SB_0179 | IFSWVMRNSVRS MGVIMVGHTVVF SWGISL | 11 | 0 | 0 | #N/A | #N/A | #N/A | #N/A | #N/A | #N/A |

| SB_0180 | SB_0180 | ISKIFRGINSRTMG MKLWFAPQISEH WP | 51 | 3 | 22.00463893 | 0.003636363636 | IFRGINSR | 961.5457216 | 1105.650919 | -2.811209079 | Yes |
| --- | --- | --- | --- | --- | --- | --- | --- | --- | --- | --- | --- |
| SB_0181 | SB_0181 | IASVFKPNVGTAH ECKTSCDVIIMRM GASIGSTTRLSTSR SRRSPGSRNNGGS  M | 26 | 1 | 17.90667062 | 0.001002004008 | RSPGSRNNGGSM | 1218.552341 | 1362.66105 | -4.858555349 | No |
| SB_0182 | SB_0182 | MRRDGRAMRTR MMAGRMVQTVS ISWASEMLVLVSF VVSVRKRAYRTRK QMRKMIMRAWS WKVMSSSMMGE VASCRPTCAACAI KMYRI | 73 | 7 | 17.2891006 | 0.008849557522 | TRMMAGR | 821.3999924 | 981.4877215 | 9.442820089 | No |
| SB_0183 | SB_0183 | MVFLMPFWKSH GGHGVGLKPALG GSIPSFFV | 4 | 0 | 0 | #N/A | #N/A | #N/A | #N/A | #N/A | #N/A |
| SB_0184 | SB_0184 | ILCMRVLRMCGR VGGIHMVTPGLW RVLLLLGLFASKRR LLKSWKLLMLLLLE KWMSLQMMGCF MWCMHRGSPSN VGAFRMGRESVV  GRKLDLRRWMW | 55 | 1 | 16.68926836 | 0.000999000999 | ESVVGRKLDLR | 1270.735698 | 1558.932275 | 4.864339307 | No |
| SB_0185 | SB_0185 | MGEISEWSLLWW QMQLLLMGHSGS GLQRSTCRVVRCL VMSLLMQCQSGH LRWKERWILGLRA LQQIISYCFRGVW RVS | 42 | 0 | 0 | #N/A | #N/A | #N/A | #N/A | #N/A | #N/A |
| SB_0186 | SB_0186 | MVLFFRSSKLQYG RLFRSLVG | 13 | 0 | 0 | #N/A | #N/A | #N/A | #N/A | #N/A | #N/A |
| SB_0187 | SB_0187 | MGVGMEWGLLL RRGRRRWCWGC GLLVV | 20 | 1 | 20.65512067 | 0.001 | MGVGMEWGLLLR | 1360.699513 | 1520.792566 | 2.603800845 | Yes |
| SB_0188 | SB_0188 | MGEVGLLWLGRI RRRGAFGIGLWQ  GVLYW | 5 | 0 | 0 | #N/A | #N/A | #N/A | #N/A | #N/A | #N/A |

| SB_0189 | SB_0189 | MRAGVGEREVRV RSLCCLCGETPYR GHRLLGELVSCQS LRLWWVLLWRRL LQMHGLWR | 43 | 2 | 25.21398491 | 0.006564551422 | AGVGEREVR | 971.5148174 | 1115.609647 | 6.50557754 | No |
| --- | --- | --- | --- | --- | --- | --- | --- | --- | --- | --- | --- |
| SB_0190 | SB_0190 | IFMLNCKFEEAAS NLPGLLPPFFPAA GEVDWSQLIRVLS C | 2 | 1 | 12.7887681 | 0.006651884701 | IFMLNCK | 867.434644 | 1212.650713 | 7.869148105 | No |
| SB_0191 | SB_0191 | ICVQLMQSGVLQS LAVTEIKYCNLLRA LKALGLYLT | 8 | 0 | 0 | #N/A | #N/A | #N/A | #N/A | #N/A | #N/A |
| SB_0192 | SB_0192 | IWAKSRLAGAGLL GRGGWMELRVLV MLACFRCEMVVG SWCWSLSWVVG  MR | 24 | 3 | 30.49572606 | 0.001004016064 | SRLAGAGLLGR | 1069.635595 | 1213.734458 | 2.661272284 | Yes |
| SB_0193 | SB_0193 | MLRFCVAGFGLIH LNCLLWWMRLRE WGEGLRLVRERF GMWLRWGLVFV MWEEAGRMSEG CLG | 65 | 1 | 19.80232168 | 0.000999000999 | LVRERFGMWLR | 1461.802668 | 1605.90347 | 0.8099306964 | Yes |
| SB_0194 | SB_0194 | MRLLAWGNTW WQLLWNEGLFF WLELE | 0 | 0 | 0 | #N/A | #N/A | #N/A | #N/A | #N/A | #N/A |
| SB_0195 | SB_0195 | MGPMAYLADLTL GWGVMGGTENF GFSGMGSILMVLE  MRGFKLLLFTLSK | 5 | 0 | 0 | #N/A | #N/A | #N/A | #N/A | #N/A | #N/A |
| SB_0196 | SB_0196 | MWDCLGYCSQC ADQGVVWVWCS PWSEDWVNG | 0 | 0 | 0 | #N/A | #N/A | #N/A | #N/A | #N/A | #N/A |
| SB_0197 | SB_0197 | MMSFTGEGALW SRPYFSCPFVQGGI  WX | 1 | 0 | 0 | #N/A | #N/A | #N/A | #N/A | #N/A | #N/A |
| SB_0198 | SB_0198 | MSWSNIEVVNPIV DMDSRMGLRCYP | 15 | 0 | 0 | #N/A | #N/A | #N/A | #N/A | #N/A | #N/A |
| SB_0199 | SB_0199 | ISLVKSKRQLNPRG AIHTGPYLRNKWL CYLCTVRVPRPLN MCHWAGGASNT GDARGDVFGKQA G | 28 | 0 | 0 | #N/A | #N/A | #N/A | #N/A | #N/A | #N/A |

| SB_0200 | SB_0200 | MLCLVMIFHLSLA VLYLLRQVSISIAYT LFG | 1 | 0 | 0 | #N/A | #N/A | #N/A | #N/A | #N/A | #N/A |
| --- | --- | --- | --- | --- | --- | --- | --- | --- | --- | --- | --- |
| SB_0201 | SB_0201 | MLEEGDGRCVRA SGPCSTKHSTLSLL LNPPSTLKFHKGY  RSFLG | 28 | 3 | 12.21386031 | 0.004243281471 | MLEEGDGR | 905.3912624 | 1065.481706 | 6.149467885 | No |
| SB_0202 | SB_0202 | MGWARGGEVDR GLSITEQAPLEGYE APPGPLSFKLWLV VFWRAVLLI | 13 | 1 | 17.17685495 | 0.000999000999 | MGWARGGEVDR | 1232.572012 | 1376.671919 | 1.585656856 | Yes |
| SB_0203 | SB_0203 | MNCGGCLWGLV GSGYGVSSGVCVL GRMGGGCIDEISS MGVGGENNVLV GGWLLKVHTAKR | 7 | 0 | 0 | #N/A | #N/A | #N/A | #N/A | #N/A | #N/A |
| SB_0204 | SB_0204 | MCLSAVARSGGG GVWWKFFVMMS VWKVAVQTFNCY YYVLQALIN | 18 | 0 | 0 | #N/A | #N/A | #N/A | #N/A | #N/A | #N/A |
| SB_0205 | SB_0205 | MGCSGSSVSQCY RVHTPQTKMPNA WRAPVSG | 21 | 6 | 11.77834789 | 0.001998001998 | MGCSGSSVSQCYR | 1363.531865 | 1621.660993 | 9.797445322 | No |
| SB_0206 | SB_0206 | MSYLRGTCGLFRL YDPEVGTRCRMQ FTLATPKCYGPGA RRVALLCGMLISR RMVVKGPLSEGG  HPWGREGIWL | 119 | 6 | 33.32383618 | 0.000999000999 | VALLCGMLISR | 1174.656587 | 1391.768589 | 4.651040481 | No |
| SB_0207 | SB_0207 | MCDSWGLIAVLA CKHGEGVLMWIG FLCTTGGQVFMV PYNIHGGWQ | 13 | 0 | 0 | #N/A | #N/A | #N/A | #N/A | #N/A | #N/A |
| SB_0208 | SB_0208 | MAVVDGWVNT WVVPKSASPWKN RE | 11 | 0 | 0 | #N/A | #N/A | #N/A | #N/A | #N/A | #N/A |
| SB_0209 | SB_0209 | MVELKTFSLICPW KKVFISGLQDWCI SLYYKDRPIWVFC  FQLGR | 30 | 0 | 0 | #N/A | #N/A | #N/A | #N/A | #N/A | #N/A |
| SB_0210 | SB_0210 | IGLVGEMLCFVV WMYGGWGLLLG WGWMVMGQGR LLVC | 0 | 0 | 0 | #N/A | #N/A | #N/A | #N/A | #N/A | #N/A |

| SB_0211 | SB_0211 | MGNIIRAWCGEG CLRGWLGYNCLG RLGGLVRMVLMS LRREGREVSRGRL WLCSKGGRWFYR NGRWFLGGCLIPF RARMGGGVLLGL Q | 182 | 5 | 19.94206225 | 0.000999000999 | MVLMSLRREGR | 1346.727458 | 1506.831371 | -4.578775128 | No |
| --- | --- | --- | --- | --- | --- | --- | --- | --- | --- | --- | --- |
| SB_0212 | SB_0212 | MLRRHWREGSG WFSHNLRLEWCG RLMKRRLRRLVSS AWLGMVLWWF GGSGRRQGVSRS FIMRRCWMGWG GRWMSG | 89 | 2 | 12.08128351 | 0.008083140878 | RCWMGWGGR | 1107.485449 | 1308.600806 | 6.261558495 | No |
| SB_0213 | SB_0213 | MLQRWLLRSILRH GGQGLRSWWVF | 2 | 0 | 0 | #N/A | #N/A | #N/A | #N/A | #N/A | #N/A |
| SB_0214 | SB_0214 | MKERGQGWFGRI LLVRGLCMIMGV D | 16 | 0 | 0 | #N/A | #N/A | #N/A | #N/A | #N/A | #N/A |
| SB_0215 | SB_0215 | IMPFWVEVMME VEIWCCEIVLGNSF SSQVRSRRSRGRF  WLVRRPR | 12 | 0 | 0 | #N/A | #N/A | #N/A | #N/A | #N/A | #N/A |
| SB_0216 | SB_0216 | IVVWKGDAGEML LVMRNPANRLPA ARRLMGFSRVGLF SLMLVRVGKRGW PVRVRRIIRVL | 77 | 1 | 18.13639941 | 0.000999000999 | GDAGEMLLVMR | 1190.578736 | 1334.687841 | -5.255125499 | No |
| SB_0217 | SB_0217 | MRVMDRAQAFV YDMFAVSMMWS LE | 23 | 0 | 0 | #N/A | #N/A | #N/A | #N/A | #N/A | #N/A |

| SB_0218 | SB_0218 | IPANARLPMVREV EVRGMVLSSPPIF RMSCSLLRLWM MDPEHMNSMAL KKAWVQMCRNA RCGWLMPIVTIM SPSWLEVEKATIFL MSFCVRAQTAAN RVVMAPKHSVRV WISGLFSARGWK RMSKKIPATTMVL EWSRAETGVGPS MAEGSQGWRPN WADLPAAARRRP SSGVRLGLAFRRAI CCGSHELECRMN HAKARMKPMSP MRLYRIAWMAAV LASARAYHQLMSK KDMIPTPSQPMN SWNRLLAVTKISM VIRKMSRYLKNWL MFGSEFMYHSEN SMMDHVTNNAT GMNIMEK | 254 | 11 | 41.69602147 | 0.000999000999 | EVEVRGMVLSSPPI FR | 1814.971241 | 1975.083261 | -7.59190579 | No |
| --- | --- | --- | --- | --- | --- | --- | --- | --- | --- | --- | --- |
| SB_0219 | SB_0219 | MMTSWSRHMNI VVGKRLMMKVD ATMDFT | 31 | 5 | 16.35315213 | 0.001210653753 | HMNIVVGK | 896.4901828 | 1200.683129 | 5.095019283 | No |
| SB_0220 | SB_0220 | MGGIREVRVRVV MVVCMVITFIWS  CTKIFGA | 10 | 0 | 0 | #N/A | #N/A | #N/A | #N/A | #N/A | #N/A |
| SB_0221 | SB_0221 | MGGEWGMGVW TWGCFLVWMRV LCC | 13 | 0 | 0 | #N/A | #N/A | #N/A | #N/A | #N/A | #N/A |
| SB_0222 | SB_0222 | MGMVQGRGRLC VLSGGWEWVWG VLYHSRLVLRVLR QVLLTQRWGLRH  GL | 15 | 3 | 17.1813702 | 0.006928406467 | MGMVQGRGR | 990.485115 | 1150.574672 | 6.469627156 | No |
| SB_0223 | SB_0223 | MLSCWLRSLMVL GQWEWVVECLVS LGCCECKLVRWV GEGSLLGCRMGS MCLRSGVLAGCLI GWW | 52 | 0 | 0 | #N/A | #N/A | #N/A | #N/A | #N/A | #N/A |
| SB_0224 | SB_0224 | MVVHWMSGVGL PWLWGVGVR | 9 | 0 | 0 | #N/A | #N/A | #N/A | #N/A | #N/A | #N/A |

| SB_0225 | SB_0225 | MPVSGGGFEAKV MFGCKVKY | 35 | 1 | 17.54348204 | 0.000999000999 | MPVSGGGFEAK | 1078.511705 | 1382.713954 | -2.302300023 | Yes |
| --- | --- | --- | --- | --- | --- | --- | --- | --- | --- | --- | --- |
| SB_0226 | SB_0226 | IKATAISRMVSRIRI VKMMSVEGRLM VDIARVALPIRCM SRWPAVMLAVRR TARAIGWMSRLM VSMMTSMGMRG VGVPCGKKWARA FLILERKPMITVPA HKGMAMARFMD SWVVGVNEWGR SPRRLVVAMKMIK DTSMRDQVRPLV LCMVIICFEVSLISH CWVVISRLLMRYL EVGINRGGNRMIS TAAGRPRIVGAM NEANRFSFILVLRV CYNFLFLWALVRE VGGSLCLMFLVG WWGMV | 515 | 22 | 34.85655731 | 0.000999000999 | AIGWMSRLMVSM MTSMGMRGVGV PCGK | 2872.353417 | 3233.551855 | 6.845155335 | No |
| SB_0227 | SB_0227 | MRALFQRFLGELIL GRWAWNCGLLH RFQSIDRSMPPVV | 12 | 0 | 0 | #N/A | #N/A | #N/A | #N/A | #N/A | #N/A |
| SB_0228 | SB_0228 | MMGEVCRSWRL VRRSRCTRRFSTIG  GQLIWW | 11 | 0 | 0 | #N/A | #N/A | #N/A | #N/A | #N/A | #N/A |
| SB_0229 | SB_0229 | MEAMGLAWNQL WGVRFLPFLSRFY VYGFFECVVGWG ASM | 8 | 0 | 0 | #N/A | #N/A | #N/A | #N/A | #N/A | #N/A |
| SB_0230 | SB_0230 | IASVECGESAKYFD AGGDSDDYGSGG EMCSCVYVYSYCK YMVCSHDKP | 8 | 0 | 0 | #N/A | #N/A | #N/A | #N/A | #N/A | #N/A |
| SB_0231 | SB_0231 | IANSKKQLQTCRG FSRLFSRRREK | 36 | 12 | 21.06855371 | 0.00218579235 | GFSRLFSRR | 1124.62028 | 1268.717786 | 3.615045629 | Yes |
| SB_0232 | SB_0232 | IGSIGYGSLSGEYIV EEDSY | 1 | 0 | 0 | #N/A | #N/A | #N/A | #N/A | #N/A | #N/A |
| SB_0233 | SB_0233 | MLEIVMGMETYH MSNARVSGRKFF HRRCMSWS | 53 | 0 | 0 | #N/A | #N/A | #N/A | #N/A | #N/A | #N/A |

| SB_0234 | SB_0234 | IVVVRVFIMMMF VYSAMKNRAKGP AAYSMLKPETSSD SPSARSKGVRLVS ASVEMNHIMAKG HDGRSNQRCSCV VMRVERLKEPLIS NVDSRMMARVTS YEIVWATARSAPI RA | 165 | 4 | 21.622387 | 0.000999000999 | LVSASVEMNHIMA K | 1528.774131 | 1816.968939 | 5.149099863 | No |
| --- | --- | --- | --- | --- | --- | --- | --- | --- | --- | --- | --- |
| SB_0235 | SB_0235 | MNRRPRLRLTRGL GMGRGVHSRRA MVRAKVGAVM | 31 | 2 | 14.75986827 | 0.005020080321 | GVHSRRAMVR | 1167.640697 | 1311.744847 | -1.567282974 | Yes |
| SB_0236 | SB_0236 | MVDVAGFRGSLV  KSFMASAKGCSSP | 17 | 1 | 16.69596443 | 0.008484848485 | MVDVAGFR | 893.4429008 | 1053.531901 | 7.592758242 | No |
| SB_0237 | SB_0237 | IFRSVSIRNAIAIR MGTMRSRRLAM GMLLRRGIEPLTV KF | 65 | 3 | 20.62049424 | 0.005773672055 | RLAMGMLLR | 1059.604495 | 1235.707948 | -9.330817665 | No |
| SB_0238 | SB_0238 | INKLKLHRVFSSCC VMPASSRAGQFH WLKVRDSWTLVE PFMQVPI | 47 | 3 | 15.62436483 | 0.00218579235 | AGQFHWLK | 985.5133602 | 1273.722598 | -3.994077948 | Yes |
| SB_0239 | SB_0239 | MLVMLEVMFLVN RRGKICRVPFTFFN LSLWACLCWVDS EGNNDLLVDCRY WAVNCQFSVLIW RRLMRRRMFSCY LY | 47 | 0 | 0 | #N/A | #N/A | #N/A | #N/A | #N/A | #N/A |
| SB_0240 | SB_0240 | MDWSNWVWGV QLYVWDFLGSGC WAWTLS | 0 | 0 | 0 | #N/A | #N/A | #N/A | #N/A | #N/A | #N/A |
| SB_0241 | SB_0241 | MGVKFFTLSTRFF PSVQRAVPLWTN  S | 4 | 1 | 13.41600805 | 0.008403361345 | FFTLSTR | 870.4599302 | 1014.555688 | 6.236398252 | No |
| SB_0242 | SB_0242 | MSFEVYLRRVTGG VYALQGPVQLSTL LLVYC | 5 | 0 | 0 | #N/A | #N/A | #N/A | #N/A | #N/A | #N/A |
| SB_0243 | SB_0243 | MRAIVVFWGRKC SPFLATSWATPW  PNVFTWVLALTL | 2 | 0 | 0 | #N/A | #N/A | #N/A | #N/A | #N/A | #N/A |
| SB_0244 | SB_0244 | MVGYLIPVWVLAI VCSDMLKPLS | 0 | 0 | 0 | #N/A | #N/A | #N/A | #N/A | #N/A | #N/A |
| SB_0245 | SB_0245 | ILCQLEFFTTQVSF SFIGEGVI | 3 | 0 | 0 | #N/A | #N/A | #N/A | #N/A | #N/A | #N/A |

| SB_0246 | SB_0246 | IAACLMLVPFDRG DLEGELTGTGMLA CVILLRANRKARTK PICLWGDVSPSKH FQCIALRR | 32 | 0 | 0 | #N/A | #N/A | #N/A | #N/A | #N/A | #N/A |
| --- | --- | --- | --- | --- | --- | --- | --- | --- | --- | --- | --- |
| SB_0247 | SB_0247 | MGLAAVCVCWV GWAGVVLMRLV VWEWEGKMMC | 21 | 0 | 0 | #N/A | #N/A | #N/A | #N/A | #N/A | #N/A |
| SB_0248 | SB_0248 | MPPKDKIWNLVR LVLGFFVFGVWQ  RCV | 22 | 0 | 0 | #N/A | #N/A | #N/A | #N/A | #N/A | #N/A |
| SB_0249 | SB_0249 | MLNVGAMNNR MRQESKTDTAT | 22 | 1 | 17.63291836 | 0.002030456853 | MLNVGAMNNR | 1118.532457 | 1262.64688 | -9.768674111 | No |

**Filtered list based on validation (only "Yes")**

| **Prot_Acces**  **sion** | **Description** | **Prot_Seq** | **PepQuery_**  **hits** | **Confident_**  **hits** | **Best_Score** | **pvalue** | **Peptide_Se**  **quence** | **Peptide_Chec**  **k_Mass** | **Exp_Pep_Mas**  **s** | **Error_mass_p**  **pm** | **Validation** | **mt gene or**  **region** | **LEN(prot)** |
| --- | --- | --- | --- | --- | --- | --- | --- | --- | --- | --- | --- | --- | --- |
| SB_0086 | SB_0086 | MFADRWLFSTNHKDIGTLYLLFGAWAGVL GTALSLLIRAELGQPGNLLGNDHIYNVIVTA HAFVMIFFMVMPIMIGGFGNWLVPLMIG APDMAFPRMNNMSFWLLPPSLLLLLASAM VEAGAGTGWTVYPPLAGNYSHPGASVDLTI FSLHLAGVSSILGAINFITTIINMKPPAMTQY QTPLFVWSVLITAVLLLLSLPVLAAGITMLLT DRNLNTTFFDPAGGGDPILYQHLFWFFGHP EVYILILPGFGMISHIVTYYSGKKEPFGYMG MVWAMMSIGFLGFIVWAHHMFTVGMDV DTRAYFTSATMIIAIPTGVKVFSWLATLHGS NMKWSAAVLWALGFIFLFTVGGLTGIVLAN SSLDIVLHDTYYVVAHFHYVLSMGAVFAIM GGFIHWFPLFSGYTLDQTYAKIHFTIMFIGV NLTFFPQHFLGLSGMPRRYSDYPDAYTTW NILSSVGSFISLTAVMLMIFMIWEAFASKRK VLMVEEPSMNLEWLYGCPPPYHTFEEPVY MKSRQKRKESNPPKLVSSQPHGLHDFFKKV  LEKPFHNFVKVKL | 151 | 91 | 90.04728257 | 0.000999000999 | VLMVEEPS MNLEWLY GCPPPYHT FEEPVYMK | 3727.711022 | 4088.940668 | -2.215073813 | Yes | RefProt COX1 | 553 |
| SB_0063 | SB_0063 | MLKLIVPTIMLLPLTWLSKKHMIWINTTTHS LIISIIPLLFFNQINNNLFSCSPTFSSDPLTTPLL MLTTWLLPLTIMASQRHLSSEPLSRKKLYLS MLISLQISLIMTFTATELIMFYIFFETTLIPTLAI ITRWGNQPERLNAGTYFLFYTLVGSLPLLIAL IYTHNTLGSLNILLLTLTAQELSNSWANNLM WLAYTMAFMVKMPLYGLHLWLPKAHVEA PIAGSMVLAAVLLKLGGYGMMRLTLILNPLT KHMAYPFLVLSLWGMIMTSSICLRQTDLKS LIAYSSISHMALVVTAILIQTPWSFTGAVILMI AHGLTSSLLFCLANSNYERTHSRIMILSQGL QTLLPLMAFWWLLASLANLALPPTINLLGEL SVLVTTFSWSNITLLLTGLNMLVTALYSLYM FTTTQWGSLTHHINNMKPSFTRENTLMFM HLSPILLLSLNPDIITGFSSCKYSLTKTSDCESD NRGLRPLIYRESSQELLTHAPMSNNMAFST  FKG | 87 | 8 | 84.24539063 | 0.000999000999 | AHVEAPIA GSMVLAAV LLK | 1889.080786 | 2177.288583 | -1.661575297 | Yes | RefProt ND4 | 506 |
| SB_0056 | SB_0056 | MAHAAQVGLQDATSPIMEELITFHDHALMI IFLICFLVLYALFLTLTTKLTNTNISDAQEMET VWTILPAIILVLIALPSLRILYMTDEVNDPSLTI KSIGHQWYWTYEYTDYGGLIFNSYMLPPLF LEPGDLRLLDVDNRVVLPIEAPIRMMITSQD VLHSWAVPTLGLKTDAIPGRLNQTTFTATR PGVYYGQCSEICGANHSFMPIVLELIPLKIFE  MGPVFTL | 779 | 656 | 76.27685888 | 0.000999000999 | MMITSQD VLHSWAVP TLGLK | 2226.154024 | 2514.353794 | 1.755039781 | Yes | RefProt COX2 | 227 |

| SB_0024 | SB_0024 | ILVQLQMKVMTMHTTMTTLTLTSLIPPILTT LVNPNKKNSYPHYVKSIVASTFIISLFPTTMF MCLDQEVIISNWHWATTQTTQLSLSFKLDY FSMMFIPVALFVTWSIMEFSLWYMNSDPN INQFFKYLLIFLITMLILVTANNLFQLFIGWEG VGIMSFLLISWWYARADANTAAIQAILYNRI GDIGFILALAWFILHSNSWDPQQMALLNA NPSLTPLLGLLLAAAGKSAQLGLHPWLPSA MEGPTPVSALLHSSTMVVAGIFLLIRFHPLA ENSPLIQTLTLCLGAITTLFAAVCALTQNDIK KIVAFSTSSQLGLMMVTIGINQPHLAFLHIC THAFFKAMLFMCSGSIIHNLNNEQDIRKMG GLLKTMPLTSTSLTIGSLALAGMPFLTGFYSK DHIIETANMSYTNAWALSITLIATSLTSAYST RMILLTLTGQPRFPTLTNINENNPTLLNPIKR LAAGSLFAGFLITNNISPASPFQTTIPLYLKLT ALAVTFLGLLTALDLNYLTNKLKMKSPLCTFY FSNMLGFYPSITHRTIPYLGLLTSQNLPLLLLD LTWLEKLLPKTISQHQISTSIITSTQKGMIKLY  FLSFFFPLILTLLLIT | 113 | 63 | 76.17222727 | 0.000999000999 | FPTLTNINE NNPTLLNPI KR | 2308.253868 | 2596.46168 | -1.396327951 | Yes | RefProt ND5 | 612 |
| --- | --- | --- | --- | --- | --- | --- | --- | --- | --- | --- | --- | --- | --- |
| SB_0057 | SB_0057 | MPQLNTTVWPTMITPMLLTLFLITQLKMLN  TNYHLPPSPKPMKMKNYNKPWEPKWTKIC SLHSLPPQS | 328 | 277 | 64.10598268 | 0.000999000999 | MLNTNYHL PPSPKPMK | 1866.948398 | 2299.260345 | -2.484358335 | Yes | RefProt ATP8 | 68 |
| SB_0089 | SB_0089 | MMTHQSHAYHMVKPSPWPLTGALSALLM TSGLAMWFHFHSMTLLMLGLLTNTLTMYQ WWRDVTRESTYQGHHTPPVQKGLRYGMI LFITSEVFFFAGFFWAFYHSSLAPTPQLGGH WPPTGITPLNPLEVPLLNTSVLLASGVSITW AHHSLMENNRNQMIQALLITILLGLYFTLLQ ASEYFESPFTISDGIYGSTFFVATGFHGLHVII GSTFLTICFIRQLMFHFTSKHHFGFEAAAWY WHFVDVVWLFLYVSIYWWGSYSFSMNSTV NFQLTSFDNIQKRVMNFALILMINTLLALLL MIITFWLPQLNGYMEKSTPYECGFDPMSPA RVPFSMKFFLVAITFLLFDLEIALLLPLPWAL QTTNLPLMVMSSLLLIIILALSLAYEWLQKGL DWTELVYSLNKTNDFDSLNYDNHIYQMPLI YMNIMLAFTISLLGMLVYRSHLMSSLLCLEG MMLSLFIMATLMTLNTHSLLANIVPIAMLV FAACEAAVGLALLVSISNTYGLDYVHNLNLL  QC | 111 | 69 | 55.11127489 | 0.000999000999 | ESTYQGHH TPPVQK | 1607.769182 | 1895.97657 | -1.701906999 | Yes | RefProt COX3 | 520 |
| SB_0017 | SB_0017 | MNENLFASFIAPTILGLPAAVLIILFPPLLIPTS KYLINNRLITTQQWLIKLTSKQMMTMHNTK GRTWSLMLVSLIIFIATTNLLGLLPHSFTPTT QLSMNLAMAIPLWAGTVIMGFRSKIKNAL AHFLPQGTPTPLIPMLVIIETISLLIQPMALAV RLTANITAGHLLMHLIGSATLAMSTINLPSTL  IIFTILILLTILEIAVALIQAYVFTLLVSLYLHDNT | 130 | 89 | 51.46183017 | 0.001020408163 | LITTQQWLI K | 1242.733572 | 1530.93992 | -1.428598097 | Yes | RefProt ATP6 | 226 |
| SB_0005 | SB_0005 | MPPSSANPDEGYKVSASTHVKTLGQGVAH EVARNGLHFLPQKTTMALMKLKGRRWI | 111 | 8 | 31.85069597 | 0.001998001998 | TLGQGVAH EVAR | 1236.657452 | 1380.764859 | -3.848103668 | Yes | 12S sens  (chevauche MOTS-c | 56 |

| SB_0085 | SB_0085 | INPLAQPVIYSTIFAGTLITALSSHWFFTWVG LEMNMLAFIPVLTKKMNPRSTEAAIKYFLTQ ATASMILLMAILFNNMLSGQWTMTNTTN QYSSLMIMMAMAMKLGMAPFHFWVPEV TQGTPLTSGLLLLTWQKLAPISIMYQISPSLN VSLLLTLSILSIMAGSWGGLNQTQLRKILAYS SITHMGWMMAVLPYNPNMTILNLTIYIILTT TAFLLLNLNSSTTTLLLSRTWNKLTWLTPLIP STLLSLGGLPPLTGFLPKWAIIEEFTKNNSLIIP TIMATITLLNLYFYLRLIYSTSITLLPMSNNVK MKWQFEHTKPTPFLPTLIALTTLLLPISPFML  MIL | 68 | 17 | 30.98510606 | 0.001048218029 | WAIIEEFTK | 1135.591332 | 1423.796681 | -0.8391981488 | Yes | RefProt ND2 | 347 |
| --- | --- | --- | --- | --- | --- | --- | --- | --- | --- | --- | --- | --- | --- |
| SB_0192 | SB_0192 | IWAKSRLAGAGLLGRGGWMELRVLVMLAC  FRCEMVVGSWCWSLSWVVGMR | 24 | 3 | 30.49572606 | 0.001004016064 | SRLAGAGLL  GR | 1069.635595 | 1213.734458 | 2.661272284 | Yes | ND2, antisens | 50 |
| SB_0110 | SB_0110 | MYYSDGYWGVSWGMGVRGWGLGECFSG VSDGGRIGAVGERVWWGGGCGKL | 54 | 4 | 28.02515 | 0.000999000999 | IGAVGERV WWGGGC GK | 1630.803791 | 1976.033302 | -1.964915008 | Yes | ND6 sens (ND6 est sur autre brin),  frameShift | 50 |
| SB_0007 | SB_0007 | MVGRFMGRGDKPTEPGDSWLSKMES | 54 | 12 | 27.52984605 | 0.001998001998 | MVGRFMG  RGDKPTEP GDSWLSK | 2450.183426 | 2914.478541 | 0.3292530069 | Yes | 16S sens | 25 |
| SB_0178 | SB_0178 | **MGLSRIEGLFGQVVCGGLGMCFLVLHRAII GMWLVCWLVGLVWGALWSGSEITWLGR RSLGGLRGPLLGVMGWVLLYDRHVIGGSL**  **CVVVQVEAY** | 48 | 10 | 26.94818027 | 0.008474576271 | RSLGGLR | 757.4558484 | 901.5583248 | -0.4346533091 | Yes | COX3,  antisens | 95 |
| SB_0055 | SB_0055 | MICCSALSPRIHLSFHRRWPDWHCISKLITR HRTTRHVLRCSPLPLCPINRSCICHHRRLHSLI SPILRLHPRPNLRQNPFHYHIHRRKSNFLPTT LSRPIRNAPTLLGLPRCMHHMKHPIICRLIHF SNSSNINNFHDLRSLRFEAKSPNSRRTLHKP  GVTMWMPPTLPHIRRTRMHKI | 99 | 8 | 26.39510071 | 0.000999000999 | TTRHVLRCS PLPLCPINR | 2075.124397 | 2333.278658 | -3.949405601 | Yes | COX1 sens, frameShift | 180 |
| SB_0125 | SB_0125 | IMCCRAGRGLLEVWKRRLGLRRQRFLG | 15 | 1 | 25.62950155 | 0.000999000999 | IMCCRAGR  GLLEVWK | 1733.889112 | 2152.138023 | -3.205659403 | Yes | ATP6, antisens | 27 |
| SB_0132 | SB_0132 | **MWSLPRRLPGWPSSARMRRLRAVPRTPA HAPNNRYSVPMSLWFVENSQRSANISGG EVKWLSEALDCKSKDRG** | 107 | 5 | 24.68961083 | 0.002040816327 | WLSEALDC K | 1063.500807 | 1408.724693 | 1.226049523 | Yes | Non-codant (antisens entre nd2 et  cox1) | 73 |
| SB_0169 | SB_0169 | **IMRMTAPVKLQGVWMRMAVTTRAMWL**  **IEEYAMSDFRSVCRRQMELVMIMPHRDST RKG** | 408 | 14 | 24.68111573 | 0.000999000999 | RQMELVMI MPHR | 1539.78359 | 1683.892152 | -3.836329152 | Yes | ND4, antisens | 58 |
| SB_0058 | SB_0058 | MTPNRGPLSPPNDLRPSHVISLPLHNAPHT RPTNQHTNHMPMMARCNTRKHMPRPPH TTCPKRPSMRDNPIYYLRSFFLRRIFLSLLPLQ PSPYPPIRRALAPNRHHPAKSPRSPTPKHIRI  TRIRSINHLSSP | 35 | 2 | 24.59374409 | 0.000999000999 | RPSMRDNP IYYLR | 1679.856552 | 1968.066962 | -3.172199407 | Yes | COX3 sens, frameShift | 134 |
| SB_0141 | SB_0141 | IRLWRMGRRGLRRIRCWSLRLVRTPLRQGR  RGFGWSLLVWR | 34 | 5 | 24.18824918 | 0.008869179601 | TPLRQGR | 826.4773112 | 970.5764446 | 3.041597953 | Yes | ND1, antisens | 41 |
| SB_0008 | SB_0008 | ILLRMSLRQIKTLNWQLTAQYLQSTNKSLLP SLSTQHRHAHKERLKKVKGTRQILPRLFTKNI TSSITSIRGTACPVTHV | 43 | 1 | 23.99545008 | 0.000999000999 | LFTKNITSSI TSIR | 1579.893313 | 1868.103082 | -2.998073063 | Yes | 16S sens, complémentai re à SHLP1 et  SHLP4 | 80 |
| SB_0136 | SB_0136 | MVAMMVGMMRLLFFVNSSMMAHLGKK  PVSGGRPPRERRVDGIKGVSHVSLFQVRDS SRVVVLEFKLSSRNAVVVRMM | 90 | 2 | 22.19791862 | 0.002688172043 | MVAMMV GMMR | 1155.509478 | 1347.595171 | 0.8468832668 | Yes | ND2, antisens | 77 |

| SB_0180 | SB_0180 | ISKIFRGINSRTMGMKLWFAPQISEHWP | 51 | 3 | 22.00463893 | 0.003636363636 | IFRGINSR | 961.5457216 | 1105.650919 | -2.811209079 | Yes | COX2,  antisens | 28 |
| --- | --- | --- | --- | --- | --- | --- | --- | --- | --- | --- | --- | --- | --- |
| SB_0003 | SB_0003 | IIFPSHSHTTNLINTTPAHPTQHTHTAANPM PRTNQTPKTPPTVYVAYLLKAMHWKCLDG LTSPHKQMGLVLAFLLALSKITHASIPVPVSS PSKSPRSKGTSIKHAAMQLKTLSLATPPRET  AVINL | 52 | 3 | 21.5925048 | 0.008743169399 | TLSLATPPR | 954.5498034 | 1098.655792 | -3.548999013 | Yes | 12S sens | 128 |
| SB_0067 | SB_0067 | MGEGLEENPTNPITKPTLNRNKAYIIILARTT TTTNDMKNHRCISTTRTPMTPMRKTNPLM KLINHSFIDLPTPSNISAWWNFGSLLGACLIL QITTGLFLAMHYSPDASTAFSSIAHITRDVNY GWIIRYLHANGASMFFICLFLHIGRGLYYGSF LYSETWNIGIILLLATMATAFMGYVLPWGQ MSFWGATVITNLLSAIPYIGTDLVQWIWGG YSVDSPTLTRFFTFHFILPFIIAALATLHLLFLH ETGSNNPLGITSHSDKITFHPYYTIKDALGLLL FLLSLMTLTLFSPDLLGDPDNYTLANPLNTPP HIKPEWYFLFAYTILRSVPNKLGGVLALLLSILI LAMIPILHMSKQQSMMFRPLSQSLYWLLA ADLLILTWIGGQPVSYPFTIIGQVASVLYFTTI  LILMPTISLIENKMLKWACPCSMN | 63 | 7 | 21.38188719 | 0.002212389381 | KTNPLMK | 830.4683856 | 1262.772811 | 1.41407854 | Yes | RefProt CYTB | 436 |
| SB_0231 | SB_0231 | IANSKKQLQTCRGFSRLFSRRREK | 36 | 12 | 21.06855371 | 0.00218579235 | GFSRLFSRR | 1124.62028 | 1268.717786 | 3.615045629 | Yes | Non-codant (antisens entre nd2 et  cox1) | 24 |
| SB_0187 | SB_0187 | MGVGMEWGLLLRRGRRRWCWGCGLLVV | 20 | 1 | 20.65512067 | 0.001 | MGVGME WGLLLR | 1360.699513 | 1520.792566 | 2.603800845 | Yes | COX1,  antisens - frameshift de Gau | 27 |
| SB_0050 | SB_0050 | IPTTQLKLQHHDPTTISHLKQANMTNTLNSI HPPLPRRPAPANRLFAQMGHYRRIHKKQ | 41 | 1 | 20.32219288 | 0.000999000999 | QANMTNT  LNSIHPPLP R | 1902.973367 | 2047.067215 | 4.036097747 | Yes | ND2 sens, frameShift | 59 |
| SB_0193 | SB_0193 | MLRFCVAGFGLIHLNCLLWWMRLREWGE  GLRLVRERFGMWLRWGLVFVMWEEAGR MSEGCLG | 65 | 1 | 19.80232168 | 0.000999000999 | LVRERFGM WLR | 1461.802668 | 1605.90347 | 0.8099306964 | Yes | ND2, antisens | 62 |
| SB_0119 | SB_0119 | MMINKRDDMTISGRLVVCRAHGRGKRRAI  SRSNNKKVMATKKNFMEKGTRAGDMGSK PHS | 64 | 4 | 19.56720182 | 0.002424242424 | DDMTISGR | 893.3912624 | 1053.484827 | 3.257502508 | Yes | ND3, antisens | 60 |
| SB_0015 | SB_0015 | MVLNLRVHRLRRTNLQLLHTSPIIPRTRRPA TPWRWQSSSTPDWSPHSYNNYITRRLALM SCPHIRLKNRCNSRTSKPNHFHRYTTGGML RSMLWNLWSKPQFHAHRPRINSPKNLWN  RARIYPMAPPLPPLEPTVKLT | 49 | 2 | 19.26859725 | 0.008849557522 | NLWNRAR | 928.4991084 | 1072.603732 | -2.365401482 | Yes | COX2 sens, frameShift | 139 |
| SB_0022 | SB_0022 | **MRHNYNKLHLPTTNRPKIAHCMLFNQPH SPRSNSHSHPNPLKLHRRSHSHNRPRAYILI TILPSKLKLRTHSQSHHNPLSRTSNSTPTNS**  **FLMTSSKPR** | 62 | 11 | 19.17011656 | 0.002018163471 | LHLPTTNRP K | 1175.677457 | 1463.877578 | 2.758316996 | Yes | ND4 sens, frameShift | 99 |
| SB_0138 | SB_0138 | MGCDRWHGEFWILRDGFDSHSPRNKGV | 20 | 5 | 18.53962327 | 0.003144654088 | DGFDSHSP R | 1016.431152 | 1160.534518 | -1.106494702 | Yes | Non-codant  (sens entre ND1 et ND2) | 27 |
| SB_0001 | SB_0001 | IWYFRLGGMHAMALRDAGAGAPYVAVSV  FDSCLILLFIAPTFNITGEHTY | 37 | 4 | 18.28136369 | 0.001020408163 | LGGMHAM  ALR | 1055.536813 | 1215.632296 | 1.251744129 | Yes | D-loop, sens | 50 |
| SB_0225 | SB_0225 | MPVSGGGFEAKVMFGCKVKY | 35 | 1 | 17.54348204 | 0.000999000999 | MPVSGGGF  EAK | 1078.511705 | 1382.713954 | -2.302300023 | Yes | COX3,  antisens | 20 |

| SB_0053 | SB_0053 | MKITSELVKRGLTPVFRFTVQCFTQPFYLTPT  DVRRPLTILYKPQRHWNTMPIIRRMSWSPR HSSKPPYSSRAGPARQPSR | 31 | 2 | 17.19087616 | 0.001001001001 | HWNTMPII RR | 1322.702959 | 1466.810724 | -3.863706822 | Yes | COX1 sens, frameShift | 81 |
| --- | --- | --- | --- | --- | --- | --- | --- | --- | --- | --- | --- | --- | --- |
| SB_0202 | SB_0202 | MGWARGGEVDRGLSITEQAPLEGYEAPPG  PLSFKLWLVVFWRAVLLI | 13 | 1 | 17.17685495 | 0.000999000999 | MGWARG  GEVDR | 1232.572012 | 1376.671919 | 1.585656856 | Yes | 12S, antisens | 47 |
| SB_0068 | SB_0068 | ILCSFMGKQIWVPPKYWLTHQQPLCISYITA  SHHEYCTVP | 13 | 4 | 16.55336317 | 0.001092896175 | ILCSFMGK | 897.4452082 | 1242.673333 | -2.022001366 | Yes | D-loop, sens | 40 |
| SB_0163 | SB_0163 | MGDCAVCDARVESEYVGEMKCA | 15 | 1 | 16.17840283 | 0.005076142132 | VESEYVGE  MK | 1169.527414 | 1617.826131 | 1.490421419 | Yes | ND5, antisens | 22 |
| SB_0238 | SB_0238 | INKLKLHRVFSSCCVMPASSRAGQFHWLKV RDSWTLVEPFMQVPI | 47 | 3 | 15.62436483 | 0.00218579235 | AGQFHWL K | 985.5133602 | 1273.722598 | -3.994077948 | Yes | 16S antisens  (chevauche humanin) | 45 |
| SB_0235 | SB_0235 | MNRRPRLRLTRGLGMGRGVHSRRAMVRA  KVGAVM | 31 | 2 | 14.75986827 | 0.005020080321 | GVHSRRA  MVR | 1167.640697 | 1311.744847 | -1.567282974 | Yes | ND1, antisens | 34 |
| SB_0046 | SB_0046 | MAEPGNRMKLKTLQSEVQFLFLTTYPWPTS YSSLYPF | 18 | 3 | 13.93237678 | 0.008869179601 | MAEPGNR | 773.3490056 | 917.4533302 | -2.447302268 | Yes | Non-codant (antisens entre 16S et  ND1) | 37 |
| SB_0032 | SB_0032 | MHQSCKPEMKTFFQGQIREKVFNSTISTQS | 20 | 2 | 13.83337106 | 0.003003003003 | MHQSCKPE MK | 1217.535493 | 1722.853313 | 2.770027249 | Yes | Non-codant (sens entre CYTB et d-  loop) | 30 |
| SB_0029 | SB_0029 | IHRPPHPIQHLRMMKLRLTPWRLPDPPNH HRTIPSHALLTRRLNRLFINRPHHSRRKLWL NHPLPSRQWRLNILYLPLPTHRARPMLRIISL LRNLKHRHYPPACNYSNSLHRLCPPVRPNIIL RGHSNYKLTIRHPMHWDRPSSMNLRRLLS RQSHPHTILYLSLHLALHYCSPSNTPPPILAR  NGIKQPPRNHLPFR | 121 | 2 | 13.82029738 | 0.004424778761 | LRLTPWR | 940.5606422 | 1084.660857 | 1.728571636 | Yes | RefProt CYTB | 199 |
| SB_0172 | SB_0172 | MIVRGRSQVVSIRRGVVRGSEEKVGEQLNR LLLIWLKNSRGMMLMIRLWVVVLIQIMCFL ESHVSGSNMIVGTISFSIGVGLGYVRSLGHM  CWRLRLVGLGPPLLRRRQRLVWQ | 155 | 1 | 12.06194623 | 0.009433962264 | GRSQVVSIR | 1000.577749 | 1144.676438 | 2.971509154 | Yes | ND4, antisens | 114 |
| SB_0049 | SB_0049 | IFYLSRPRNKHASFYSSSNQKNKPSFHRSCH QVFPHASNRIHNPSNSYPLQQYTLRTMNH NQYYQSMLIINNHNSYSNKTRNSPLSLLSPR GYPRHPSDIRPASSHMTKTSPHLNHMPNLS LTKRKPSPHSLNLIHHSRQLRWIKPNPATQN  LSMLLNYPHRMNNSSSTVQP | 122 | 4 | 12.05265959 | 0.007658643326 | IFYLSRPR | 1050.59742 | 1338.79671 | 3.632896529 | Yes |  | 172 |
| SB_0103 | SB_0103 | MLTSKYSNQPSTITHQLQLQSHPSPTRMPT  NLPTLNST | 125 | 3 | 11.44115982 | 0.002997002997 | MPTNLPTL  NST | 1187.585595 | 1347.676777 | 4.321420165 | Yes | D-loop, sens | 38 |

**ANTIBODY PRODUCTION (antigen in red)**

**MGLSRIEGLFGQVVCGGLGMCFLVLHRAIIGMWLVCWLVGLVWGALWSGSEITWLGRRSLGGLRGPLLGVMGWVLLYDRHVIGGSLCVVVQVEAY**

**MWSLPRRLPGWPSSARMRRLRAVPRTPAHAPNNRYSVPMSLWFVENSQRSANISGGEVKWLSEALDCKSKDRG**

**IMRMTAPVKLQGVWMRMAVTTRAMWLIEEYAMSDFRSVCRRQMELVMIMPHRDSTRKG**

**MRHNYNKLHLPTTNRPKIAHCMLFNQPHSPRSNSHSHPNPLKLHRRSHSHNRPRAYILITILPSKLKLRTHSQSHHNPLSRTSNSTPTNSFLMTSSKPR**

**Table S5. List of proteins found by mass spectrometry following immunoprecipitation or pull down assay**

| **Immunoprecipitation** | | | | |
| --- | --- | --- | --- | --- |
| **PROTEIN NAME** | **ACCESSION NUMBER** | **MOLECULA R WEIGHT** | **UNIQUE PEPTIDES COUNT** | **PROBABILITY*** |
| Protein [synthetic construct, MTALTND4] | AEX63613.1 | 16 kDa | 5 | 100% |
| Glutathione peroxidase 1 | P07203 | 22 kDa | 13 | 100% |
| Echinoderm microtubule-associated protein-like 4 OS=Homo sapiens GN=EML4 PE=1 SV=3 | Q9HC35 | 109 kDa | 12 | 100% |
| Arf-GAP with coiled-coil, ANK repeat and PH domain-containing protein 2 OS=Homo sapiens GN=ACAP2 PE=1 SV=3 | Q15057 | 88 kDa | 2 | 100% |
| Cleavage stimulation factor subunit 2 tau variant OS=Homo sapiens GN=CSTF2T PE=1 SV=1 | Q9H0L4 | 64 kDa | 2 | 100% |
| Glutaredoxin domain-containing cysteine-rich protein 2 OS=Homo sapiens GN=GRXCR2 PE=3 SV=1 | A6NFK2 | 28 kDa | 2 | 100% |
| **Complement component 1 Q subcomponent-binding protein, mitochondrial OS=Homo sapiens GN=C1QBP PE=1 SV=1** | **Q07021** | **31 kDa** | **2** | **100%** |
| **Complement C1q subcomponent subunit C OS=Homo sapiens GN=C1QC PE=1 SV=3** | **P02747** | **26 kDa** | **2** | **100%** |
| **Complement C3 alpha chain (Fragment) OS=Oryctolagus cuniculus OX=9986 GN=C3 PE=2 SV=1** | **P12247** | **82 kDa** | **2** | **100%** |
| 60S ribosomal protein L26 | P61254 | 17 kDa | 3 | 100% |
| Histone H4 OS=Homo sapiens | P62805 | 11 kDa | 3 | 100% |
| Glutamine amidotransferase-like class 1 domain- containing protein 1 | Q8NB37 | 23 kDa | 2 | 100% |
| 40S ribosomal protein S9 | P46781 | 23 kDa | 2 | 100% |
| 40S ribosomal protein S3a | P61247 | 30 kDa | 2 | 100% |
| Putative 60S ribosomal protein L13a protein RPL13AP3 | Q6NVV1 | 12 kDa | 2 | 100% |
| 40S ribosomal protein S11 | P62280 | 18 kDa | 2 | 100% |

| **Pull down #1** | | | | |
| --- | --- | --- | --- | --- |
| **PROTEIN NAME** | **ACCESSION NUMBER** | **MOLECULA R WEIGHT** | **UNIQUE PEPTIDES COUNT** | **PROBABILITY** |
| Nucleophosmin OS=Homo sapiens OX=9606 GN=NPM1 PE=1 SV=2 | P06748 | 33 kDa | 14 | 100% |
| Elongation factor 1-gamma OS=Homo sapiens OX=9606 GN=EEF1G PE=1 SV=3 | P26641 | 50 kDa | 13 | 100% |
| **Complement component 1 Q subcomponent-binding protein, mitochondrial OS=Homo sapiens OX=9606 GN=C1QBP PE=1 SV=1** | **Q07021** | **31 kDa** | **12** | **100%** |
| Nucleosome assembly protein 1-like 1 OS=Homo sapiens OX=9606 GN=NAP1L1 PE=1 SV=1 | P55209 | 45 kDa | 10 | 100% |
| Protein phosphatase 1G OS=Homo sapiens OX=9606 GN=PPM1G PE=1 SV= | O15355 | 59 kDa | 9 | 100% |
| Protein SET OS=Homo sapiens OX=9606 GN=SET PE=1 SV= | Q01105 | 33 kDa | 9 | 100% |
| Carbonyl reductase [NADPH] 1 OS=Homo sapiens OX=9606 GN=CBR1 PE=1 SV=3 | P16152 | 30 kDa | 8 | 100% |
| Nucleosome assembly protein 1-like 4 OS=Homo sapiens OX=9606 GN=NAP1L4 PE=1 SV= | Q99733 | 43 kDa | 8 | 100% |
| 40S ribosomal protein S18 OS=Homo sapiens OX=9606 GN=RPS18 PE=1 SV=3 | P62269 | 18 kDa | 7 | 100% |
| Proliferating cell nuclear antigen OS=Homo sapiens OX=9606 GN=PCNA PE=1 SV=1 | P12004 | 29 kDa | 7 | 100% |
| Cluster of Tubulin alpha-1B chain OS=Homo sapiens OX=9606 GN=TUBA1B PE=1 SV=1 (P68363) | P68363 | 50 kDa | 6 | 100% |
| 60 kDa heat shock protein, mitochondrial OS=Homo sapiens OX=9606 GN=HSPD1 PE=1 SV=2 | P10809 | 61 kDa | 6 | 100% |
| Acidic leucine-rich nuclear phosphoprotein 32 family member B OS=Homo sapiens OX=9606 GN=ANP32B PE=1 SV=1 | Q92688 | 29 kDa | 6 | 100% |
| Cluster of Elongation factor 1-alpha 1 OS=Homo sapiens OX=9606 GN=EEF1A1 PE=1 SV=1 (P68104) | P68104 | 50 kDa | 5 | 100% |
| 40S ribosomal protein S3 OS=Homo sapiens OX=9606 GN=RPS3 PE=1 SV=2 | P23396 | 27 kDa | 5 | 100% |
| 40S ribosomal protein S13 OS=Homo sapiens OX=9606 GN=RPS13 PE=1 SV=2 | P62277 | 17 kDa | 5 | 100% |
| Cluster of Heterogeneous nuclear ribonucleoprotein H OS=Homo sapiens OX=9606 GN=HNRNPH1 PE=1 SV=4 (P31943) | P31943 | 49 kDa | 5 | 100% |
| ATP synthase subunit beta, mitochondrial OS=Homo sapiens OX=9606 GN=ATP5F1B PE=1 SV=3 | P06576 | 57 kDa | 5 | 100% |
| Endoplasmic reticulum chaperone BiP OS=Homo sapiens OX=9606 GN=HSPA5 PE=1 SV=2 | P11021 | 72 kDa | 4 | 100% |
| Elongation factor Tu, mitochondrial OS=Homo sapiens OX=9606 GN=TUFM PE=1 SV=2 | P49411 | 50 kDa | 4 | 100% |
| Elongation factor 1-beta OS=Homo sapiens OX=9606 GN=EEF1B2 PE=1 SV=3 | P24534 | 25 kDa | 4 | 100% |
| Casein kinase II subunit alpha OS=Homo sapiens OX=9606 GN=CSNK2A1 PE=1 SV=1 | P68400 | 45 kDa | 4 | 100% |
| GTP-binding nuclear protein Ran OS=Homo sapiens OX=9606 GN=RAN PE=1 SV=3 | P62826 | 24 kDa | 4 | 100% |
| Prostaglandin E synthase 2 OS=Homo sapiens OX=9606 GN=PTGES2 PE=1 SV=1 | Q9H7Z7 | 42 kDa | 4 | 100% |
| ATP synthase subunit alpha, mitochondrial OS=Homo sapiens OX=9606 GN=ATP5F1A PE=1 SV=1 | P25705 | 60 kDa | 3 | 100% |
| Peroxiredoxin-1 OS=Homo sapiens OX=9606 GN=PRDX1 PE=1 SV=1 | Q06830 | 22 kDa | 3 | 100% |
| Glyceraldehyde-3-phosphate dehydrogenase OS=Homo sapiens OX=9606 GN=GAPDH PE=1 SV=3 | P04406 | 36 kDa | 3 | 100% |
| 40S ribosomal protein S14 OS=Homo sapiens OX=9606 GN=RPS14 PE=1 SV=3 | P62263 | 16 kDa | 3 | 100% |
| 40S ribosomal protein S20 OS=Homo sapiens OX=9606 GN=RPS20 PE=1 SV=1 | P60866 | 13 kDa | 3 | 100% |
| 40S ribosomal protein S10 OS=Homo sapiens OX=9606 GN=RPS10 PE=1 SV= | P46783 | 19 kDa | 3 | 100% |
| 60S acidic ribosomal protein P0 OS=Homo sapiens OX=9606 GN=RPLP0 PE=1 SV=1 | P05388 | 34 kDa | 3 | 100% |
| 60S ribosomal protein L5 OS=Homo sapiens OX=9606 GN=RPL5 PE=1 SV=3 | P46777 | 34 kDa | 3 | 100% |
| 60S ribosomal protein L30 OS=Homo sapiens OX=9606 GN=RPL30 PE=1 SV=2 | RPL30 | 13 kDa | 3 | 100% |
| Histone H2A type 1-B/E OS=Homo sapiens OX=9606 GN=H2AC4 PE=1 SV=2 | P04908 | 14 kDa | 3 | 100% |
| Y-box-binding protein 1 OS=Homo sapiens OX=9606 GN=YBX1 PE=1 SV=3 | P67809 | 36 kDa | 3 | 100% |
| 40S ribosomal protein S16 OS=Homo sapiens OX=9606 GN=RPS16 PE=1 SV=2 | P62249 | 16 kDa | 2 | 100% |
| 40S ribosomal protein S25 OS=Homo sapiens OX=9606 GN=RPS25 PE=1 SV=1 | P62851 | 14 kDa | 2 | 100% |
| 40S ribosomal protein S5 OS=Homo sapiens OX=9606 GN=RPS5 PE=1 SV=4 | P46782 | 23 kDa | 2 | 100% |
| 60S ribosomal protein L22 OS=Homo sapiens OX=9606 GN=RPL22 PE=1 SV=2 | P35268 | 15 kDa | 2 | 100% |
| 60S ribosomal protein L27a OS=Homo sapiens OX=9606 GN=RPL27A PE=1 SV=2 | P46776 | 17 kDa | 2 | 100% |
| ATP synthase subunit O, mitochondrial OS=Homo sapiens OX=9606 GN=ATP5PO PE=1 SV=1 | P48047 | 23 kDa | 2 | 100% |
| Eukaryotic initiation factor 4A-I OS=Homo sapiens OX=9606 GN=EIF4A1 PE=1 SV=1 | P60842 | 46 kDa | 2 | 100% |
| Protein arginine N-methyltransferase 5 OS=Homo sapiens OX=9606 GN=PRMT5 PE=1 SV=4 | O14744 | 73 kDa | 2 | 100% |
| GrpE protein homolog 1, mitochondrial OS=Homo sapiens OX=9606 GN=GRPEL1 PE=1 SV=2 | Q9HAV7 | 24 kDa | 2 | 100% |

*Only results with 100% probability and two or more unique peptides count are presented.

| **Pull down #2 (only C1Q is presented)** | | | | | |
| --- | --- | --- | --- | --- | --- |
| **PROTEIN NAME** | **ACCESSION NUMBER** | **MOLECULA R WEIGHT** | **CELL TYPE** | **UNIQUE PEPTIDES COUNT** | **PROBABILITY** |
| **Complement component 1 Q subcomponent-binding protein, mitochondrial OS=Homo sapiens OX=9606 GN=C1QBP PE=1 SV=1** | **Q07021** | **31 kDa** | **HEK-293T** | **37** | **100%** |
| **Complement component 1 Q subcomponent-binding protein, mitochondrial OS=Homo sapiens OX=9606 GN=C1QBP PE=1 SV=1** | **Q07021** | **31 kDa** | **HeLa** | **16** | **100%** |
| **VALIDATION OF THE INTERACTION BY WESTERN BLOT**  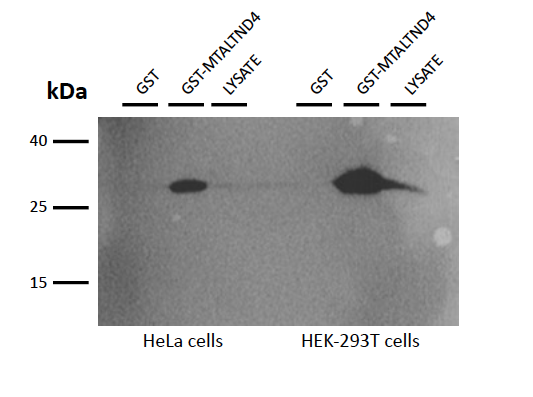  **GST pull-down assay indicating that MTALTND4 and Complement component 1 Q interact.** Western blot probed with anti-GC1q antibodies. Lane 1: pull-down products from GST bound glutathione beads (HeLa cells). Lane 2: pull-down products from GST-MTALTND4–bound glutathione beads (HeLa cells). Lane 3: HeLa cells lysate. Lane 4: pull-down products from GST bound glutathione beads (HEK-293T cells). Lane 2: pull-down products from GST-MTALTND4–bound glutathione beads (HEK-293T cells). Lane 3: HEK-293T cells lysate. | | | | | |

**Table S6. Dose-dependent impact of MTALTND4 on routine respiration of intact HeLa and HEK-293T cells.** Respirometry data were normalized for their own routine respiration (i.e. flux control ratios - FCR) in absence of the peptide. For each titration point, FCR have been expressed as fraction of their own simultaneous paired control. MTALTND4 was sequentially titrated to final concentrations of: 0.1 µM, 1 µM, 5 µM, 10 µM, 15µM, 20 µM, 25 µM, 30 µM, 35 µM, 40 µM, 45 µM. HeLa (*n*=4, up to 30 µM), HEK-293T (*n*=3, up to 45 µM). Controls were titrated with an equal volume of H_2_O. Factors: *'cell type'* (*'hek'* = HEK-293T, *'hela'* = HeLa); *'ID'* (paired sample identificator); *'treatment'* ('*ctrl'* = control, *'pept'* = peptide treatment). For each parameter referring to a specific titration point, the impact of peptide addition (*':treatment', i.e.* control vs treatment) was inferred by means of a two-sided one sample *t* test. *p*-values were adjusted using Holm's correction for multiple testing. ·0.05 > *p* ≤ 0.09; **p* ≤ 0.05; ***p* ≤ 0.01; ****p* ≤ 0.001. Data are reported separately and also as mean ± standard error of the mean (sem). Data refer to figure 3*A,B*.

| Table S6 | | | **MTALTND4 concentration** | | | | | | | | | | |
| --- | --- | --- | --- | --- | --- | --- | --- | --- | --- | --- | --- | --- | --- |
| **cell type** | **ID** | **treatment** | **0.1 µM** | **1 µM** | **5 µM** | **10 µM** | **15 µM** | **20 µM** | **25 µM** | **30 µM** | **35 µM** | **40 µM** | **45 µM** |
| *hek*  *hek* | *1hk*  *1hk* | *ctrl*  *pept* | 1  0.979 | 1  0.961 | 1  0.971 | 1  0.939 | 1  0.902 | 1  0.892 | 1  0.754 | 1  0.663 | 1  0.630 | 1  0.581 | 1  0.511 |
| *hek*  *hek* | *2hk*  *2hk* | *pept*  *ctrl* | 0.989  1 | 1.036  1 | 0.927  1 | 0.937  1 | 0.901  1 | 0.740  1 | 0.739  1 | 0.720  1 | 0.597  1 | 0.599  1 | 0.562  1 |
| *hek*  *hek* | *3hk*  *3hk* | *pept*  *ctrl* | 0.994  1 | 0.959  1 | 1.001  1 | 0.990  1 | 0.948  1 | 0.697  1 | 0.663  1 | 0.689  1 | 0.652  1 | 0.548  1 | 0.559  1 |
| *hela*  *hela* | *1hl*  *1hl* | *ctrl*  *pept* | 1  0.998 | 1  0.970 | 1  0.742 | 1  0.500 | 1  0.380 | 1  0.350 | 1  0.301 | 1  0.148 | *na* | *na* | *na* |
| *hela*  *hela* | *0hl*  *0hl* | *ctrl*  *pept* | 1  0.946 | 1  0.899 | 1  0.604 | 1  0.384 | 1  0.270 | 1  0.171 | 1  0.101 | 1  0.097 |  |  |  |
| *hela*  *hela* | *2hl*  *2hl* | *ctrl*  *pept* | 1  1.004 | 1  0.899 | 1  0.677 | 1  0.551 | 1  0.475 | 1  0.431 | 1  0.299 | 1  0.215 |  |  |  |
| *hela*  *hela* | *3hl*  *3hl* | *ctrl*  *pept* | 1  1.039 | 1  1.016 | 1  0.848 | 1  0.661 | 1  0.560 | 1  0.506 | 1  0.466 | 1  0.271 |  |  |  |
| *HEK-293T* | *Stats* | *ctrl (mean ± sem)*  *pept (mean ± sem)* | 1 ± 0  0.987 ± 0.004 | 1 ± 0  0.985 ± 0.025 | 1 ± 0  0.967 ± 0.021 | 1 ± 0  0.955 ± 0.017 | 1 ± 0  0.917 ± 0.016 | 1 ± 0  0.776 ± 0.059 | 1 ± 0  0.719 ± 0.028 | 1 ± 0  0.691 ± 0.016 | 1 ± 0  0.626 ± 0.016 | 1 ± 0  0.576 ± 0.015 | 1 ± 0  0.544 ± 0.017 |
|  |  | *transformation* | *none* | *none* | *none* | *none* | *none* | *none* | *none* | *none* | *none* | *none* | *none* |
|  |  | *:treatment* | *t*=-2.95, *P*=0.098 | *t*=-0.58, *P*=0.62 | *t*=-1.56, *P*=0.259 | *t*=-2.59, *P*=0.122 | ***t*=-5.31, *P*=0.0336*** | *t*=-3.79, *P*=0.063∙ | ***t*=-9.99, *P*=9.856e-03**** | ***t*=-18.84, *P*=2.8e-03**** | ***t*=-23.33, *P*=1.8e-03**** | ***t*=-28.75, *P*=1.2e-03**** | ***t*=-27.56, *P*=1.3e-03**** |
|  |  | *p adjusted* | 0.39272 | 0.6205 | 0.5184 | 0.39272 | 0.20196 | 0.3154 | 0.068992∙ | **0.022448*** | **0.016479*** | **0.013277*** | **0.013277*** |
| *HeLa* | *Stats* | *ctrl (mean ± sem)*  *pept (mean ± sem)* | 1 ± 0  0.997 ± 0.019 | 1 ± 0  0.946 ± 0.029 | 1 ± 0  0.718 ± 0.052 | 1 ± 0  0.524 ± 0.058 | 1 ± 0  0.421 ± 0.062 | 1 ± 0  0.364 ± 0.072 | 1 ± 0  0.292 ± 0.075 | 1 ± 0  0.183 ± 0.038 | *na* | *na* | *na* |
|  |  | *transformation* | *none* | *none* | *none* | *none* | *none* | *none* | *none* | *none* |  |  |  |
|  |  | *:treatment* | *t*=-0.16, *P*= 0.879 | *t*=-1.88, *P*=0.156 | ***t*=-5.45, *P*=1.2e-02*** | ***t*=-8.27, *P*=3.6e-03**** | ***t*=-9.27, *P*=2.6e-03**** | ***t*=-8.84, *P*=3e-03**** | ***t*=-9.5, *P*=2.4e-03**** | ***t*=-21.5, *P*=2.2e-04***** |  |  |  |
|  |  | *p adjusted* | 0.8798 | 0.3134 | **0.03642*** | **0.017269*** | **0.017269*** | **0.017269*** | **0.017269*** | **0.0017608**** |  |  |  |

**Table S7. Impact of 10 µM MTALTND4 on mitochondrial respiration of intact HeLa and HEK-293T cells.** Respirometry data were normalized for their own routine respiration in absence of the peptide (*'ce_R'*) and expressed as flux control ratios (FCR). HeLa (*n*=6-9, depending the parameter), HEK-293T (*n*=6). Factors: *'cell type'* (*'hek'* = HEK-293T, *'hela'* = HeLa); *'ID'* (paired sample identificator); *'treatment'* (*'ctrl'* = control, *'pept'* = peptide treatment). *'ce_R'*: intact cells routine respiration; *'ce-pept_R'*: intact cells routine respiration with MTALTND4; *'ce-pept_L'*: intact cell leak respiration with MTALTND4; *'ce-pept_E'*: intact cell maximal uncoupled respiration with MTALTND4; *'SRC'*: mitochondrial spare reserve capacity (*'ce-pept_E'* − *'ce_R'*) of intact cells with MTALTND4; Controls were titrated with an equal volume of H_2_O. The impact of peptide addition (*':treatment'*, i.e. control vs treatment) was inferred by means of a two samples paired *t* test. *p*-values were adjusted using Holm's correction for multiple testing. For SRC, a linear mixed model (LMM) was implemented. Fixed effects: *"cell type"* and *"treatment"*. Random effect: *"ID"*. *':treatment'* = main effect of treatment; *':cell type'* = main effect of cell type; *':interaction'* = interaction effect. **·**0.05 > *p* ≤ 0.09; ******p* ≤ 0.05; *******p* ≤ 0.01; ********p* ≤ 0.001. Data are reported separately and also as mean ± standard error of the mean (sem). Data refer to figure 3*C,D,E*.

| **cell type** | **ID** | **treatment** | **ce_R** | **ce-pept_R** | **ce-pept_L** | **ce-pept_E** |  | **SRC (E-R)** | |
| --- | --- | --- | --- | --- | --- | --- | --- | --- | --- |
| *hek*  *hek* | *1hk*  *1hk* | *ctrl*  *pept* | 1  1 | 0.908  0.962 | 0.249  0.175 | 3.687  2.988 |  |  | 2.687  1.988 |
| *hek*  *hek* | *2hk*  *2hk* | *ctrl*  *pept* | 1  1 | 0.972  0.950 | 0.219  0.210 | 3.007  2.258 |  |  | 2.007  1.258 |
| *hek*  *hek* | *3hk*  *3hk* | *ctrl*  *pept* | 1  1 | 0.971  0.894 | 0.219  0.195 | 3.826  1.924 |  |  | 2.826  0.924 |
| *hek*  *hek* | *4hk*  *4hk* | *ctrl*  *pept* | 1  1 | 0.941  0.808 | 0.241  0.220 | 3.658  2.255 |  |  | 2.658  1.255 |
| *hek*  *hek* | *5hk*  *5hk* | *ctrl*  *pept* | 1  1 | 0.987  0.825 | 0.295  0.230 | 3.520  2.224 |  |  | 2.520  1.224 |
| *hek*  *hek* | *6hk*  *6hk* | *pept*  *ctrl* | 1  1 | 0.886  0.956 | 0.184  0.219 | 3.264  4.109 |  |  | 2.264  3.109 |
| *hela*  *hela* | *1hl*  *1hl* | *ctrl*  *pept* | 1  1 | 0.976  0.464 | 0.241  0.138 | 2.705  0.592 |  |  | 1.705  -0.408 |
| *hela*  *hela* | *2hl*  *2hl* | *ctrl*  *pept* | 1  1 | 0.867  0.786 | 0.140  0.112 | 2.130  1.967 |  |  | 1.130  0.967 |
| *hela*  *hela* | *3hl*  *3hl* | *pept*  *ctrl* | 1  1 | 0.283  0.566 | 0.066  0.062 | 0.339  1.647 |  |  | -0.661  0.647 |
| *hela*  *hela* | *4hl*  *4hl* | *ctrl*  *pept* | 1  1 | 0.951  0.692 | 0.126  0.102 | 2.480  1.582 |  |  | 1.480  0.582 |
| *hela*  *hela* | *5hl*  *5hl* | *ctrl*  *pept* | 1  1 | 0.730  0.635 | 0.114  0.099 | 1.854  1.254 |  |  | 0.854  0.254 |
| *hela*  *hela* | *6hl*  *6hl* | *pept*  *ctrl* | 1  1 | 0.557  0.803 | 0.125  0.221 | 0.877  1.490 |  |  | -0.123  0.490 |
| *hela*  *hela* | *x1hl*  *x1hl* | *ctrl*  *pept* | 1  1 | 0.878  0.460 | *na* | *na* |  |  | *na* |
| *hela*  *hela* | *x2hl*  *x2hl* | *ctrl*  *pept* | 1  1 | 0.922  0.459 |  |  |  |  |  |
| *hela*  *hela* | *x3hl*  *x3hl* | *ctrl*  *pept* | 1  1 | 0.946  0.389 |  |  |  |  |  |
|  |  | *ctrl (mean ± sem)*  *pept (mean ± sem)* | 1 ± 0  1 ± 0 | 0.956 ± 0.012  0.888 ± 0.026 | 0.241 ± 0.012  0.202 ± 0.009 | 3.634 ± 0.15  2.486 ± 0.212 |  | *HEK293* | 2.634 ± 0.15  1.486 ± 0.212 |
| *HEK-293T* | *Stats* | *transformation* | *none* | *none* | *none* | *none* |  | *HELA* | 1.051 ± 0.195  0.102 ± 0.251 |
|  |  | *:treatment* |  | *t*=2.15, *P*=0.084∙ | ***t*=3.56, *P*=0.016*** | ***t*=5.97, *P*=0.0018**** |  |  |  |
|  |  | *p adjusted* |  | 0.08412 | **0.03232*** | **0.005628**** |  | LMM | |
| *HELA* | *Stats* | *ctrl (mean ± sem)*  *pept (mean ± sem)* | 1 ± 0  1 ± 0 | 0.849 ± 0.044  0.525 ± 0.053 | 0.151 ± 0.028  0.107 ± 0.01 | 2.051 ± 0.195  1.102 ± 0.251 |  |  |  |
|  |  |  |  |  |  |  |  | *transformation* | *none* |
|  |  | *transformation* | *none* | *none* | *none* | *none* |  | *:cell type* | ***F*1,12=47.68, *P*=1.639e-05 ***** |
|  |  | *:treatment* |  | ***t*=5.58, *P*=5.1e-04***** | *t*=2.38, *P*=0.062∙ | ***t*=3.4, *P*=0.019*** |  | *:treatment* | ***F*1,12=45.99, *P*=1.952e-05 ***** |
|  |  | *p adjusted* |  | **0.0015507**** | 0.06281∙ | **0.03848*** |  | *:interaction* | *F*1,12=0.4165, *P*=0.5308 |

**Table S8.** **Impact of 10 µM MTALTND4 on mitochondrial respiration of permeabilized HEK-293T cells.** Respirometry data were expressed as flux control ratios (FCR), normalized for the maximal coupled respiration (state 3) sustained by CI+CII-linked substrates in absence of MTALTND4 (*'CI+II_P'*). HEK-293T (*n*=6). Factors: *'cell type'* (*'hek'* = HEK-293T); *'ID'* (paired sample identificator); *'treatment'* (*'ctrl'* = control, *'pept'* = peptide treatment). *'ce_R'*: routine respiration of intact cells; *'CI_L'*: CI-sustained leak respiration; *'CI_P'*: CI-linked coupled respiration (state 3); *'c_P'*: CI-linked coupled respiration in presence of cytochrome *c*; *'CI+II_P'*: CI+II-linked coupled respiration (state 3); *'CI+II-pept_P'*: CI+II-linked coupled respiration with MTALTND4 (state 3 + peptide); *'CI+II-pept_E'*: CI+II-linked uncoupled respiration with MTALTND4 (state 3u + peptide); *'CIV-pept_E'*: cytochrome *c* oxidase standalone capacity with MTALTND4; Controls were titrated with an equal volume of H_2_O. The impact of peptide addition (*':treatment'*) was inferred by means of a two samples paired *t* test. *p*-values were adjusted using Holm's correction for multiple testing. **·**0.05 > *p* ≤ 0.09; ******p* ≤ 0.05; *******p* ≤ 0.01; ********p* ≤ 0.001. Data are reported separately and also as mean ± standard error of the mean (sem). Data refer to figure 3*F*.

| **cell type** | **ID** | **treatment** | **ce_R** | **CI_L** | **CI_P** | **c_P** | **CI+II_P** | **CI+II-pept_P** | **CI+II-pept_E** | **Rot** | **CIV-pept_E** |
| --- | --- | --- | --- | --- | --- | --- | --- | --- | --- | --- | --- |
| *hek*  *hek* | *1hk*  *1hk* | *pept*  *ctrl* | 0.386  1.470 | 0.167  0.521 | 0.595  1.374 | 0.637  1.411 | 1  1 | 0.303  0.822 | 0.399  1.995 | 0.007  0.023 | 0.643  9.993 |
| *hek*  *hek* | *2hk*  *2hk* | *pept*  *ctrl* | 0.505  0.806 | 0.174  0.302 | 0.678  0.891 | 0.696  0.932 | 1  1 | 0.072  0.773 | 0.046  1.017 | 0.027  0.026 | 1.016  4.656 |
| *hek*  *hek* | *3hk*  *3hk* | *pept*  *ctrl* | 0.682  1.034 | 0.267  0.364 | 0.643  0.889 | 0.737  0.930 | 1  1 | 0.538  0.855 | 0.661  1.259 | 0.001  0.042 | 2.014  7.802 |
| *hek*  *hek* | *4hk*  *4hk* | *ctrl*  *pept* | 0.536  0.495 | 0.234  0.180 | 0.636  0.716 | 0.649  0.706 | 1  1 | 0.978  0.640 | 1.923  0.992 | -0.003  0.004 | 2.174  1.576 |
| *hek*  *hek* | *5hk*  *5hk* | *ctrl*  *pept* | 0.612  0.926 | 0.324  0.435 | 0.575  0.699 | 0.597  0.730 | 1  1 | 0.940  0.320 | 1.453  0.449 | 0.006  0.013 | 2.452  1.969 |
| *hek*  *hek* | *6hk*  *6hk* | *pept*  *ctrl* | 0.418  0.387 | 0.124  0.094 | 0.811  0.796 | 0.762  0.827 | 1  1 | 0.536  1.021 | 0.700  1.149 | 0.001  0.026 | 0.883  1.219 |
| *HEK-293T* | *Stats* | *ctrl (mean ± sem)*  *pept (mean ± sem)* | 0.807 ± 0.161  0.569 ± 0.083 | 0.307 ± 0.058  0.225 ± 0.046 | 0.86 ± 0.116  0.69 ± 0.03 | 0.891 ± 0.119  0.711 ± 0.018 | 1 ± 0  1 ± 0 | 0.898 ± 0.039  0.402 ± 0.085 | 1.466 ± 0.167  0.541 ± 0.131 | 0.02 ± 0.007  0.009 ± 0.004 | 4.716 ± 1.428  1.35 ± 0.238 |
|  |  | *transformation* | *none* | *none* | *none* | *none* | *none* | *none* | *none* | *none* | *none* |
|  |  | *:treatment* | *t*=1.21, *P*=0.27 | *t*=1.25, *P*=0.26 | *t*=1.24, *P*=0.27 | *t*=1.36, *P*=0.23 |  | ***t*=8.01, *P*=4.8e-04***** | ***t*=5.69, *P*=0.0023**** | *t*=1.39, *P*=0.22 | *t*=2.25, *P*=0.073∙ |
|  |  | *p adjusted* | 1 | 1 | 1 | 1 |  | **0.003912**** | **0.016331*** | 1 | 0.44346 |

**Table S9.** **Impact of MTALTND4 on intact HEK-293T cell's lactic fermentation, antioxidant capacity, ATP content and hydrogen peroxide efflux rate.** HEK-293T cells (*n* = 5) were incubated with 30 µM MTALTND4 for 4 hours. Data are reported either as enzymatic activity (U∙mg proteins^−1^) for lactate dehydrogenase I catalase, H_2_O_2_ efflux rate (pmol H_2_O_2_∙mg proteins^−1^∙min^−1^) and ATP content (nmol ATP∙mg proteins^−1^). LDH: lactate dehydrogenase; CAT: catalase; H_2_O_2_ efflux: hydrogen peroxide efflux rate; ATP content: amount of ATP present in the cells. The main effect of the fixed factor *‘:treatment’* was assessed for each parameter through two sample paired *t*-test or Wilcoxon signed rank test. *p*-values were adjusted using H’lm's correction for multiple testing. **·**0.05 > *p* ≤ 0.09; ******p* ≤ 0.05; *******p* ≤ 0.01; ********p* ≤ 0.001. Data are reported separately and also as mean ± standard error of the mean (sem). Data refer to figure 3*G,H,I,L*.

| Table S9 | | | U∙mg−1 pm | | ol H2O2∙mg−1∙min−1 | nmol ATP∙mg−1 |
| --- | --- | --- | --- | --- | --- | --- |
| **Cell-type** | **ID** | **Treatment** | **LDH** | **CAT** | **H2O2 efflux pmol** | **ATP content** |
| *hek* | *1hk* | *ctrl* | 0.427 | 3.249 | 9.948 | 1.359 |
| *hek* | *2hk* | *ctrl* | 0.318 | 2.729 | 8.258 | 1.296 |
| *hek* | *3hk* | *ctrl* | 0.384 | 2.951 | 8.807 | 1.459 |
| *hek* | *4hk* | *ctrl* | 0.310 | 2.728 | 7.109 | 1.200 |
| *hek* | *5hk* | *ctrl* | 0.321 | 2.761 | 7.835 | 1.549 |
| *hek* | *1hk* | *pept-30uM* | 0.305 | 3.154 | 14.375 | 1.827 |
| *hek* | *2hk* | *pept-30uM* | 0.289 | 1.876 | 12.150 | 1.845 |
| *hek* | *3hk* | *pept-30uM* | 0.346 | 3.023 | 15.021 | 1.940 |
| *hek* | *4hk* | *pept-30uM* | 0.292 | 2.385 | 9.361 | 1.593 |
| *hek* | *5hk* | *pept-30uM* | 0.391 | 2.873 | 9.861 | 2.271 |
|  |  | *ctrl (mean ± sem)* | 0.352 ± 0.023 | 2.884 ± 0.1 | 8.392 ± 0.478 | 1.373 ± 0.061 |
|  |  | *pept (mean ± sem)* | 0.325 ± 0.02 | 2.662 ± 0.236 | 12.154 ± 1.145 | 1.895 ± 0.11 |
| *HEK-293T* | *Stats* | *transformation* | *none* | *none* | *none* | *none* |
|  |  | *:treatment* | *v*=11, *P*=0.437 | *t*=1.25, *P*=0.279 | ***t*=-4.9, *P*=0.008**** | ***t*=-9.39, *P*=0.00071***** |
|  |  | *p adjusted* | 0.558 | 0.558 | **0.024*** | **0.00284**** |

**Table S10.** **Dose- and time-dependent impact of MTALTND4 on HeLa cells proliferation and viability.** HeLa (*n*=3-5, depending on the parameter). Factors: *'cell type'* ('hela' = HeLa); *'ID'* (paired sample identificator); *'treatment'* (*'ctrl'* = control, *'0.1 µM'*, *'10 µM'*, *'30 µM'* = different peptide concentration); *'time'* (*'24h'*, *'48h'*, *'72h'* = different incubation time). Parameters: *'proliferation'*: cell number (Mx cells); 'viability': amount of living cells (%). The impact of peptide addition was inferred by means of a linear mixed model (LMM), implemented for each parameter and each time-point separately. Fixed effect = *'treatment'* (4 levels). Random effect = *'ID'*. *':treatment'* = main effect of factor treatment. Simple main effects were assessed through a *post hoc* pairwise comparison. **·**0.05 > *p* ≤ 0.09; ******p* ≤ 0.05; *******p* ≤ 0.01; ********p* ≤ 0.001. Data are reported separately and also as mean ± standard error of the mean (sem). Data refer to figure 3*M,N*.

| **cell type** | **ID** | **treatment** | **time** | **proliferation (Mx cells)** | **viability (%)** |
| --- | --- | --- | --- | --- | --- |
| *hela* | *1hl* | *ctrl* | 24h | 27500 | 83.33 |
| *hela* | *2hl* | *ctrl* | 24h | 19500 | 85.71 |
| *hela* | *3hl* | *ctrl* | 24h | 45500 | 93.33 |
| *hela* | *4hl* | *ctrl* | 24h | 34500 | *na* |
| *hela* | *5hl* | *ctrl* | 24h | 31000 | *na* |
| *hela* | *1hl* | *ctrl* | 48h | 176000 | 100.00 |
| *hela* | *2hl* | *ctrl* | 48h | 130000 | 100.00 |
| *hela* | *3hl* | *ctrl* | 48h | 115000 | 95.83 |
| *hela* | *4hl* | *ctrl* | 48h | 109000 | *na* |
| *hela* | *5hl* | *ctrl* | 48h | 106000 | *na* |
| *hela* | *1hl* | *ctrl* | 72h | 495000 | 90.00 |
| *hela* | *2hl* | *ctrl* | 72h | 172500 | 84.15 |
| *hela* | *3hl* | ctrl | 72h | 310000 | 97.64 |
| *hela* | *4hl* | ctrl | 72h | 153500 | *na* |
| *hela* | *5hl* | ctrl | 72h | 280000 | *na* |
| *hela* | *1hl* | 0,1µM | 24h | 27500 | 71.43 |
| *hela* | *2hl* | 0,1µM | 24h | 61750 | 95.00 |
| *hela* | *3hl* | 0,1µM | 24h | 19500 | 100.00 |
| *hela* | *4hl* | 0,1µM | 24h | 32500 | *na* |
| *hela* | *5hl* | 0,1µM | 24h | 30000 | *na* |
| *hela* | *1hl* | 0,1µM | 48h | 143000 | 89.66 |
| *hela* | *2hl* | 0,1µM | 48h | 87500 | 100.00 |
| *hela* | *3hl* | 0,1µM | 48h | 62500 | 96.15 |
| *hela* | *4hl* | 0,1µM | 48h | 106500 | *na* |
| *hela* | *5hl* | 0,1µM | 48h | 94000 | *na* |
| *hela* | *1hl* | 0,1µM | 72h | 412500 | 92.78 |
| *hela* | *2hl* | 0,1µM | 72h | 252500 | 99.02 |

| *hela* | *3hl* | 0,1µM | 72h | 362500 | 98.64 |
| --- | --- | --- | --- | --- | --- |
| *hela* | *4hl* | 0,1µM | 72h | 210500 | *na* |
| *hela* | *5hl* | 0,1µM | 72h | 288500 | *na* |
| *hela* | *1hl* | 10µM | 24h | 60500 | 100.00 |
| *hela* | *2hl* | 10µM | 24h | 42250 | 92.86 |
| *hela* | *3hl* | 10µM | 24h | 16500 | 100.00 |
| *hela* | *4hl* | 10µM | 24h | 43000 | *na* |
| *hela* | *5hl* | 10µM | 24h | 38500 | *na* |
| *hela* | *1hl* | 10µM | 48h | 143000 | 89.66 |
| *hela* | *2hl* | 10µM | 48h | 82500 | 94.29 |
| *hela* | *3hl* | 10µM | 48h | 55000 | 91.67 |
| *hela* | *4hl* | 10µM | 48h | 61500 | *na* |
| *hela* | *5hl* | 10µM | 48h | 70500 | *na* |
| *hela* | *1hl* | 10µM | 72h | 291500 | 97.40 |
| *hela* | *2hl* | 10µM | 72h | 280000 | 97.39 |
| *hela* | *3hl* | 10µM | 72h | 260000 | 98.11 |
| *hela* | *4hl* | 10µM | 72h | 226500 | *na* |
| *hela* | *5hl* | 10µM | 72h | 214000 | *na* |
| *hela* | *1hl* | 30µM | 24h | 27500 | 83.33 |
| *hela* | *2hl* | 30µM | 24h | 39000 | 92.31 |
| *hela* | *3hl* | 30µM | 24h | 9750 | 42.86 |
| *hela* | *4hl* | 30µM | 24h | 43500 | *na* |
| *hela* | *5hl* | 30µM | 24h | 21500 | *na* |
| *hela* | *1hl* | 30µM | 48h | 77000 | 93.33 |
| *hela* | *2hl* | 30µM | 48h | 45000 | 90.00 |
| *hela* | *3hl* | 30µM | 48h | 47500 | 82.61 |
| *hela* | *4hl* | 30µM | 48h | 77500 | *na* |
| *hela* | *5hl* | 30µM | 48h | 62000 | *na* |
| *hela* | *1hl* | 30µM | 72h | 346500 | 94.64 |
| *hela* | *2hl* | 30µM | 72h | 75000 | 85.71 |
| *hela* | *3hl* | 30µM | 72h | 87500 | 89.74 |
| *hela* | *4hl* | 30µM | 72h | 288000 | *na* |
| *hela* | *5hl* | 30µM | 72h | 101000 | *na* |
|  |  | *ctrl* |  | 31600 ± 4273.172 | 87.457 ± 3.016 |
|  |  | *0,1*µ*M* |  | 34250 ± 7212.836 | 88.81 ± 8.809 |
|  |  | *10µM*  *30µM* | 24h | 40150 ± 7031.003  28250 ± 6068.361 | 97.62 ± 2.38  72.833 ± 15.209 |

|  | *LMM* | *transformation* |  | *none* | *none* |
| --- | --- | --- | --- | --- | --- |
|  |  | *:treatment* |  | *F*3,15=0.82, *P*=0.5025 | *F*3,12=1.95, *P*=0.1745 |
|  |  | *ctrl* |  | 127200 ± 12881.77 | 98.61 ± 1.39 |
|  |  | *0,1*µ*M* |  | 98700 ± 13193.37 | 95.27 ± 3.017 |
|  |  | *10µM* |  | 82500 ± 15813.76 | 91.873 ± 1.34 |
|  |  | *30µM* |  | 61800 ± 6943.702 | 88.647 ± 3.168 |
|  |  | *transformation* |  | *none* | *none* |
| *Stats* |  | *:treatment* | 48h | ***F*3,15=18.83, *P*=2.39e-05***** | ***F*3,9=5.22, *P*=0.023*** |
|  |  | *0,1*µ*M - ctrl* |  | **0.004587**** | 0.60412 |
|  |  | *10µM - ctrl* |  | **3.34e-06***** | 0.05639∙ |
|  |  | *30µM - ctrl* |  | **2.12e-12***** | **0.00107**** |
|  |  | *10µM - 0,1µM* |  | 0.071642∙ | 0.60412 |
|  |  | *30µM - 0,1µM* |  | **0.000163***** | 0.05639∙ |
|  |  | *30µM - 10µM* |  | **0.042696*** | 0.60412 |
|  |  | *ctrl* |  | 282200 ± 61097.995 | 90.597 ± 3.906 |
|  |  | *0,1*µ*M* |  | 305300 ± 36620.213 | 96.813 ± 2.02 |
|  |  | *10µM*  *30µM* | 72h | 254400 ± 14956.102  179600 ± 57099.781 | 97.633 ± 0.238  90.03 ± 2.582 |
|  |  | *transformation* |  | *none* | *none* |
|  |  | *:treatment* |  | *F*3,15=2.86, *P*=0.071∙ | *F*3,9=3.72, *P*=0.054∙ |
